# Inverse directions of association of higher physical activity and higher insulin resistance with human skeletal muscle cell typae abundance and fiber-type-level gene expression

**DOI:** 10.1101/2025.10.27.683567

**Authors:** Dan L. Ciotlos, Sarah C. Hanks, Arushi Varshney, Michael R. Erdos, Nandini Manickam, Heather M. Stringham, Peter Orchard, Erin M. Hill-Burns, Narisu Narisu, Lori L. Bonnycastle, Michael D. Sweeney, Jouko Saramies, Markku Laakso, Jaakko Tuomilehto, Timo A. Lakka, Karen L. Mohlke, Michael Boehnke, Francis S. Collins, Heikki A. Koistinen, Stephen C.J. Parker, Laura J. Scott

**Affiliations:** Department of Biostatistics and Center for Statistical Genetics, University of Michigan, Ann Arbor, MI 48109, USA; Gilbert S. Omenn Department of Computational Medicine and Bioinformatics, University of Michigan, Ann Arbor, MI 48109, USA; Center for Precision Health Research, National Human Genome Research Institute, National Institutes of Health, Bethesda, MD 20892, USA; South Karelia Social and Health Care District, Wellbeing Services County of South Karelia, South Karelia, Finland; Institute of Clinical Medicine, Internal Medicine, Kuopio University Hospital and University of Eastern Finland, Kuopio, Finland; Department of Public Health, University of Helsinki, Helsinki, Finland; Institute of Biomedicine, School of Medicine, University of Eastern Finland, Kuopio, Finland; Kuopio Research Institute of Exercise Medicine, Kuopio, Finland; Department of Clinical Physiology and Nuclear Medicine, Kuopio University Hospital, Kuopio, Finland; Department of Genetics, University of North Carolina, Chapel Hill, NC 27599, USA; University of Helsinki, Faculty of Medicine, Research Programs Unit, Clinical and Molecular Metabolism (CAMM), Helsinki, Finland; Department of Medicine, University of Helsinki and Helsinki University Hospital, Helsinki, Finland; Minerva Foundation Institute for Medical Research, Helsinki, Finland; Department of Human Genetics, University of Michigan, Ann Arbor, MI 48109, USA

**Keywords:** gene expression, chromatin accessibility, physical activity, insulin resistance, Type 2 diabetes, single nucleus, skeletal muscle, snRNA-seq, snATAC-seq

## Abstract

To investigate the interplay between physical activity and cardiometabolic traits in human skeletal muscle, we characterized gene expression and chromatin accessibility across skeletal muscle cell types in 263 Finnish individuals from the FUSION Tissue Biopsy Study. We analyzed skeletal muscle single-nucleus RNA-seq data (168,309 nuclei, 23,849 genes), ATAC-seq data (242,069 nuclei, 927,588 peaks), and bulk RNA-seq data (22,309 genes). Lower insulin resistance (HOMA-IR) and higher total physical activity were both associated with higher proportions of Type 1 nuclei and lower proportions of Type 2x nuclei. We identified cell-type-level and tissue-level gene expression–trait and gene set–trait associations for cardiometabolic and physical activity traits, and a smaller proportion of cell-type-level chromatin accessibility–trait associations. Traits typically associated with better health—lower trait values of cardiometabolic traits (BMI, HOMA-IR, normal glucose tolerance vs. type 2 diabetes, 2-hour plasma glucose) and higher physical activity levels (total and vigorous)—were associated with higher expression of energy metabolism genes and lower expression of signaling pathway genes across muscle fiber types, total pseudobulk, and to some extent in bulk tissue. For HOMA-IR and physical activity, these directions of association remained when adjusting for both traits in the same model, indicating apparently independent associations in the same pathways.

## Introduction

Low physical activity levels and high insulin resistance are risk factors for type 2 diabetes (T2D); both risk factors influence and are influenced by the physiological state of skeletal muscle.^1–4^ A significant proportion of glucose disposal after an oral glucose load occurs in skeletal muscle, and insulin-stimulated glucose uptake is reduced in individuals with insulin resistance or a sedentary lifestyle.^5,6^ Regular physical activity improves skeletal muscle insulin sensitivity and reduces the risk of T2D.^7,8^ These physiological adaptations are mediated by skeletal muscle cell type abundance and gene expression within cells.

Skeletal muscle fiber type abundance varies by physical activity, degree of insulin resistance, and T2D status. In interventional studies, strength training is associated with a transition from fast-twitch, oxidative-glycolytic Type 2a and fast-twitch, glycolytic Type 2x muscle fibers to slow-twitch, oxidative Type 1 muscle fibers, although changes in fiber type abundance vary depending on the type of strength or endurance exercise performed.^9–11^ Normal glucose tolerance (NGT), as compared to T2D, is associated with higher proportions of Type 1 muscle fibers and lower proportions of Type 2x muscle fibers.^12–15^ These studies quantify cell type proportions by histochemical staining or fiber type-specific myosin heavy chain isoform expression, but provide no information on gene expression.

In analyses of bulk skeletal muscle, gene expression levels differ by physical activity, insulin resistance, and T2D status.^10,16–20^ Expression of mitochondrial function genes is widely reported to vary by T2D status. Individuals with NGT, as compared to those with T2D, show lower expression of cellular respiration genes.^18^ Participation in aerobic training is associated with higher expression of cellular respiration genes relative to an untrained baseline.^10^ Other biological processes, including protein modification and cell signaling, are also associated across these physiological states. Bulk muscle RNA-seq data measures exonic reads from both cytoplasmic and nuclear mRNA at the whole-tissue level^21^. In contrast, single-nucleus RNA-seq (snRNA-seq) cell-type-level pseudobulk and total pseudobulk (all cell types combined) data capture exonic and intronic nuclear mRNA.^22,23^ These data types provide complementary perspectives on the relationship between gene expression and physiological traits; however, bulk RNA-seq studies may be confounded by individual differences in cell type abundance and cannot identify cell-type-specific gene expression differences.^24^

Single-nucleus studies enable the quantification of cell type abundance, cell-type-level gene expression, and cell-type-level chromatin accessibility. A small study in four humans found both shared and skeletal muscle fiber-type-specific differentially expressed genes and accessible chromatin regions in response to endurance exercise.^25^ Another study in 38 individuals identified differences in the transcriptional profiles of skeletal muscle fiber myonuclei, particularly Type 2a myonuclei, between individuals with and without T2D after 2-week high-intensity interval training.^26^ However, little is known about how insulin resistance affects gene expression or chromatin accessibility within specific skeletal muscle fiber types. Most bulk tissue and single-nucleus studies have focused on gene expression–trait associations for either physical activity or cardiometabolic traits, but few have investigated the association of both traits in the same individuals.

To investigate the interplay between physical activity and cardiometabolic traits in human skeletal muscle, we characterized gene expression and chromatin accessibility across skeletal muscle cell types in 263 Finnish individuals from the Finland-United States Investigation of NIDDM Genetics (FUSION) Tissue Biopsy Study. We analyzed skeletal muscle snRNA-seq data (168,309 nuclei, 23,849 genes), single-nucleus ATAC-seq (snATAC) data (242,069 nuclei, 927,588 peaks), and bulk RNA-seq data (22,309 genes). We tested for associations of physical activity and cardiometabolic traits, including body mass index (BMI), insulin resistance (Homeostatic Model Assessment of Insulin Resistance, HOMA-IR), and drug-naive T2D status (T2D vs. NGT), with cell type abundance, cell-type-level and bulk tissue gene expression, and cell-type-level chromatin accessibility. We identified opposite and apparently independent directions of association of higher physical activity and higher insulin resistance with skeletal muscle cell type abundance and fiber-type-level gene expression.

## Results

### Vastus lateralis muscle biopsies

Our goal was to understand the pathways through which cardiometabolic and physical activity traits influence or are influenced by skeletal muscle gene expression or chromatin accessibility. We studied vastus lateralis biopsies from 263 Finnish individuals as part of the FUSION Tissue Biopsy Study.^18^ Insulin resistance was measured using HOMA-IR.^27^ Leisure time physical activity was assessed using a 12-month activity questionnaire.^28–30^ For analyses of cell type abundance, gene expression, and chromatin accessibility, we used a false discovery rate (FDR) threshold of <5% to define statistical significance.

Participants had a mean age of 60.3 ± 7.6 years, a mean BMI of 27.7 ± 4.2 kg/m², and 43% were female (Table S1). 26.6% of participants had newly-diagnosed, drug-naive T2D, 12.9% had impaired fasting plasma glucose (IFG), 24.7% had impaired glucose tolerance (IGT), 35.7% had NGT.^31^ Individuals with T2D tended to have higher insulin resistance (HOMA-IR) than those with NGT (Figure S1), although some individuals with NGT had clinically high insulin resistance (HOMA-IR>2.9 as defined by da Silva et al).^32^ Total and vigorous physical activity capture different aspects of energy expenditure, as some individuals reported high amounts of total physical activity despite engaging in little or no vigorous activity (Figure S2).

Cardiometabolic traits, including BMI, HOMA-IR, T2D status (T2D vs. NGT), and 2-hour plasma glucose, were positively correlated with each other (⍴ range: 0.32 to 0.73); total and vigorous physical activity were positively correlated with each other (⍴=0.58, p-value=1.5x10⁻²⁴). 2-hour fasting glucose was negatively correlated with vigorous physical activity (⍴=-0.20, p-value=1.3x10⁻³) (Figures 1A and S2).

**Figure 1.**
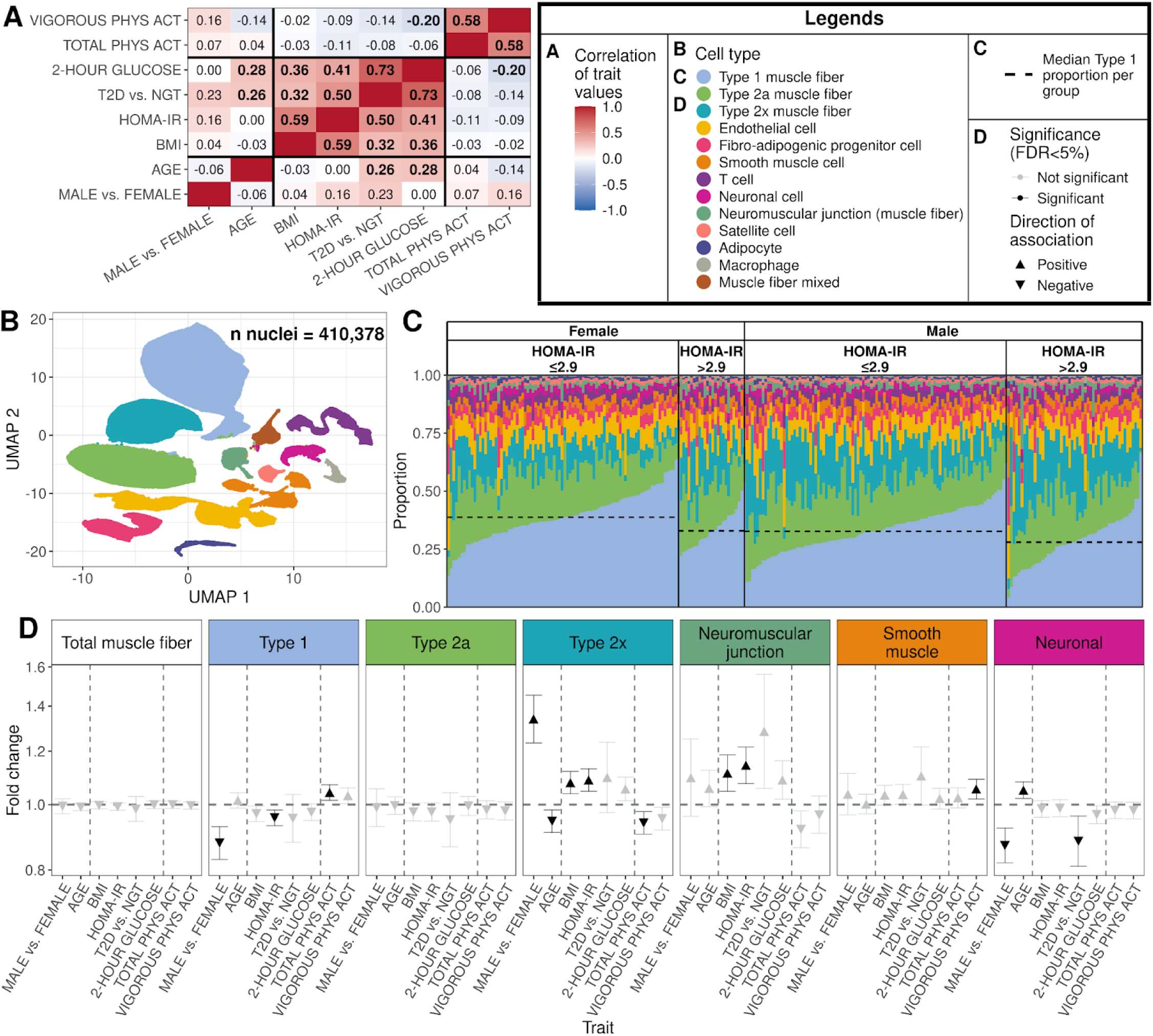
Correlation of trait values and differences in cell type abundance in human skeletal muscle. (A) Heatmap of Spearman correlations of trait values in 263 individuals. Sample characteristics for each trait can be found in Table S1. Male vs. female and T2D vs. NGT variables were coded as binary 0/1 variables. Physical activity is abbreviated as PHYS ACT. Numbers in bold represent correlations significantly different than 0 tested at a Bonferroni-adjusted significance level of 0.05/28. (B) UMAP projection of 410,378 RNA and ATAC nuclei across 263 individuals with both snATAC and snRNA nuclei. Each point is a nucleus. (C) Cell type proportions for 263 individuals with both snATAC and snRNA nuclei separated by sex and insulin resistance level (HOMA-IR≤2.9 or >2.9). (D) Fold change and 95% confidence intervals comparing the number of snRNA and snATAC nuclei by trait, adjusting for the number of total nuclei (FDR<5% across all tests). Sample sizes for each trait can be found in the methods.

### Skeletal muscle cell type abundance

We analyzed 12 cell types, identified in jointly-clustered skeletal muscle snRNA-seq (168,309 nuclei) and snATAC-seq (242,069 nuclei) nuclei (Figure 1B).^33^ We visualized cell type abundance across males and females and stratified by HOMA-IR to examine across-person variability. On average, muscle fiber types made up 70% of the snRNA and snATAC nuclei (34% Type 1, 20% Type 2a, and 16% Type 2x) (Table S2). In barplots of cell type proportions, individuals with clinically high insulin resistance (HOMA-IR>2.9) appeared to have a lower median proportion of Type 1 muscle fiber nuclei than individuals without insulin resistance (HOMA-IR ≤ 2.9) in both sexes (Figure 1C).

We used negative binomial models to test for associations between cell type abundance of snRNA and snATAC nuclei, and cardiometabolic and physical activity traits (Figure 1D and Table S3). Throughout this paper, we include results for sex for comparison to other traits (see Hanks et al. for a more extensive analysis of the association of sex in skeletal muscle).^34^ Males had a higher proportion of Type 2x and a lower proportion of Type 1 and neuronal cell nuclei than females. Higher BMI was associated with higher proportions of Type 2x and neuromuscular junction nuclei. Lower HOMA-IR and higher total physical activity—traits typically associated with better health—were both associated with higher proportions of Type 1 nuclei and lower proportions of Type 2x nuclei. Higher HOMA-IR was associated with higher proportions of neuromuscular junction nuclei, while higher vigorous physical activity was associated with higher proportions of smooth muscle cell nuclei. For all traits and cell types, the directions of statistically significant cell type abundance–trait associations were consistent when analyzing nuclei from snATAC-seq or snRNA-seq separately (Figure S3).

The opposite directions of association with cell type abundance for higher HOMA-IR and higher physical activity could reflect shared underlying biology or confounding between these two traits. To help distinguish between these possibilities, we fitted new regression models that included both HOMA-IR and total physical activity as covariates. We found minimal to no change in fold change estimates for cell type abundance-trait associations for either HOMA-IR or total physical activity when adjusting for the other trait (Figure S3), indicating that insulin resistance and total physical activity were potentially independently associated with the abundance of Type 1 and Type 2x muscle fiber nuclei.

### Gene expression–trait associations in skeletal muscle cell types, total pseudobulk, and bulk tissue

To explore how cardiometabolic and physical activity traits were associated with skeletal muscle gene expression, we tested for associations between physiological traits and gene expression using a negative binomial model across three data types: (i) cell-type-level pseudobulk data in the 10 most abundant muscle cell types, (ii) total pseudobulk data, and (iii) bulk RNA-seq data. For total pseudobulk and bulk muscle RNA-seq data, we adjusted for each sample’s observed single-nucleus cell type proportions to reduce confounding from inter-individual differences in cell type abundance. Across muscle fiber types, total pseudobulk, and bulk tissue, we found the highest number and percentage of associated genes for sex (male vs. female, 1974–7192 genes, 11.3–32.2% tested genes), followed by BMI (257–1377 genes, 1.5–6.2%), HOMA-IR (260–1955 genes, 1.5–8.8%), age (53–1275 genes, 0.2–5.7%), and vigorous physical activity (22–403 genes, 0.1–1.7%) (Figure 2A and Table S4).

**Figure 2.**
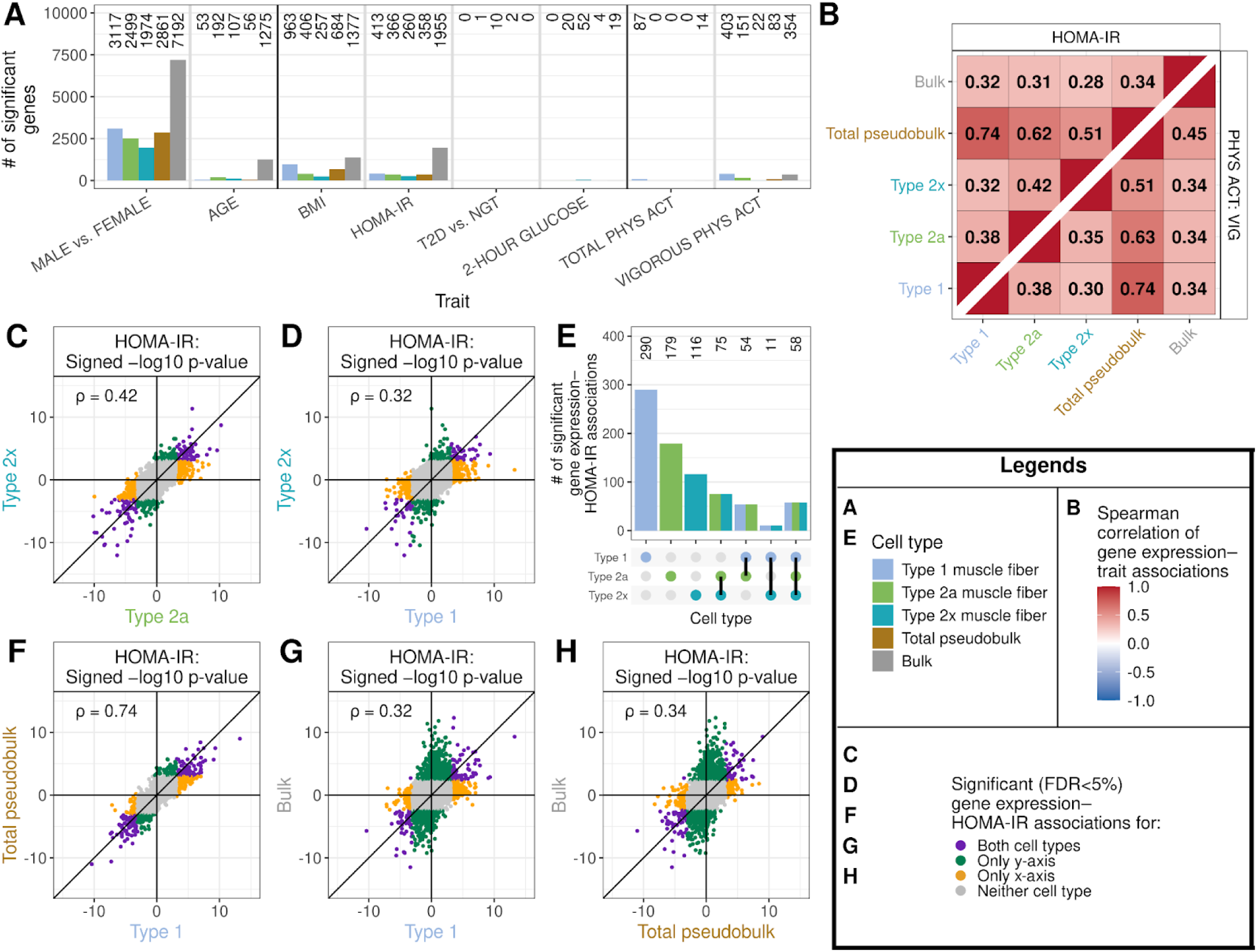
Gene expression–trait associations across cell types. (A) Number of significant (FDR<5%) genes for each trait across muscle fiber types, total pseudobulk, and bulk tissue. (B) Heatmap of Spearman correlations of signed (by beta coefficient) −log10 p-value of gene expression–trait associations between cell types for (top left) HOMA-IR and (bottom right) vigorous physical activity. Numbers in bold represent correlations significantly different than 0 tested at a Bonferroni-adjusted significance level of 0.05/132. (C, D, F, G, H) Scatter plots of signed −log10 p-values of gene expression–HOMA-IR associations in (C) Type 2a vs. 2x, (D) Type 1 vs. 2x, (F) Type 1 vs. total pseudobulk, (G) Type 1 vs. bulk tissue, and (H) total pseudobulk vs. bulk tissue. Each point is a gene. (E) Upset plot of the number of significant gene expression–HOMA-IR associations in the muscle fiber types. Number of genes tested and sample sizes for each trait and cell type combination can be found in Tables S15 and S16.

For each trait, we typically found a larger number of associated genes in bulk tissue than in total pseudobulk, and in Type 1 fiber than in Type 2a or 2x fibers; we found few associations in the remaining cell types (Figure 2A and Tables S4 and S5). As the number of total unique molecular identifiers (UMIs) summed across all genes and individuals vary by cell type, we downsampled Type 1 muscle fibers (to match the total UMIs in each cell type, see Methods) to investigate whether differences in the number of gene expression–trait associations between Type 1 muscle fiber and other cell types were due to differences in power (Figure S4). Compared to the downsampled Type 1 fiber data, in Type 2a and 2x fibers we found more differentially-associated genes for age, similar numbers for HOMA-IR, and fewer differentially-associated genes for BMI and vigorous physical activity (Figure S5 and Table S6). In other cell types, we found fewer differentially-associated genes than in downsampled Type 1 fibers. Overall, age and HOMA-IR appear to have a stronger influence on gene expression in Type 2a and 2x fibers relative to Type 1 fibers.

### Comparison of gene expression–trait associations between skeletal muscle cell types, total pseudobulk, and bulk tissue

We next assessed if, for each trait, the same or different genes were associated with that trait across cell types, total pseudobulk, and bulk tissue. Identifying shared or distinct gene expression–trait associations could reveal cell-type-specific differential expression patterns that may be missed when gene expression is aggregated across cell types. We focused first on associations with HOMA-IR as this trait yielded more associated genes than T2D vs. NGT status (which includes a smaller proportion of the sample size), and because skeletal muscle plays a central role in the body’s response to insulin.^6^

Starting with the muscle fiber types, we observed strong, positive correlations of the signed (by beta coefficient) −log10 p-values of gene expression–HOMA-IR associations between all muscle fibers, with the strongest correlation between Type 2a and 2x (⍴=0.42, p-value<10⁻²⁰⁰) (Figures 2B, 2C, 2D, and S6). We used an Upset plot to visualize the overlap of significantly-associated genes with HOMA-IR in the three muscle fiber types (Figure 2E). A large proportion of genes were uniquely, significantly associated in a single fiber type (Type 1: 70% of all Type 1 associated genes, Type 2a: 49%, Type 2x: 45%). Type 2a and Type 2x shared the highest number of significantly associated genes (n=75), followed by Type 1 and Type 2a (n=54), and Type 1 and Type 2x (n=11) (Figure 2E). 58 genes were significantly associated across all three muscle fiber types. Genes that were differentially expressed in more than one fiber type always had concordant directions of association (Table S7).

Compared to the correlations between muscle fiber types, we found substantially weaker correlations of gene expression–HOMA-IR associations between non-muscle fiber cell types and any other cell type (all |⍴|<0.08) (Figure S7); these weaker correlations may reflect differences in statistical power to detect trait associations in the less abundant non-muscle fiber types. However, we found no diminution of correlations of gene expression–HOMA-IR results for Type 2a or 2x with those of Type 1 fibers when the latter was downsampled to match UMI levels in the less abundant cell types (see Methods) (Figure S8). This suggests that differences in association patterns between muscle fiber and non-muscle fiber cell types are not solely due to differences in power, but likely reflect true cell-type-level biological differences.

We next compared cell-type-level gene expression–HOMA-IR association results to those observed in total pseudobulk and bulk tissue. Association results from the total pseudobulk dataset were most strongly correlated with Type 1 muscle fiber (⍴=0.74, p-value<10⁻²⁰⁰), followed by the other muscle fiber types (Figures 2B and 2F). This likely reflects the substantial contribution of muscle fibers, in particular Type 1 fiber, to total pseudobulk data. Similarly, bulk tissue was most strongly correlated with Type 1 muscle fiber (⍴=0.32, p-value<10⁻²⁰⁰) (Figures 2B and 2G). As expected, we found stronger correlations between cell-type-level results with total pseudobulk than with bulk tissue, potentially because total pseudobulk aggregates the same nuclei profiled at the cell type level and captures only nuclear mRNA.

We further compared gene expression–HOMA-IR associations between total pseudobulk and bulk tissue. Differences between the two data types may arise because nuclear and cytoplasmic mRNA are regulated differently or because the proportions of cell types captured in the single-nucleus experiment differ from those in the intact tissue. Of 115 significant gene expression–HOMA-IR associations in both total pseudobulk and bulk tissue, 114 (99%) had the same direction of association (Table S7). However, of the total 1569 significantly-associated genes in bulk tissue also tested in total pseudobulk, only 1222 genes (78%, including non-significant total pseudobulk genes) had a concordant direction of association in total pseudobulk. Overall, gene expression–HOMA-IR associations were significantly, positively correlated between total pseudobulk and bulk tissue (⍴=0.34, p-value<10⁻²⁰⁰) (Figure 2H); when only considering genes with high mean gene counts (counts>100) in both data types, we observed a stronger correlation (⍴: 0.41-0.68, Figure S9). This suggests a generally consistent pattern of gene expression–HOMA-IR associations between total pseudobulk and bulk tissue, particularly for genes highly expressed in both datasets. Likewise, we observed similar patterns of gene expression–vigorous physical activity associations between cell types, total pseudobulk, and bulk tissue (Figures 2B, S7, S8, S10, and Table S7).

### Opposite directions of association with gene expression for higher HOMA-IR and higher physical activity

Previous bulk skeletal muscle tissue studies have reported that insulin resistance and physical activity were associated with sets of genes involved in energy metabolism and signaling pathways;^10,16–20^ however, the extent to which these associations are identified in the same individuals and in specific muscle cell types remains unclear. To investigate this, we focused on vigorous physical activity as our trait of interest as a larger proportion of genes were differentially expressed for vigorous physical activity than total physical activity. To compare the directions of association for gene expression–trait associations between HOMA-IR and vigorous physical activity, we plotted the signed (by beta coefficient) −log10 p-values of associations for each trait in the muscle fiber types, total pseudobulk, and bulk tissue. We found that gene expression association results between vigorous physical activity and HOMA-IR were significantly negatively correlated in all muscle fibers, total pseudobulk, and bulk tissue (range ⍴: -0.27 to -0.37) (Figures 3A, 3B, and S11). Nearly all genes that are differentially expressed for both traits showed opposite directions of association for higher HOMA-IR and higher vigorous physical activity for each muscle fiber type (100% opposite association, n gene range: 5-56), total pseudobulk (100%, n=14), and bulk tissue (94%, n=69) (Table S8).

**Figure 3.**
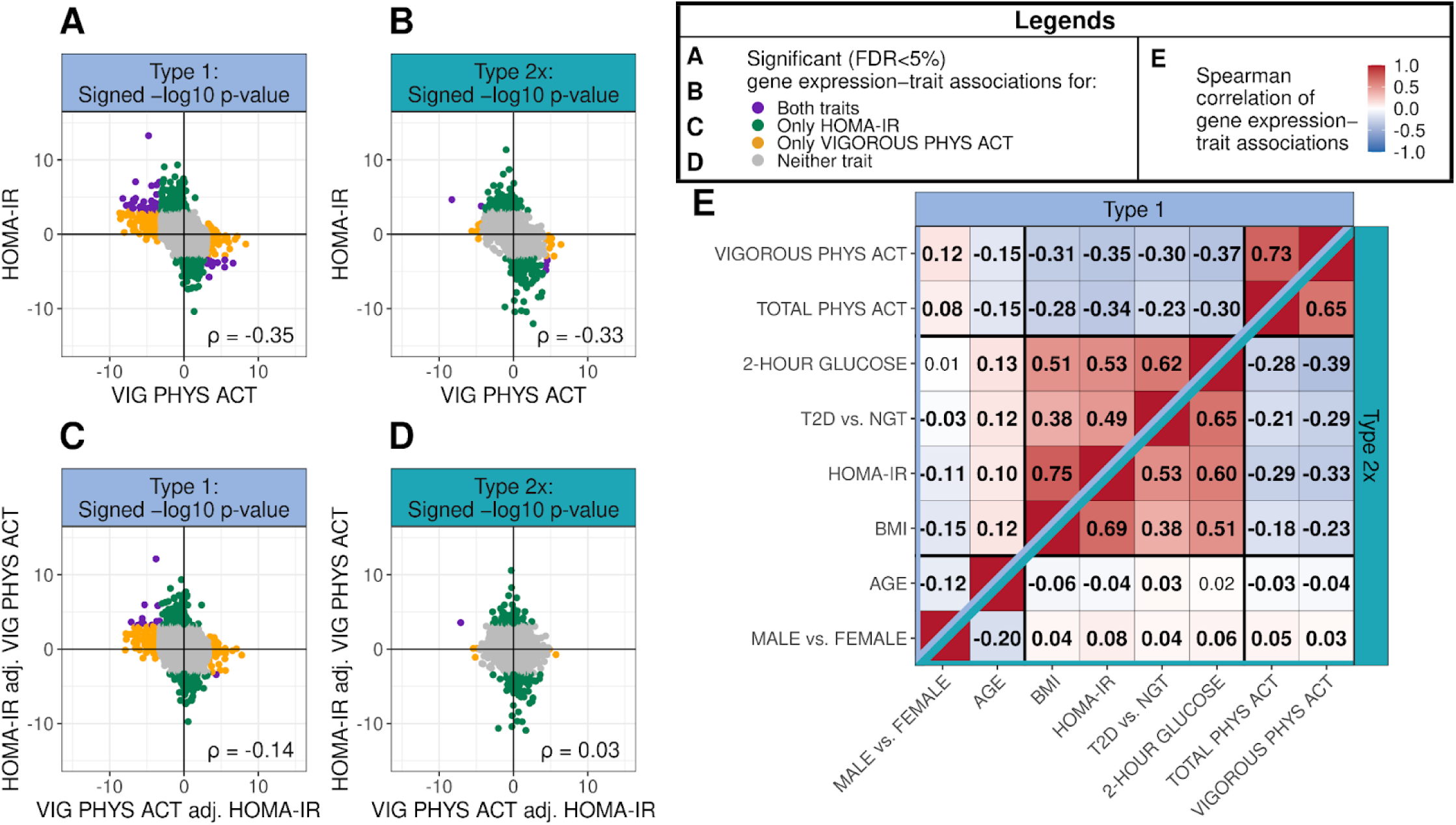
Gene expression–trait associations across traits. (A, B) Scatter plots of signed −log10 p-values of gene expression–trait associations for HOMA-IR vs. vigorous physical activity (VIG PHYS ACT) when only adjusting for base covariates in (A) Type 1 and (B) Type 2x. Each point is a gene. (C, D) Scatter plots of signed −log10 p-values of gene expression–trait associations for HOMA-IR vs. vigorous physical activity (VIG PHYS ACT) when jointly adjusting (adj.) for both traits in addition to base covariates in (C) Type 1 and (D) Type 2x. Each point is a gene. (E) Heatmap of Spearman correlations of signed (by beta coefficient) −log10 p-values of gene expression–trait associations between traits in (top left) Type 1 and (bottom right) Type 2x from models only adjusting for base covariates. Numbers in bold represent correlations significantly different than 0 tested at a Bonferroni-adjusted significance level of 0.05/224. Number of genes tested and sample sizes for each trait and cell type combination can be found in Tables S15 and S16.

The opposite directions of association for gene expression results for higher HOMA-IR and higher vigorous physical activity in muscle fiber types could reflect shared underlying biology or confounding between physical activity and HOMA-IR. To evaluate the potential independence of these associations, we included both HOMA-IR and vigorous physical activity in the same regression model. Overall, the correlations of jointly-adjusted HOMA-IR and vigorous physical activity gene expression–trait associations were attenuated relative to the unadjusted analyses, but remained negatively correlated in Type 1 (⍴=-0.14, p-value=1.6x10⁻¹⁰³), Type 2a (⍴=-0.06, p-value=7.6x10⁻¹⁷), total pseudobulk (⍴=-0.12, p-value=8.8x10⁻⁶⁴), and bulk tissue (⍴=-0.15, p-value=2.4x10⁻¹¹²), but not in Type 2x (⍴=0.03, p-value=4.1x10⁻⁵) fibers (Figures 3C, 3D, S11, S12, and S13). This suggests potentially independent associations with lower insulin resistance and higher vigorous physical activity for a subset of genes.

We hypothesized that other cardiometabolic traits would mirror the opposite directions of gene expression association results observed for HOMA-IR and physical activity. Consistent with our expectation, we found significant, negative correlations for gene expression–trait associations between cardiometabolic traits and physical activity traits in the muscle fiber types, total pseudobulk, and bulk tissue (⍴ range: -0.13 to -0.42). Similar, though weaker, negative correlations were observed in other non-muscle fiber cell types with the exception of BMI and physical activity which did not show a consistent pattern of correlation (Figures 3E and S14).

### Opposite gene set directions of association for higher HOMA-IR and higher physical activity

Differences in gene expression associated with a cardiometabolic or physical activity trait may be coordinated, with multiple genes involved in a similar function or biological process showing concordant changes in expression. We performed Gene Ontology (GO) term enrichment analysis using RNA-Enrich^35^ to identify biological processes enriched for genes with positive or negative associations with these traits across cell types, total pseudobulk, and bulk tissue. Across muscle fiber types, total pseudobulk, and bulk, we found that bulk tissue had the greatest number of associated gene sets for every trait except for HOMA-IR, which was consistent with finding the most differentially expressed genes (DEGs) in bulk tissue (Figure 4A). Physical activity traits yielded the highest number of significant gene sets in Type 1 muscle fibers, while cardiometabolic traits yielded the most in Type 2a muscle fibers. Fewer than 35 significant gene sets were identified in any non-muscle fiber cell types for any trait (Tables S9 and S10).

**Figure 4.**
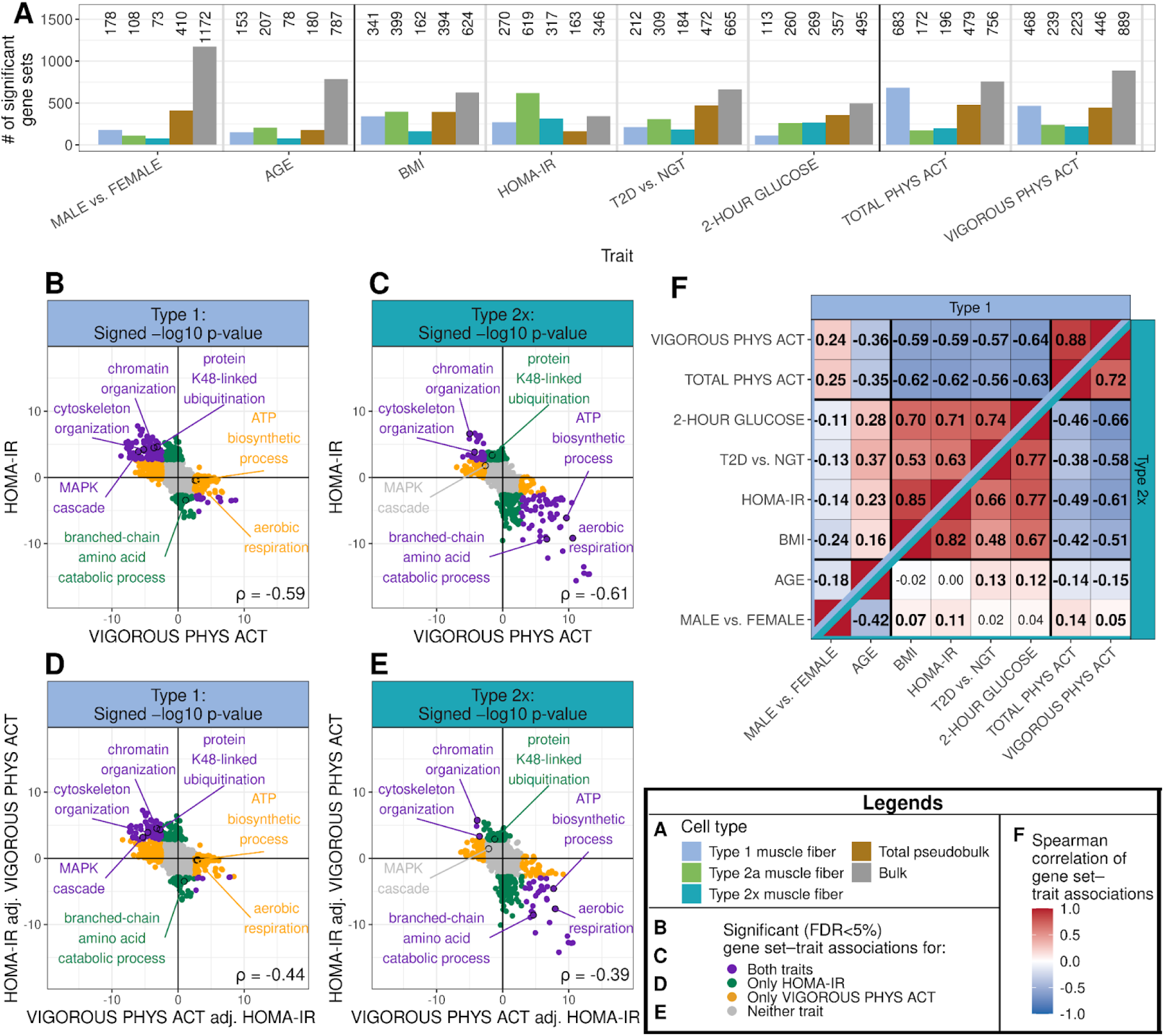
Gene set–trait associations across traits. (A) Number of significant (FDR<5%) gene sets for gene set–trait associations across muscle fiber types, total pseudobulk, and bulk. Gene sets tested are Gene Ontology biological processes. (B, C) Scatter plots of signed −log10 p-values of gene set–trait associations for HOMA-IR vs. vigorous physical activity when only adjusting for base covariates in gene expression models in (B) Type 1 and (C) Type 2x. Each point is a gene set. (D, E) Scatter plots of signed −log10 p-values of gene set–trait associations for HOMA-IR vs. vigorous physical activity when jointly adjusting (adj.) for both traits in gene expression models in (D) Type 1 and (E) Type 2x. Each point is a gene set. (F) Heatmap of Spearman correlations of signed (by beta coefficient) −log10 p-values of gene set–trait associations between traits in (top left) Type 1 and (bottom right) Type 2x from gene expression models only adjusting for base covariates. Numbers in bold represent correlations significantly different than 0 tested at a Bonferroni-adjusted significance level of 0.05/224. Sample sizes for each trait and cell type combination can be found in Table S16.

Opposite directions of association with gene expression results for higher HOMA-IR and higher physical activity in muscle fiber types suggest that insulin resistance and physical activity might influence shared biological pathways. We compared gene set results for vigorous physical activity and HOMA-IR and observed significant, negative correlations between gene set–trait association results in Type 1 and 2x muscle fibers (Figures 4B, and 4C), as well as Type 2a muscle fiber, total pseudobulk, and bulk tissue (⍴ range: -0.61 to -0.30) (Figure S15). Consistent with these correlations, nearly all significant gene sets for both traits showed opposite directions of association for higher HOMA-IR and higher vigorous physical activity in each muscle fiber type (100% opposite association, n gene set range: 98-112), total pseudobulk (100%, n=78), and bulk tissue (94%, n=131) (Table S11).

The opposite directions of association of gene set enrichment results for HOMA-IR and physical activity could reflect shared underlying biology or confounding between the traits. To evaluate the potential independence of these associations, we reran GO term enrichment analysis in each data type using the jointly-adjusted gene expression–trait association data. Overall, the jointly-adjusted HOMA-IR and vigorous physical activity gene set–trait expression associations were attenuated but still significantly negatively correlated in all muscle fiber types, total pseudobulk, and bulk tissue (⍴ range: -0.44 to -0.16) (Figures 4D, 4E, and S15). Most gene sets were significant for HOMA-IR or vigorous physical activity with and without jointly adjusting for the other trait in all muscle fiber types and bulk tissue (% range: 63-88%, Figure S16); however, in total pseudobulk, only 33% of the originally-significant gene sets were still significant for HOMA-IR after adjusting for vigorous physical activity. Overall, HOMA-IR and vigorous physical activity gene set associations were less significant when adjusting for the other trait, except for vigorous physical activity in bulk tissue (Figures S17 and S18).

In line with our earlier hypothesis, we expected that other cardiometabolic traits would mirror the opposite directions of gene set association results observed between HOMA-IR and physical activity. Consistently, when comparing traits within a cell type, we found significant, moderate, negative correlations of gene set results between cardiometabolic traits and physical activity traits in Type 1 and 2x muscle fibers (Figures 4F), as well as in Type 2a muscle fiber, total pseudobulk, and bulk tissue; and weaker negative correlations in other non-muscle fiber cell types (Figure S19).

### Higher expression of genes in energy metabolism-related gene sets associated with lower insulin resistance and higher physical activity

To better understand the cell-type-specific and trait-specific context of GO terms, we show results for seven GO terms across muscle fiber types, total pseudobulk, and bulk tissue (Figure 5A).

**Figure 5.**
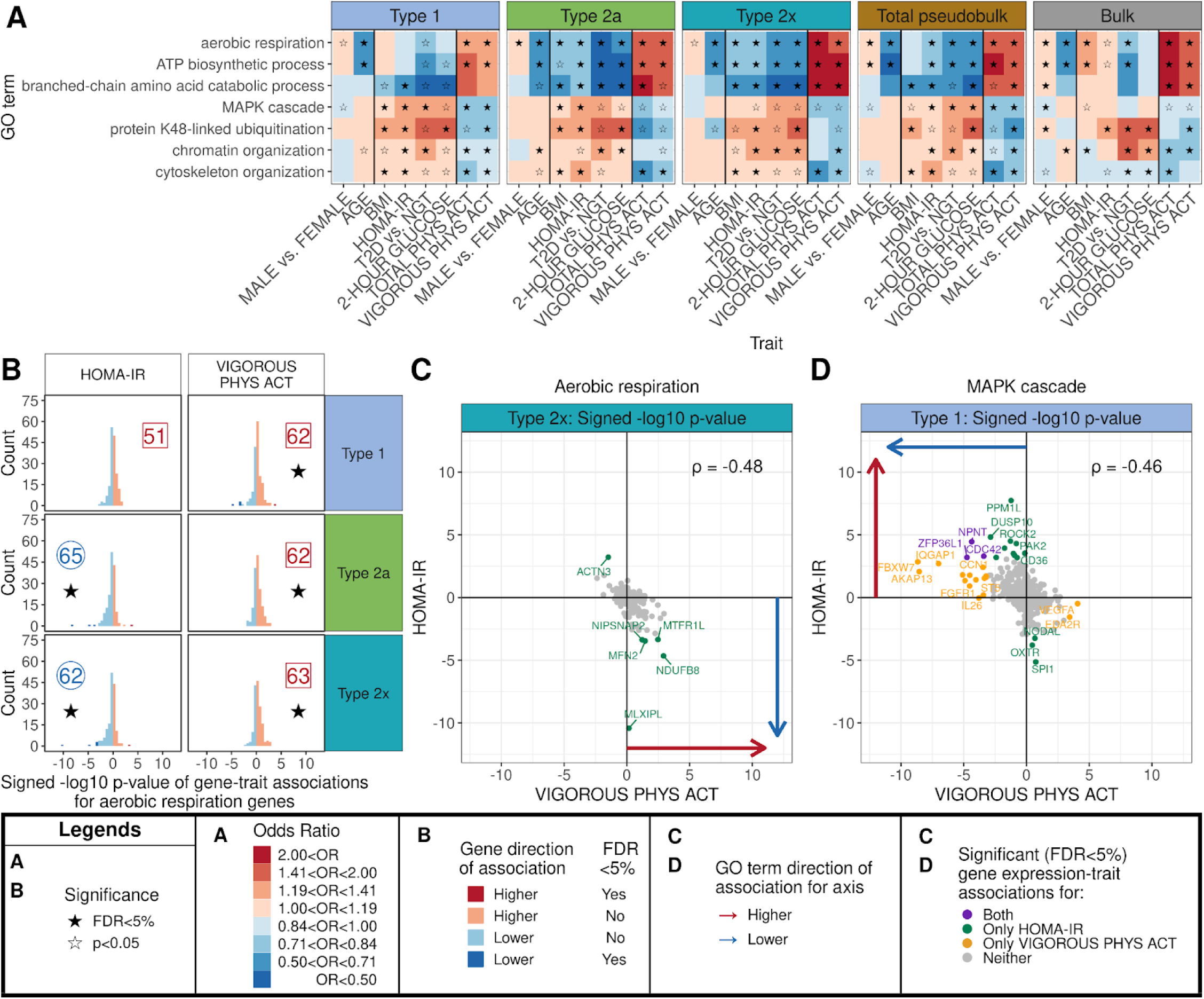
Selected significant Gene Ontology (GO) terms across data types and traits. (A) Gene set results for seven GO terms across all traits and muscle fibers, total pseudobulk, and bulk tissue. A filled-in star indicates that the gene set–trait association is significant (FDR<5%). (B) Histograms of the signed −log10 p-values of gene expression–trait associations of aerobic respiration genes with HOMA-IR and vigorous physical activity in the muscle fiber types. A star indicates that the aerobic respiration–trait association was significant for that trait in that cell type. The number indicates the percentage of genes with positive (red square) or negative (blue circle) gene expression–trait associations for the gene set. (C, D) Scatter plots of the signed −log10 p-values of gene expression–trait associations for HOMA-IR vs. vigorous physical activity for (C) aerobic respiration genes in Type 2x and (D) MAPK cascade genes in Type 1. Arrows indicate the direction of effect of the gene set–trait association from panel A for each trait. Sample sizes for each trait and cell type combination can be found in Table S16.

Complete results for significant GO terms by trait are provided in the supplementary materials (Table S10). We observed far fewer significant GO term associations in non-muscle fiber cell types.

We first examined energy metabolism-related gene sets that have been widely reported to associate with T2D and physical activity in the literature.^10,18^ Across muscle fiber types, total pseudobulk, and bulk data, we observed higher expression of ATP biosynthetic process and aerobic respiration genes associated with higher physical activity (total and vigorous), lower 2-hour plasma glucose, younger age, and NGT compared with T2D (Figure 5A). We observed similar results across multiple energy metabolism-related gene sets (Figure S20A). In Type 2a and 2x fibers and in total pseudobulk, we observed lower expression of ATP biosynthetic process and aerobic respiration genes associated with higher HOMA-IR and BMI. In contrast, in bulk data, we observed higher expression of ATP biosynthetic process and aerobic respiration genes associated with higher HOMA-IR and BMI. To assess the sensitivity of bulk data analysis to the adjustment of estimated cell type proportions, we omitted cell type proportions from the model and reran the analysis. We found that energy metabolism-related gene sets tended to change their direction of association, showing lower expression associated with higher BMI or HOMA-IR when cell type proportions were not included (Figure S21). For other traits, we found consistent aerobic respiration gene set results in the adjusted and unadjusted analyses.

To assess the consistency of the gene expression–trait associations within gene sets, we calculated the proportion of genes with the same direction of association as the gene set results.

In significantly-associated gene sets, we observed that 57–87% of genes had gene expression–trait associations in the same direction as the gene set result (Figure S22). Across the three muscle fiber types, within significantly-associated HOMA-IR and physical activity aerobic respiration gene sets, we found 62–65% of genes had directions of association consistent with the GO term direction of association, although ≤6 genes were significantly associated with either trait (Figure 5B).

The aerobic respiration gene set associations for HOMA-IR and physical activity could reflect the involvement of the same or different subsets of aerobic respiration genes. We compared aerobic respiration gene expression–trait association results for vigorous physical activity and HOMA-IR in Type 2x muscle fibers. These associations were more strongly negatively correlated (⍴=-0.48, p-value=2.6x10⁻¹⁰) (Figure 5C) than the associations across all genes (⍴=-0.33, p-value<10⁻²⁰⁰). The most significantly-associated aerobic respiration gene for HOMA-IR (*MLX1PL*) showed no association with vigorous physical activity; however, other significantly-associated aerobic respiration genes for HOMA-IR (*NDUFB8*, *MTFR1L*) were among the most strongly associated (though not significant) with vigorous physical activity (Figure 5C). To assess the effect of physical activity and insulin resistance traits on each other, we reran GO term enrichment analysis using the jointly-adjusted gene expression–trait association data. The adjusted HOMA-IR and physical activity energy metabolism-related gene set associations were attenuated compared to unadjusted gene set analysis (in all but bulk data), but had the same direction of association and remained significantly associated (Figures S20B, S23, and S24). Thus, we saw overlap in the gene sets that are affected by physical activity and HOMA-IR that was, at least in part, independent of their correlation with each other.

### Enrichment differences for signalling pathways and other gene sets by trait

While energy metabolism gene sets are some of the most widely reported insulin resistance or physical activity-associated gene sets, we found a wide variety of significantly-associated gene sets. Higher expression of branched-chain amino acid catabolic process genes was associated with lower HOMA-IR (in all muscle fiber types and total pseudobulk) and higher vigorous physical activity (Type 2x, total pseudobulk, and bulk tissue) (Figure 5A); we found similar directions of association across an expanded set of metabolic and catabolic processes gene sets relating to amino acids, lipids, nucleotides, and organic acids (Figure S25A).

The directions of association for the MAPK cascade gene set within muscle fibers and total pseudobulk were opposite those observed for the aerobic respiration and branched chain amino acid gene sets. Higher expression of MAPK cascade genes was associated with less favorable metabolic trait levels (higher BMI, higher HOMA-IR, T2D compared with NGT) and lower physical activity (total and vigorous) with statistically significant associations observed in Type 1 fibers and total pseudobulk (Figure 5A); MAPK cascade gene expression–trait associations between vigorous physical activity and HOMA-IR in Type 1 were more strongly, negatively correlated (⍴=-0.46, p-value=3.8x10⁻²⁹) than the associations across all genes (⍴=-0.35, p-value<10⁻²⁰⁰).

Three MAPK cascade genes (*NPNT*, *ZFP36L1*, *CDC42*) were significantly associated with higher HOMA-IR and lower vigorous physical activity (Figure 5D). We found directions of association similar to the MAPK cascade across an expanded set of signaling pathway gene sets, though there was less consistency in the pattern of gene-set directions of association between total pseudobulk and bulk data for metabolic traits (Figure S26A).

Similar to the MAPK cascade gene sets, we observed that higher expression of protein K48-linked ubiquitination (Type 1 and bulk tissue), chromatin organization (Type 1, Type 2x, total pseudobulk), cytoskeleton organization (all muscle fiber types and total pseudobulk), mRNA processing (Type 1), and muscle structure development (all muscle fiber types and total pseudobulk) genes were associated with higher HOMA-IR and lower vigorous physical activity.

We found similar directions of association across related gene sets (Figures 5A, S27A, S28A, S29A). We observed that, as for gene expression, most of the significant gene set–trait associations for HOMA-IR and vigorous physical activity were still significant when both traits were included in the same model (Figures 4B, 4C, 4D, 4E, S25B, S26B, S27B, S28B, and S29B). Overall, the directions of association for gene set–trait associations for higher HOMA-IR were similar to other cardiometabolic traits, including higher BMI, higher 2-hour plasma glucose, and T2D compared to NGT.

### Chromatin accessibility–trait associations in skeletal muscle cell types

Gene expression is regulated in part by the extent of open, or accessible, chromatin at promoters and enhancers. To investigate how cardiometabolic and physical activity traits were associated with chromatin accessibility in the ten most abundant muscle cell types, we tested for association of cell type chromatin accessibility (snATAC-seq data) with traits using a negative binomial model. Within each muscle fiber type, we found the highest number and percentage of differentially expressed peaks for sex (male vs. female, 36919–62206 peaks, 5.1–12.7%), and far fewer for BMI (1200–3194 peaks, 0.2–0.5%) and HOMA-IR (471–1882 peaks, 0.1–0.4%) (Table S12 and S13). We did not assess the relationship between chromatin accessibility and gene expression due to the limited number of significant peaks associated with cardiometabolic and physical activity traits. Overall, we found minimal differences in skeletal muscle fiber chromatin accessibility across these traits, a stark contrast to fifteen times more associations by sex than other traits.

## Discussion

This study highlights that muscle fiber type abundance and expression of energy metabolism genes exhibit inverse directions of association for higher insulin resistance and higher physical activity. While we cannot determine which differences are detrimental or causal for disease phenotypes, it is possible that physical activity could counteract muscle fiber type-specific gene expression patterns observed in individuals with insulin resistance.

The FUSION Tissue Biopsy Study integrates single-nucleus and bulk skeletal muscle tissue data from the same 263 individuals with detailed cardiometabolic and physical activity phenotyping. We identified numerous gene expression–trait and gene set–trait associations at the cell type level and tissue level (total pseudobulk and bulk tissue) for cardiometabolic and physical activity traits.

In contrast, we identified a relatively smaller proportion of cell-type-level chromatin accessibility–trait associations. This suggests that differences in chromatin accessibility may play a relatively minor role in the steady-state regulation of gene expression in the context of cardiometabolic and physical activity traits, or that we have not detected more subtle differences in chromatin accessibility that affect gene expression.^36^

Overall, in single-nucleus data, lower values of cardiometabolic traits (BMI, HOMA-IR, NGT vs. T2D, and 2-hour glucose) and higher levels of both total and vigorous physical activity—traits typically associated with better health—showed similar directions of association with cell type abundance, as well as gene expression at the muscle fiber type and whole-tissue levels. For both HOMA-IR and physical activity, these directions of association persisted even after adjusting for the other trait. While insulin resistance pathways can be driven by physical inactivity, our results suggest that insulin resistance can act independently of physical activity on overlapping biological pathways.^37^

Lower insulin resistance and higher total physical activity were associated with higher proportions of Type 1 nuclei and lower proportions of Type 2x nuclei, confirming findings from previous smaller single-nucleus experiments and histological studies.^9–15^ Higher proportions of Type 1 and lower of Type 2x muscle fiber have previously been associated with healthier phenotypes, such as better metabolic function and increased fat utilization.^38,39^

Higher expression of energy metabolism genes was associated with lower cardiometabolic trait levels (BMI, HOMA-IR, NGT vs. T2D, 2-hour plasma glucose) and higher physical activity levels (total and vigorous) across muscle fiber types and in total pseudobulk data. We confirmed that previously-identified associations of insulin resistance and physical activity with energy metabolism-related gene sets in bulk tissue were also consistent across muscle fiber types and were not confounded by the other trait.^10,16–20^ Overall, we identified similar, shared gene expression–trait and gene set–trait associations between all muscle fiber types, from the oxidative Type 1 fibers to the glycolytic Type 2x fibers, for both insulin resistance (HOMA-IR) and vigorous physical activity. However, a substantial proportion of associated genes were significant in only one fiber type, highlighting potential nuances in the gene expression patterns that occur between similarly functioning, but highly specialized, muscle fiber types. Our statistical power to detect gene expression associations in non-muscle fiber types was lower, as single-nucleus experiments in skeletal muscle do not enrich for these less abundant cell types.

Gene expression–trait associations for HOMA-IR and vigorous physical activity in total pseudobulk and bulk tissue mostly recapitulated those associations identified in individual muscle fiber types, the most abundant cell types in skeletal muscle. Gene expression–trait associations between single-nucleus total pseudobulk (nucleic RNA) and bulk tissue (nucleic + cytoplasmic RNA) datasets were most similar for genes with higher expression levels in each experiment, suggesting some differences may reflect differences in statistical power to detect associations. In addition, differences between total pseudobulk and bulk tissue may reflect differences in the relative abundance of cell types between the two data types or trait-specific regulation of nuclear versus cytoplasmic mRNA.^40–42^ For example, energy metabolism-related gene set associations showed similar directions of association between total pseudobulk and bulk tissue for physical activity and T2D status; however, for BMI and insulin resistance (HOMA-IR), directions of association in bulk tissue were opposite those identified in the muscle fiber types and total pseudobulk, and contrary to previously-published bulk RNA-seq experiments.^18^ In contrast to the bulk RNA-seq results in Scott et al.^18^, our study adjusted for single-nucleus cell type proportions inferred from single-nucleus data, which may account for some of these differences. Skeletal muscle bulk tissue studies remain valuable for identifying gene expression associations that are consistent across muscle fiber types, but results may be influenced by differences in cell type abundance, especially for traits with cell-type-specific expression patterns.

The skeletal muscle biopsies in the FUSION Tissue Biopsy Study were obtained from older, Finnish individuals (mean age: 60.3 years), with >98% of participants older than 45 years of age. Thus, the results cannot be generalized to younger individuals. However, older individuals are at a greater risk for insulin resistance, and ultimately T2D.^43,44^ As older individuals typically have lower levels of physical activity than younger individuals, the observed range of physical activity may influence gene expression.^45,46^ We used self-reported physical activity, developed and validated in observational studies among Finnish adults. As such, we cannot assess the interventional effects of physical activity on gene expression at the cell type and bulk tissue level. However, the cell-type-level gene expression–physical activity associations observed in our study recapitulate previously-reported associations of energy metabolism gene sets from small-scale interventional single-nucleus studies and bulk tissue studies.^10,25,26^ Our study was enriched for individuals with newly-diagnosed, drug-naive T2D, and thus the results may not be fully representative of the general population. Their inclusion allowed us to test for differences between individuals with T2D and those with NGT. We were not able to perform direct replication of our results as we did not identify other single-nucleus skeletal muscle datasets that included measurements of insulin resistance, T2D status, and physical activity within the same individuals. Our data can serve as a well-phenotyped atlas of skeletal muscle.

Our results do not establish causal links between differences in cardiometabolic or physical activity traits and differences in gene expression levels.^47^ While RNA is the active form of some genes, the majority of genes exert their biological effects through their protein products. Protein isoforms can have distinct biological functions, and differences in protein levels can alter the activity of biological processes.^48,49^ Although changes in gene expression may not be causal for disease, they can indicate a deviation from an optimal or healthy cellular state. Gene expression profiles are tightly regulated and perturbation or misregulation may contribute to disease development.^50^ For example, we found that higher expression of MAPK cascade genes was associated with higher insulin resistance and lower vigorous physical activity across muscle fiber types and in total pseudobulk. While higher expression of these genes does not imply a disease state, it does suggest that the MAPK pathway may be perturbed; misregulation of MAPK signaling has been linked to increased risk of both cancer and diabetes.^51^ These findings enhance our understanding of the complex interplay between physical activity and insulin resistance in shaping skeletal muscle abundance and muscle fiber type-specific gene expression, and highlight a potential benefit of regular physical activity.

## Methods

### FUSION Tissue Biopsy Study

The Finland-United States Investigation of NIDDM Genetics (FUSION) Tissue Biopsy Study enrolled 331 living Finnish participants from three study sites in Finland (Helsinki, Kuopio, and Savitaipale) between 2009 and 2013. Clinical phenotyping was conducted during a clinical study visit that occurred on average two weeks (SD: 15 days) before the biopsy visit. During the biopsy visit, ∼250 mg of vastus lateralis skeletal muscle tissue was collected from each participant.

#### Further study details can be found in Scott et al.^18^

The FUSION Tissue Biopsy Study was approved by the ethics committee of the Hospital District of Helsinki and Uusimaa. All participants provided written informed consent.

### Glucose tolerance status

During the clinical visit, after a 12-hour overnight fast, participants underwent a 2-hour, 4-point oral glucose tolerance test with fasting serum insulin also measured. We characterized an individual’s glycemic status as normal glucose tolerance (NGT), impaired fasting glucose (IFG), impaired glucose tolerance (IGT), or type 2 diabetes (T2D), using the WHO diagnostic classification.^31^ Individuals with T2D were newly diagnosed and were not using glucose-lowering drugs. We approximated insulin resistance using the HOMA-IR method (Homeostatic Model Assessment for Insulin Resistance).^27^

### Leisure time physical activity

Participants completed a validated 12-month leisure time physical activity history questionnaire that included the most common types of physical activity among Finnish adults.^28–30^ For each activity, participants reported the mean intensity (on a scale ranging from light to very heavy), average duration of each physical activity session, and frequency per month. The energy expenditure of each activity was defined as the metabolic equivalent of task (MET) value for each intensity value based on empirical data. The weekly duration (hr/wk) of each activity was calculated as the product of the average weekly frequency and the average duration of the activity. The energy expenditure of each activity (METS*kg*hr/wk) was calculated as the product of the weekly duration (hr/wk), body weight of the participant (kg), and the MET value assigned to that activity. Total weekly energy expenditure (PHYS ACT: TOTAL) was calculated as the sum of energy expenditures for all activities. Weekly energy expenditure from vigorous activities was calculated as the sum of energy expenditures of activities with MET values ≥6, corresponding to vigorous-intensity physical activity as defined by the Compendium of Physical Activities.^52^

### Single-nucleus RNA-seq and ATAC-seq

We performed single-nucleus RNA-seq (snRNA-seq) and single-nucleus ATAC-seq (snATAC-seq) (on separate nuclei from the same isolation) in 287 frozen muscle tissue biopsy samples, processed in 10 batches.^33^ snRNA-seq reads were mapped using STARsolo^53^ and snATAC-seq reads were mapped using the Burrows-Wheeler Aligner (BWA-MEM)^54^ to the human genome (hg38) reference. After applying droplet-level quality control thresholds, there were 180,583 RNA-seq nuclei and 268,543 ATAC-seq nuclei across all samples. Ambient RNA decontamination for snRNA-seq droplets was carried out with DecontX.^55^ We used Liger^56^ to jointly cluster snRNA-seq and snATAC-seq nuclei and used known cell type markers to identify the clusters as adipocyte, endothelial cell, macrophage, fibro-adipogenic progenitor cell, mixed muscle fiber, neuromuscular junction (muscle fiber), neuronal cell, satellite cell, smooth muscle cell, T cell, Type 1 muscle fiber, Type 2a muscle fiber, and Type 2x muscle fiber. We removed the mixed muscle fiber type cluster from further analyses due to overrepresentation of exonic reads.^33^ We also excluded samples (a) with <100 snRNA-seq nuclei or <100 snATAC-seq nuclei (n=2), (b) from one non-Finnish individual, (c) from one individual from each of two first-degree relative pairs, and (d) from individuals who did not consent to be included in the Terveyden ja Hyvinvoinnin Laitos (THL) Biobank (n=19). After exclusions, we retained 168,309 snRNA-seq nuclei and 242,069 snATAC-seq nuclei from 263 participants.

### Bulk RNA-seq

Detailed methods for sample collection, processing, and quality control for bulk RNA-seq can be found in Scott et al.^18^ Briefly, total RNA from 301 frozen muscle tissue biopsy samples was sequenced to a mean depth of 91.3M strand-specific paired-end reads per sample.

### Differential cell type abundance by phenotypic traits

We tested for associations between cell type proportions and sex (male vs. female, n=263), BMI (n=263), HOMA-IR (n=263), 2-hour glucose (n=262), T2D vs. NGT (n=164), total physical activity (n=262), and vigorous physical activity (n=262) in three sets of nuclei: snRNA-seq, snATAC-seq, and combined snRNA-seq + snATAC-seq datasets. For each cell type and for the combined muscle fiber types (1 + 2a + 2x), we used negative binomial models to test the association between the number of nuclei and each trait using the glm.nb() function from the MASS package (version 7.3-65).^57^ We adjusted for sex, age, batch, and city of collection, and included the log10 number of total nuclei as an offset in all models. We inverse normalized all continuous traits to reduce the effect of extreme values. To assess potential confounding, we tested models adjusting for both HOMA-IR and total physical activity. We corrected for multiple testing by applying a false discovery rate (FDR) threshold of 5% across all trait–cell type associations within each set of nuclei.

### Single-nucleus cell type differential gene expression by phenotypic traits

We tested single-nucleus cell-type-level gene expression data for association with continuous cardiometabolic traits, physical activity, sex (male vs. female), age, and T2D vs. NGT. To create cell-type-level pseudobulk counts, for each participant we summed the UMI counts for each gene across all nuclei of a cell type. We excluded genes with zero counts for more than 75% of samples by cell type. For each cell type, we excluded samples with <10 snRNA nuclei of that cell type. We excluded adipocytes and T cells from the analysis for all traits, and macrophage and neuronal cells from the analysis of T2D status as fewer than 25 samples had at least 10 nuclei.

Across cell types, the number of samples with ≥10 nuclei ranged from 2 in T cells to 263 in Type 1 (Table S14). The mean number of snRNA nuclei per sample ranged from 12.5 in T cells to 258.0 in Type 1 (Table S14). The number of genes tested ranged from 11,490 to 23,849 (Table S15).

For each cell type, we used a negative binomial model to test for the association between gene expression counts and a trait. Sample size varied across models depending on trait availability (Table S16). We ran the analysis using DESeq2 (version 1.44.0) with the single cell recommendations (likelihood ratio test and setting the minmu parameter to 1x10^-6^).^58^ We adjusted for batch, city of collection, median mitochondrial fraction (across nuclei), total number of RNA nuclei, sex, and age. To test for confounding, we included additional models adjusted for both HOMA-IR and vigorous physical activity. We inverse normalized all continuous covariates to remove the possible effect of extreme values. We corrected for multiple testing by applying an FDR threshold of 5% across all gene expression–trait associations for a given trait and cell type. No independent filtering was performed when using DESeq2.

### Single-nucleus pseudobulk differential gene expression by phenotypic traits

For each individual, we created total pseudobulk counts by summing the UMI counts for each gene across all nuclei. We excluded genes with <5 counts in >75% of the samples, and retained all samples with ≥100 nuclei. This filtering results in 18,618 genes tested in 263 samples with a mean number of 635.3 nuclei per sample.

We used a negative binomial model to test for associations between gene expression counts and each trait. Sample size varied across models depending on trait availability (Table S16). Because these gene expression data were less sparse than those at the cell type level, we conducted the analysis using DESeq2 (version 1.44.0), following bulk RNA recommendations (Wald test instead of likelihood ratio test).^58^ Models were adjusted for the observed cell type proportions and covariates used in the cell-type-level analysis. To assess potential confounding, we also ran additional models, adjusted for both HOMA-IR and vigorous physical activity. We corrected for multiple testing by applying an FDR threshold of 5% across all gene expression–trait associations for a given trait. No independent filtering was performed when using DESeq2.

### Bulk tissue differential gene expression by phenotypic traits

To enable direct comparisons with total pseudobulk data, we restricted the bulk RNA-seq dataset to samples that were included in the single-nucleus analysis. We excluded genes with <5 counts in >75% of the samples, resulting in 22,309 genes tested in 252 samples.

We used a negative binomial model to test for the association between bulk gene expression counts and each trait. Sample size varied across models depending on trait availability (Table S16). We implemented the analysis using DESeq2 (version 1.44.0) with the bulk RNA recommendations (Wald test instead of likelihood ratio test).^58^ We adjusted for sex, age, median insert size, mean RNA integrity number (RIN), median transcript integrity number (TIN), batch, city of collection, mean GC content, and cell type proportions from the single-nucleus experiment. To test for the possibility of confounding, additional models adjusted for both HOMA-IR and vigorous physical activity. We corrected for multiple testing by applying an FDR threshold of 5% across all gene expression–trait associations for a given trait. No independent filtering was performed when using DESeq2.

### Downsampling Type 1 fiber gene counts

To approximate the power to detect gene expression–trait associations in other cell types, we downsampled gene counts from Type 1 muscle fiber in additional models. For each cell type comparison, we only included individuals in the downsampled Type 1 analysis who were also included in the analysis for the respective target cell type. To downsample the total UMIs for Type 1 to each target cell type, we multiplied the Type 1 UMIs for each gene (rounded to the nearest integer) by the total UMIs in the target cell type divided by the total UMIs in Type 1 muscle fiber. We tested for differential gene expression by trait in each of the downsampled Type 1 samples using DESeq2 (version 1.44.0) as previously described.

### Gene set enrichment analysis by phenotypic trait

We used RNA-Enrich to identify Gene Ontology (GO) terms enriched for genes with higher or lower expression with trait values.^35^ Briefly, RNA-Enrich models the association between GO term membership and the signed −log10 p-value of gene expression–trait association while adjusting for each gene’s mean read count using a logistic regression model. We mapped ENSEMBL IDs to Entrez IDs using the library org.Hs.eg.db version 3.20.0. We acquired GO term definitions using the libraries org.Hs.eg.db version 3.20.0 and GO.db version 3.20.0. We ran models separately for each trait and cell type combination. We considered all GO biological process gene sets (n=6486) containing 10–500 genes in at least one dataset (muscle fiber type, total pseudobulk, or bulk tissue). To reduce the potential impact of extreme values on gene set enrichment results, we winsorized all p-values <10⁻²⁵ to 10⁻²⁵ prior to running RNA-Enrich (only gene expression–sex associations had p-values <10⁻²⁵ and were winsorized). To assess the independence of results for a given trait, we also ran RNA-Enrich on gene expression–trait associations in a subset of models adjusted for a second trait. We corrected for multiple testing by applying an FDR threshold of 5% across all gene set–trait associations for a given trait and cell type.

### Comparison of sets of gene expression–trait and gene set–trait associations

We compared gene expression–trait and gene set–trait associations within each cell type by plotting the signed (by beta coefficient) −log10 p-values of association for trait 1 on the x-axis and trait 2 on the y-axis. To quantify these associations, we calculated the Spearman correlation between the signed −log10 p-values of association for pairs of traits using the cor.test() function in R, restricting to complete observations. We tested these correlation coefficients against a null hypothesis that the correlation is equal to 0. We corrected for multiple testing using a Bonferroni correction. When testing across all pairwise combinations of eight traits across eight cell types, the significance threshold was set to 0.05/(28*8), where 28 represents the number of unique trait pairs.

We compared gene expression–trait and gene set–trait associations between cell types for the same trait by first plotting the signed −log10 p-values of association for cell type 1 on the x-axis and cell type 2 on the y-axis. To quantify these relationships, we calculated the Spearman correlation between the signed −log10 p-values of association for pairs of cell types using the cor.test() function in R, restricting to complete observations. We tested these correlation coefficients against a null hypothesis that the correlation is equal to 0. We corrected for multiple testing using a Bonferroni correction. When testing all pairwise comparisons of 12 cell types (including total pseudobulk and bulk tissue) across two traits (HOMA-IR and vigorous physical activity), the significance threshold was set to 0.05/(66*2), where 66 represents the number of unique cell type pairs.

### Single-nucleus cell type differential chromatin accessibility by phenotypic traits

We tested single-nucleus cell-type-level chromatin accessibility data for associations with continuous cardiometabolic traits, physical activity, sex (male vs. female), age, and T2D vs. NGT. To generate cell-type-level pseudobulk peak counts, for each individual we summed the peak counts from all nuclei of a given cell type. ATAC peak calls were harmonized to create a list of consensus peak summits across cell types.^33^ For each cell type, we included only samples with ≥10 nuclei of that cell type. No cell types were excluded from this analysis. We excluded peaks with a mean peak count <1. Across cell types, the number of samples with ≥10 nuclei ranged from 73 in macrophages to 263 in Type 1 muscle fiber (Table S14). The mean number of ATAC nuclei per sample ranged from 16.8 in macrophages to 273.9 in Type 1 (Table S14). The number of peaks tested ranged from 20,706 to 927,588 (Table S15).

For each cell type, we used a negative binomial model to test for the association between peak counts and each trait. Sample size varied across models depending on the trait availability (Table S17). We ran the analysis using DESeq2 (version 1.44.0) following the single cell recommendations.^58^ Models adjusted for batch, city of collection, median mitochondrial fraction (across nuclei), transcription start site enrichment, total number of ATAC nuclei, sex (male vs. female), and age. We inverse normalized all continuous covariates to remove the possible effect of extreme values. We corrected for multiple testing by applying an FDR threshold of 5% across all peak–trait associations for a given trait and cell type. No independent filtering was performed when using DESeq2.

## Acknowledgements

The samples/data used for the research were obtained from THL Biobank (study number THLBB2025_5). We thank all study participants for their generous participation in Finnish health research.

This work was supported by the National Institutes of Health grants T32HG000040 (NHGRI) to M.B. (D.L.C., S.C.H., and M.D.S. supported), R01DK072193 (NIDDK) to K.L.M., UM1DK126185 (NIDDK) to K.L.M. and S.C.J.P., U01DK062370 (NIDDK) to M.B. and L.J.S., R01HG009976 (NHGRI) to M.B., R01DK117960 (NIDDK) to S.C.J.P., and P30AR069620 (NIAMS) to S.C.J.P., and by the Foundation for the National Institutes of Health (FNIH) grant N033979 to S.C.J.P. and L.J.S.

## Author contributions

Conceptualization, D.L.C., A.V., S.C.J.P., and L.J.S.; study design, M.R.E., J.S., M.L., J.T., T.A.L., M.B., F.S.C., H.A.K., S.C.J.P., and L.J.S; data curation, A.V., M.R.E., N.M., H.M.S., P.O., N.N., L.L.B., and M.D.S.,; formal analysis, D.L.C. and S.C.H..; funding acquisition, M.L., J.T., T.A.L., K.L.M., M.B., F.S.C., H.A.K., S.C.J.P., and L.J.S.; investigation, D.L.C., S.C.H., A.V., M.R.E., N.M., H.M.S., E.M.H-B., N.N., L.L.B., and M.D.S.; methodology, D.L.C., S.C.H., A.V., P.O., M.D.S., and L.J.S.; visualization, D.L.C.; supervision, S.C.J.P. and L.J.S.; writing—original draft, D.L.C. and L.J.S.; writing—review & editing, D.L.C., S.C.H., H.M.S., E.M.H-B., J.T., T.A.L., M.B., H.A.K., and L.J.S.

## Supplemental information

Supplementary materials includes: Figures S1 to S29

Tables S1-S2, S4, S6-S9, S11-12, S14-17

Other supplementary materials for this manuscript include the following: Tables S3, S5, S10, and S13

