## Supplementary Material for "Inverse directions of association of higher physical activity and higher insulin resistance with human skeletal muscle cell typae abundance and fiber-type-level gene expression"

The PDF file includes:

- Figures S1 to S29
- Tables S1-S2, S4, S6-S9, S11-12, S14-17

Other Supplementary Materials for this manuscript include the following:

- Tables S3, S5, S10, and S13

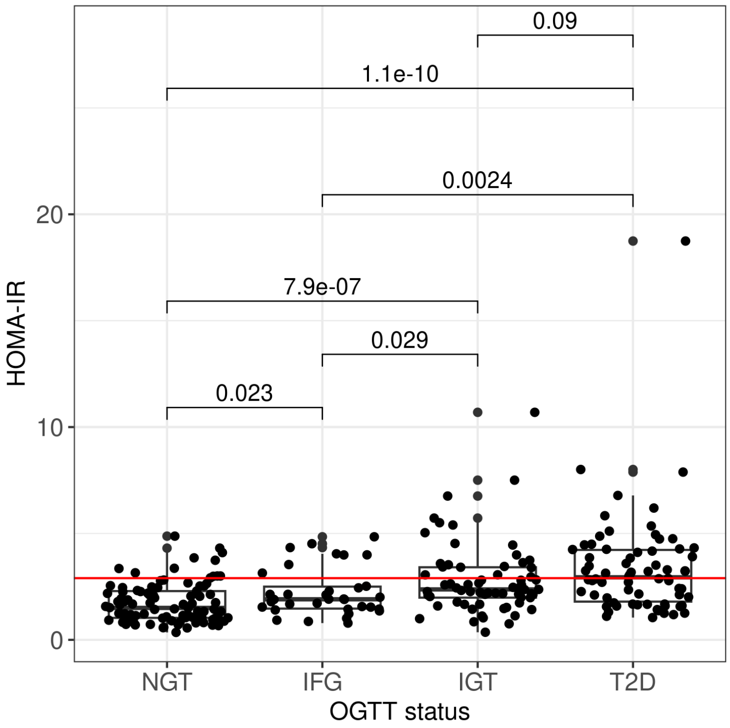


**Figure S1. Boxplots of HOMA-IR by oral glucose tolerance test (OGTT) status.** Each individual's oral glucose tolerance test (OGTT) status was characterized as normal glucose tolerance (NGT), impaired fasting glucose (IFG), impaired glucose tolerance (IGT), and type 2 diabetes (T2D). The red line denotes HOMA-IR equal to 2.9 where individuals with HOMA-IR>2.9 are considered to have clinically high insulin resistance. To test for differences between groups, we used the Wilcoxon rank-sum test. p-values<0.05/6 are considered significant after a Bonferroni correction.

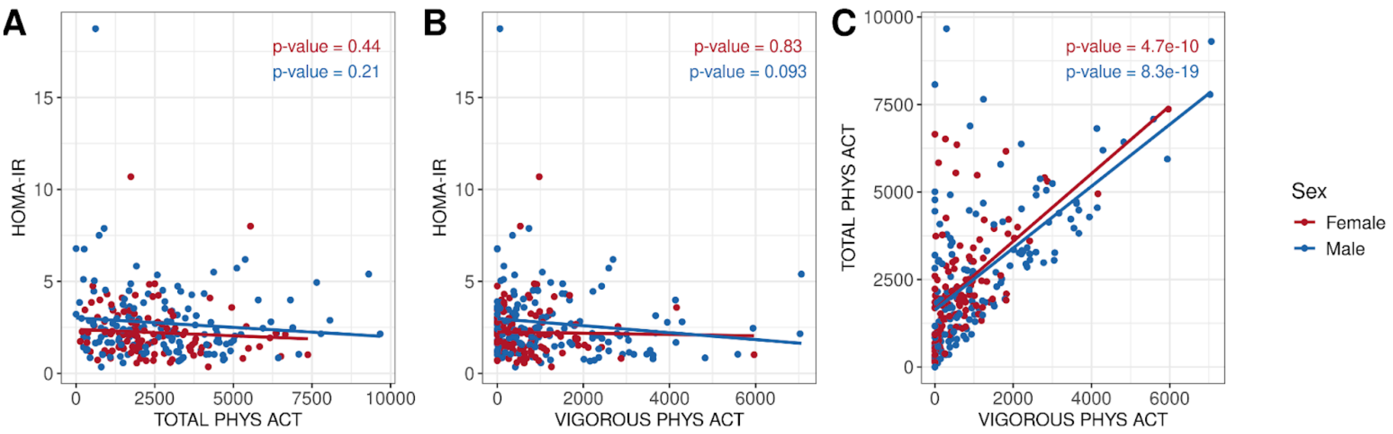


**Figure S2. Scatter plots of trait values.** (A) Scatter plot of total physical activity and HOMA-IR. (B) Scatter plot of vigorous physical activity and HOMA-IR. (C) Scatter plot of vigorous physical activity and total physical activity. Each point represents an individual colored by sex. Each line represents a simple linear regression line colored by sex. p-values<0.05/6 are significant after a Bonferroni correction.

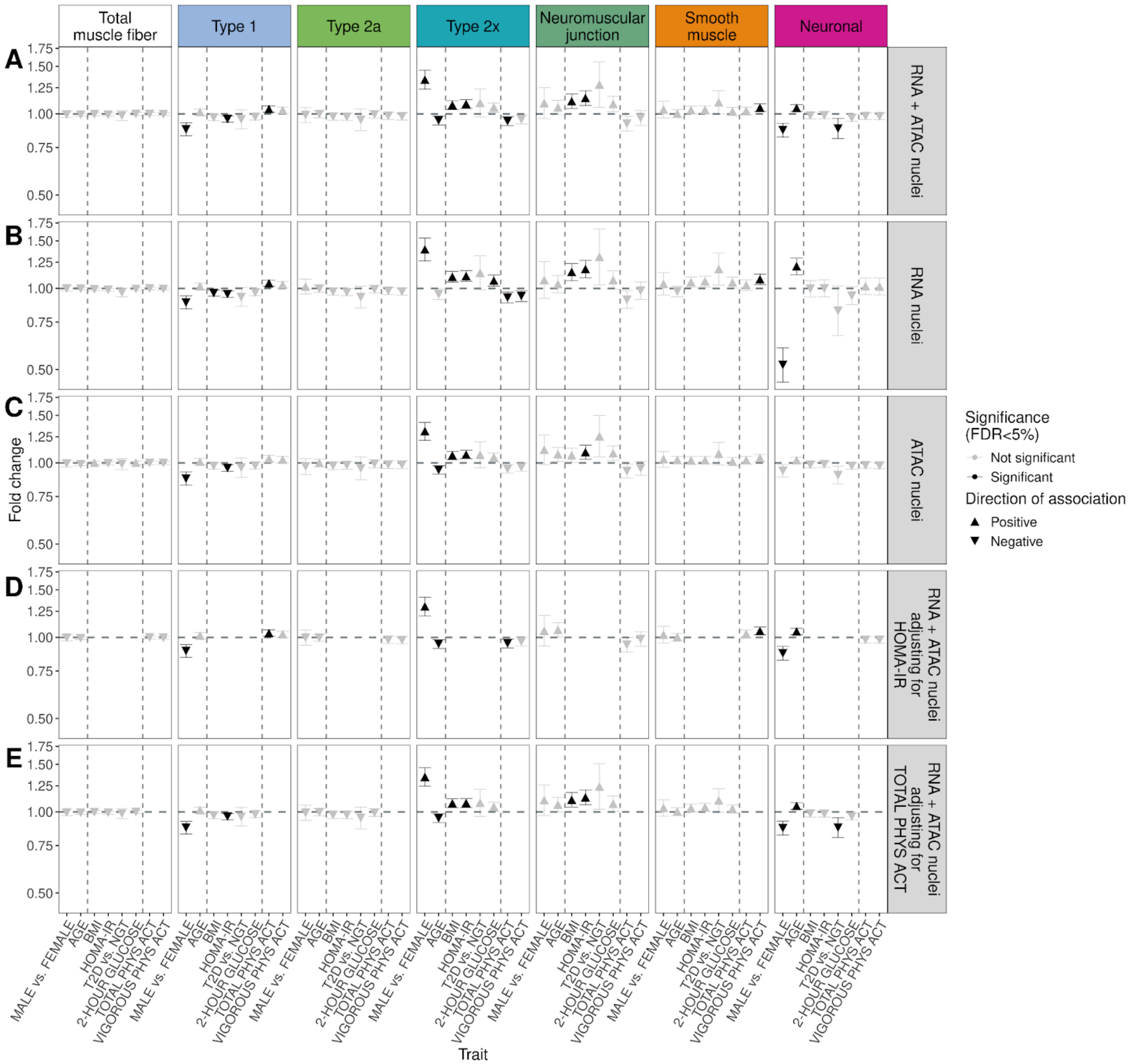


**Figure S3. Differences in cell type abundance of human skeletal muscle.** (A, B, C) Fold change and 95% confidence intervals comparing the number of (A) combined snRNA+snATAC nuclei (B) snRNA nuclei or (C) snATAC nuclei by trait across cell types and total muscle fiber. (D, E) Fold change and 95% confidence intervals comparing the number of snRNA and snATAC nuclei by trait across cell types and total muscle fiber while adjusting for (D) HOMA-IR or (E) TOTAL PHYS ACT in addition to base covariates. Significance was calculated as FDR<5% across all tests for each panel individually. Sample sizes for each trait can be found in the methods.

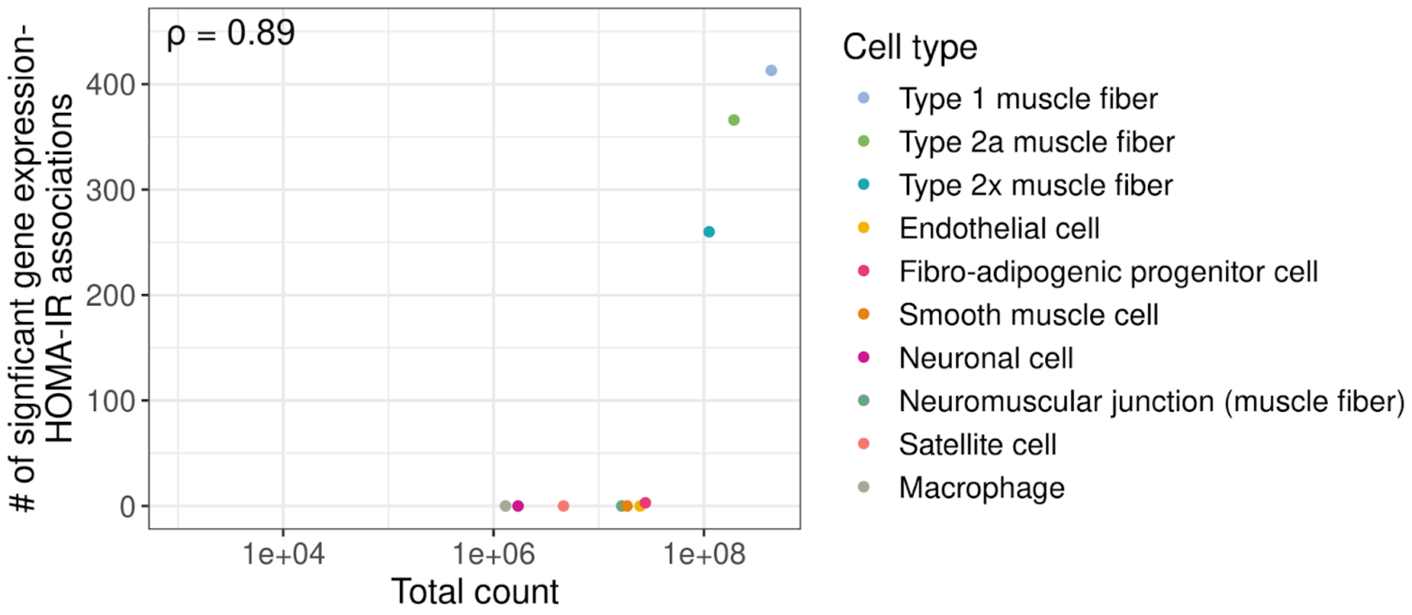


**Figure S4. Number of significant gene expression–HOMA-IR associations by total gene counts observed in each cell type.** Spearman correlation was calculated between all cell types.

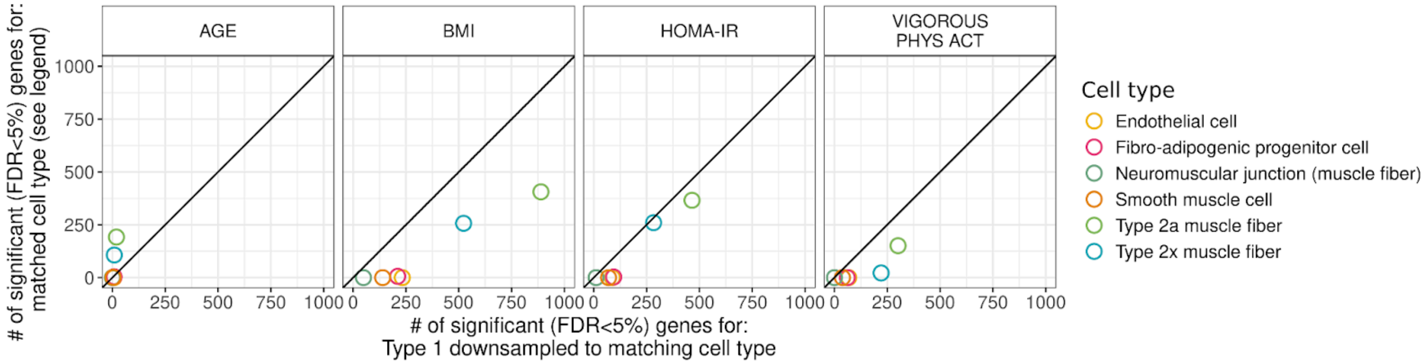


**Figure S5. Scatter plots of number of significant (FDR<5%) genes per trait for Type 1 sample size and UMI count downsampled to the matched cell type.** Type 1 was downsampled to the same mean gene count and sample size as the matched cell type, and gene expression analysis was repeated on the downsampled dataset. The x-axis is the number of significant genes for Type 1 downsampled to the matching cell type while the y-axis is the number of significant genes for the matched cell type. The color of the point denotes the cell types matched to.

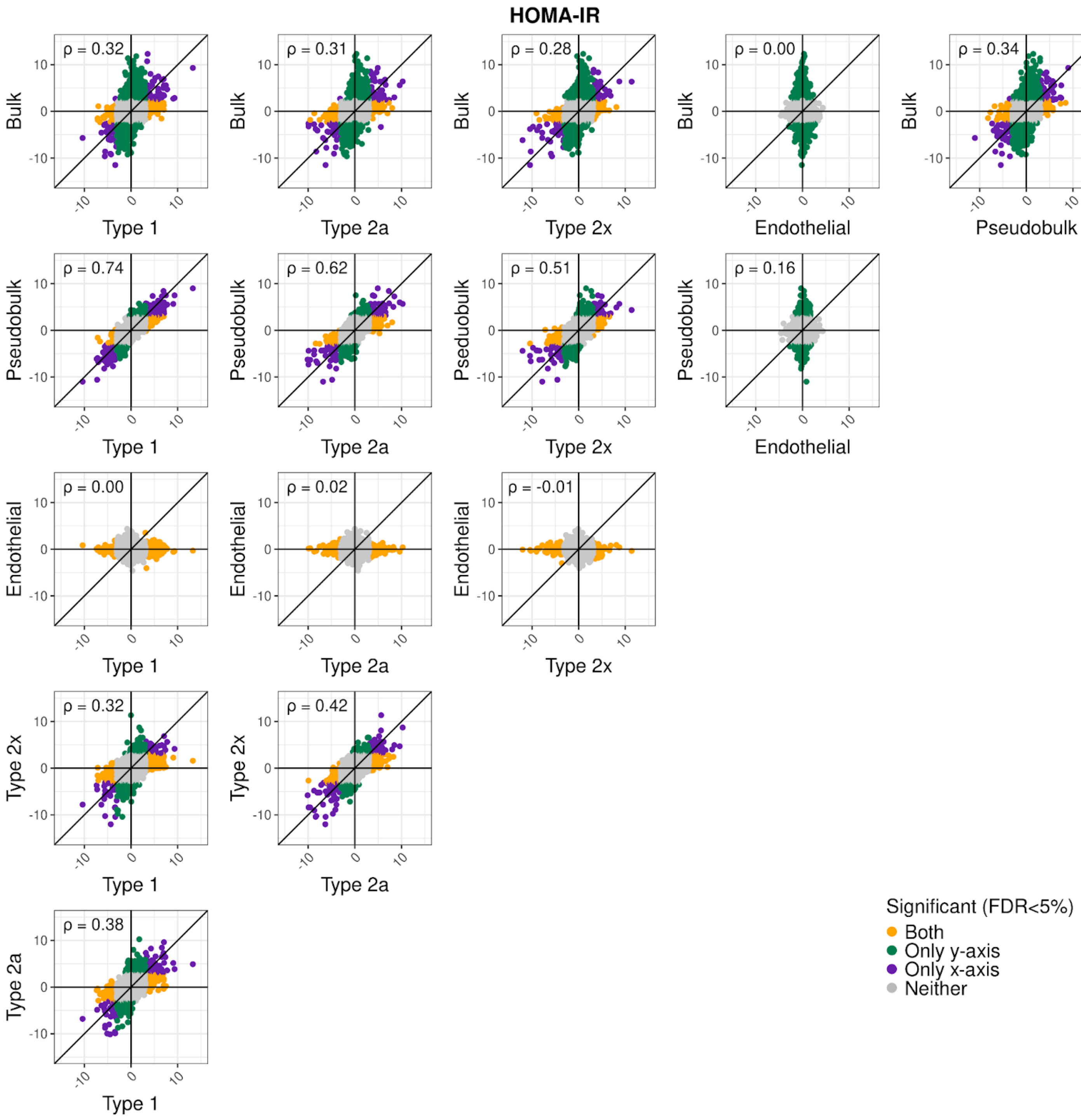


**Figure S6. Scatter plots of signed −log10 p-value of gene expression–HOMA-IR associations between muscle fiber types, endothelial cells, total pseudobulk (pseudobulk), and bulk tissue.** Each point is a gene and is colored based on which cell types the gene is significant (FDR<5%) in.

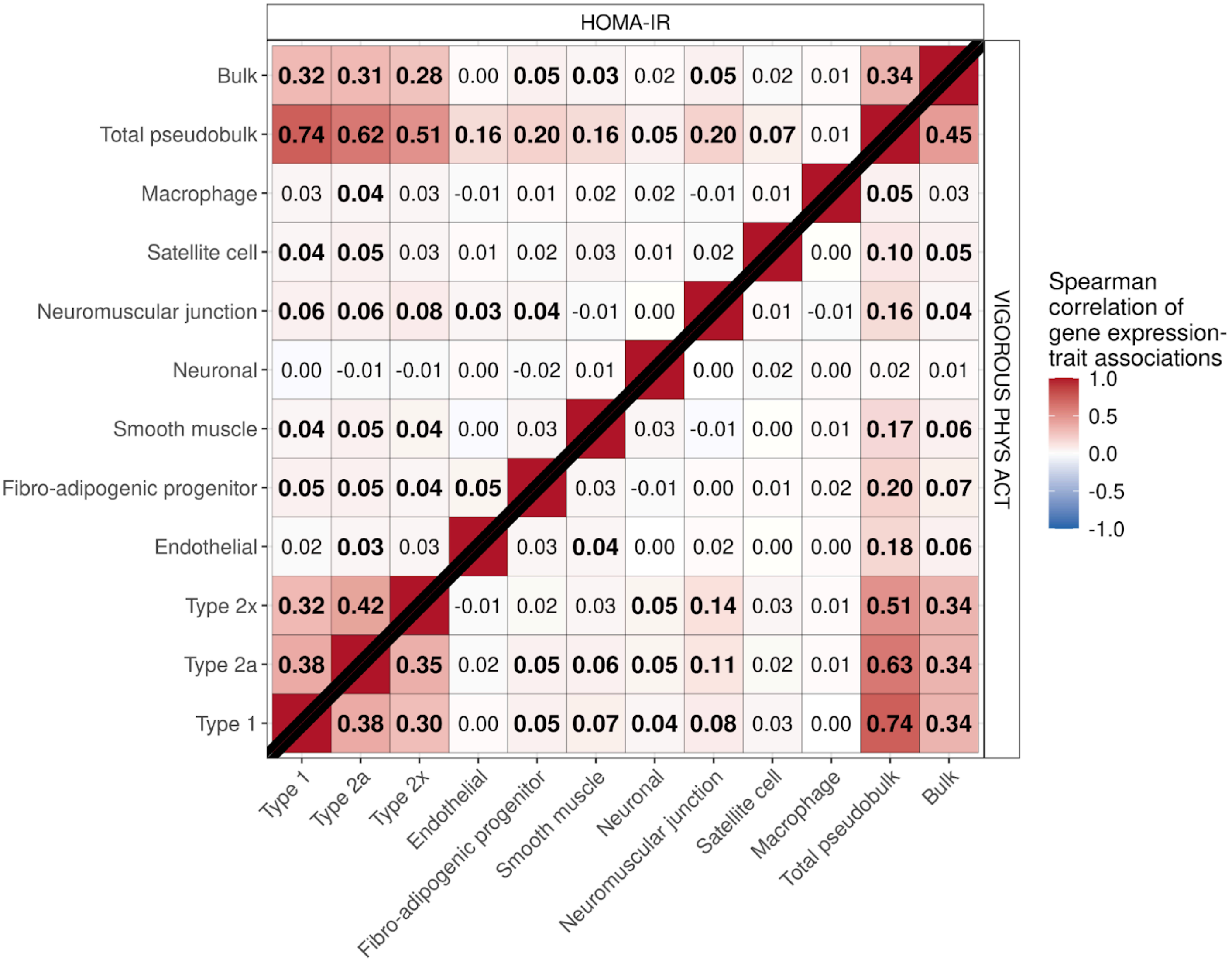


**Figure S7. Heatmap of Spearman correlations of gene expression–trait associations between cell types for (top left) HOMA-IR and (bottom right) vigorous physical activity.** Spearman correlations were calculated between the signed (by beta coefficient) −log10 of gene expression–trait associations. Numbers in bold represent correlations significantly different than 0 tested at a Bonferroni-adjusted significance level of $p<0.05/132$.

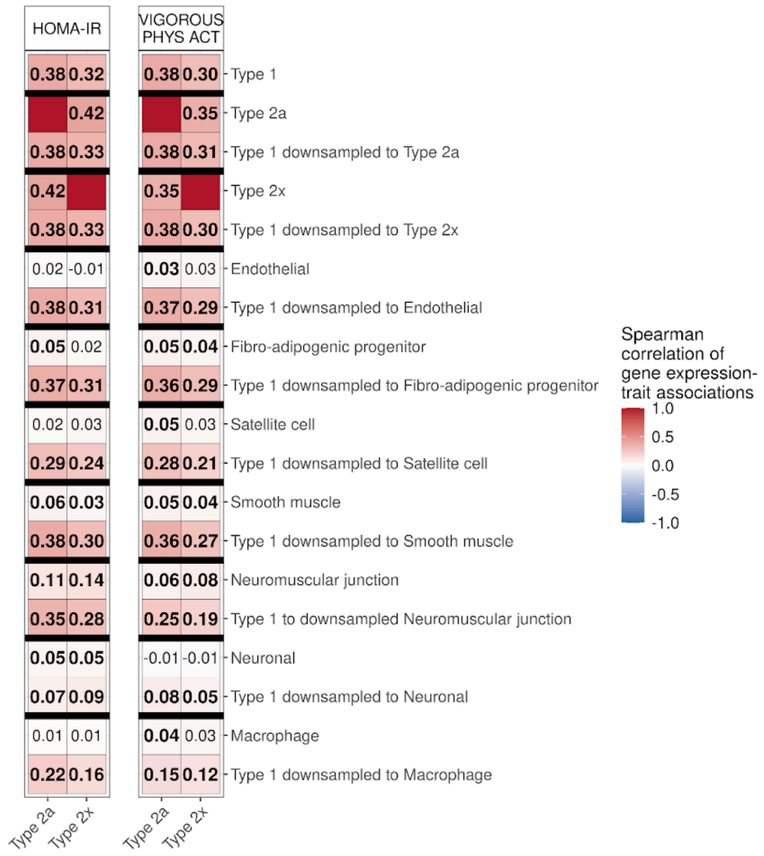


**Figure S8. Heatmap of Spearman correlations of gene expression–trait associations between Type 2a or Type 2x and the original Type 1 dataset or Type 1 datasets downsampled to match other cell types.** Type 1 was downsampled to the same mean gene count and sample size as the matched cell type and gene expression analysis was repeated on the downsampled dataset. Spearman correlations were calculated between the signed (by beta coefficient) −log10 of (left) gene expression–HOMA-IR and (right) gene expression–vigorous physical activity associations. Numbers in bold represent correlations significantly different than 0 tested at a Bonferroni adjusted significance level of $p<0.05/66$.

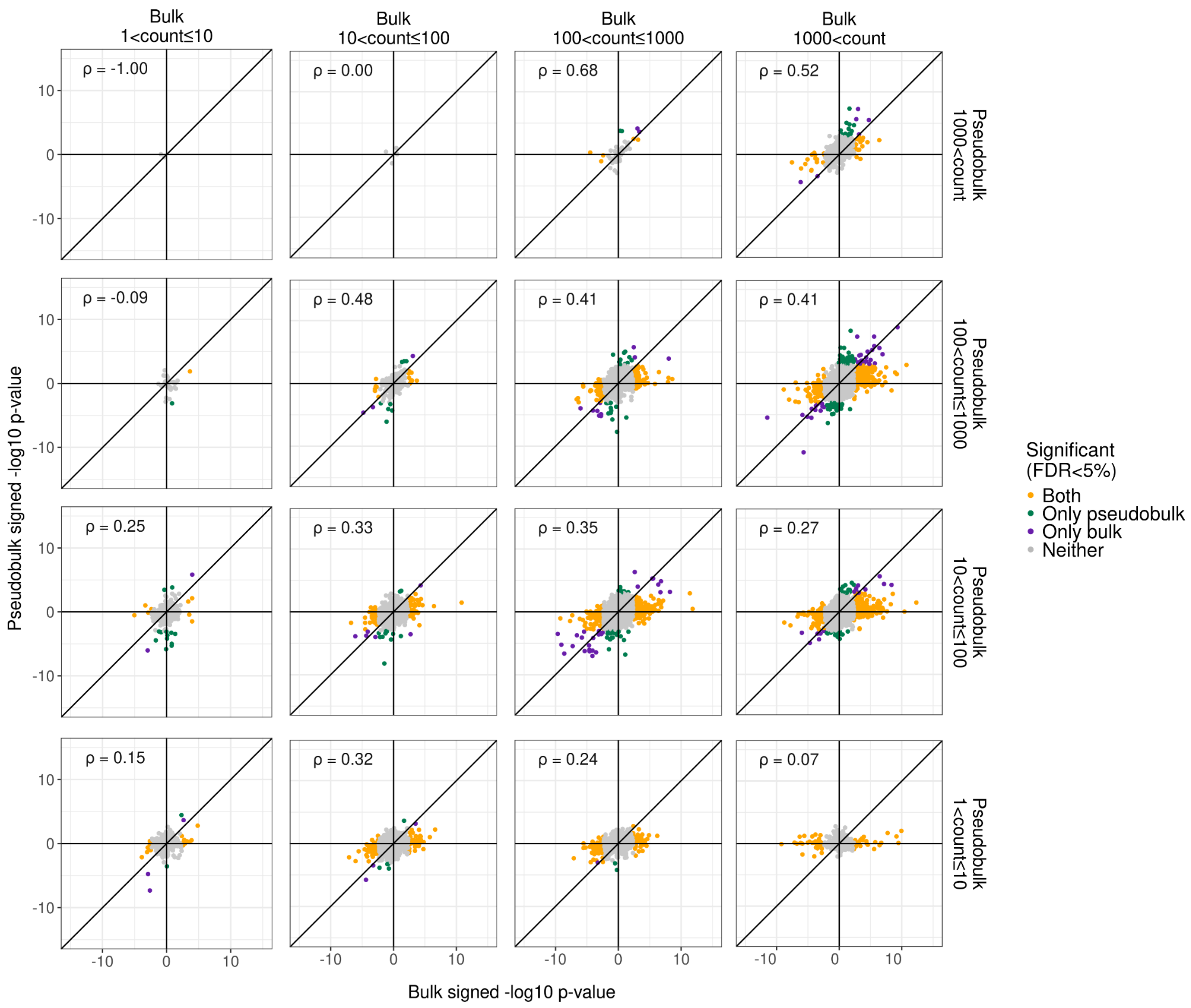


**Figure S9. Scatter plots of signed −log10 p-value of gene expression–HOMA-IR associations between total pseudobulk (pseudobulk) and bulk tissue binned by average gene count in each aggregated dataset.** Genes were binned based on the average gene count in total pseudobulk and bulk tissue. Each point is a gene and is colored based on which aggregated dataset the gene is significant (FDR<5%) in.

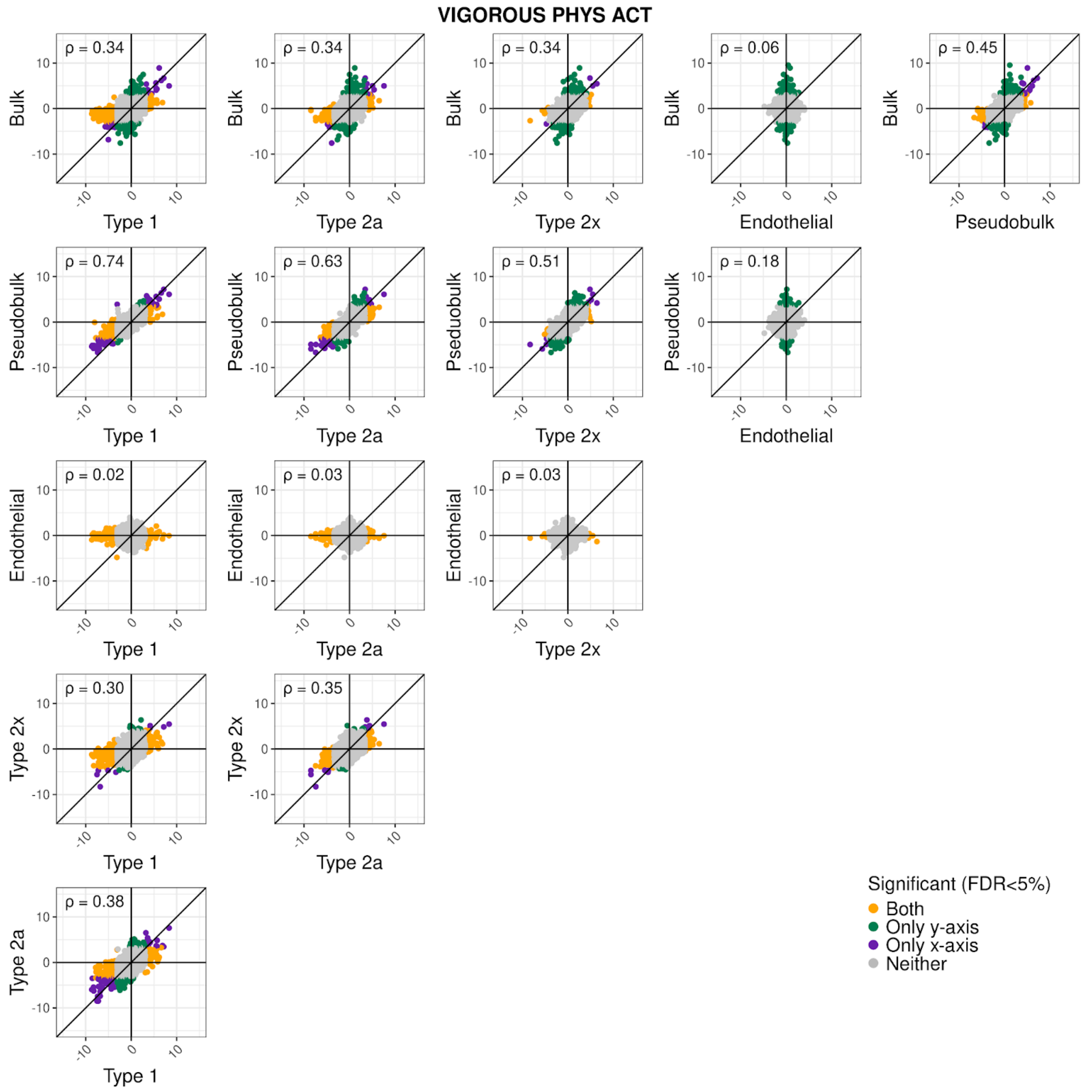


**Figure S10. Scatter plots of signed −log10 p-value of gene expression–vigorous physical activity associations between muscle fiber types, endothelial cells, total pseudobulk (pseudobulk), and bulk tissue.** Each point is a gene and is colored based on which cell types the gene is significant (FDR<5%) in.

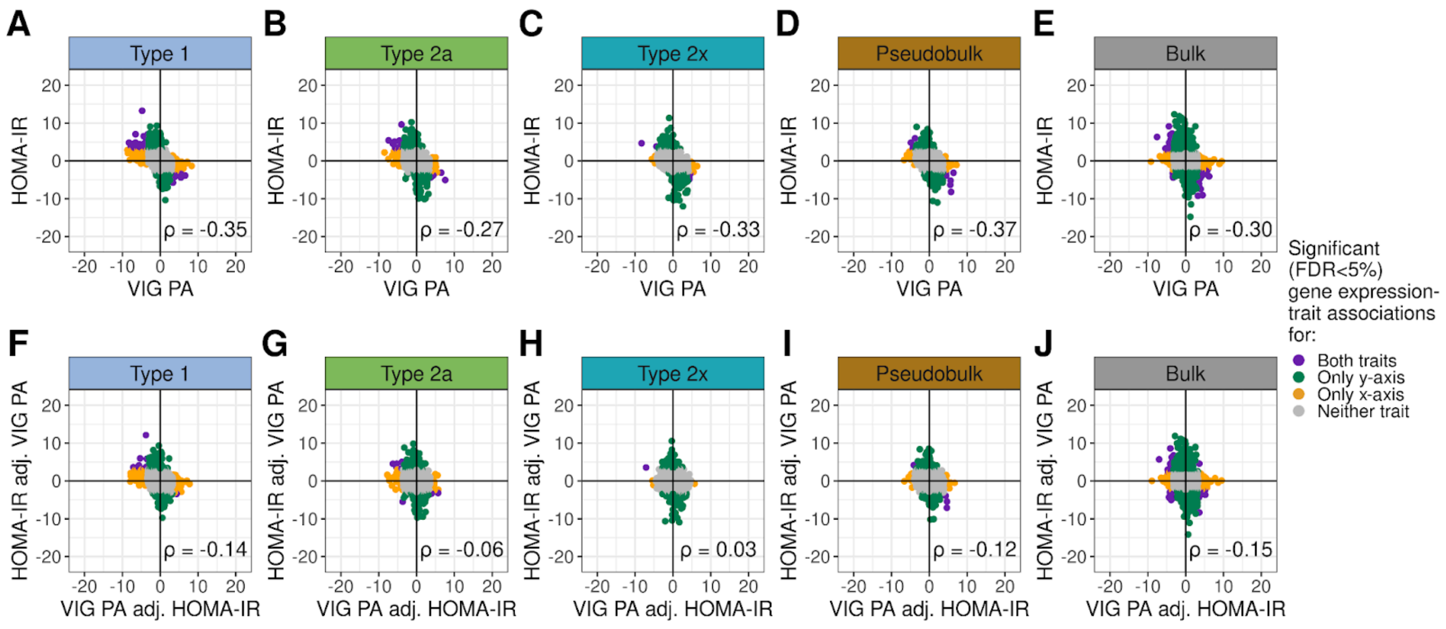


**Figure S11. Scatter plots of signed −log10 p-value of gene expression–trait associations for vigorous physical activity vs. HOMA-IR.** (A-E) Signed −log10 p-value of gene expression–trait associations for vigorous physical activity (VIG PA) vs. HOMA-IR in models only adjusting for base covariates in muscle fiber types, total pseudobulk (pseudobulk), and bulk tissue. (F-J) Signed −log10 p-value of gene expression–trait associations for vigorous physical activity and HOMA-IR when adjusting (adj.) for both traits in addition to base covariates in the same model in muscle fiber types, total pseudobulk, and bulk tissue. Each point is a gene and is colored based on which cell types the gene is significant (FDR<5%) in.

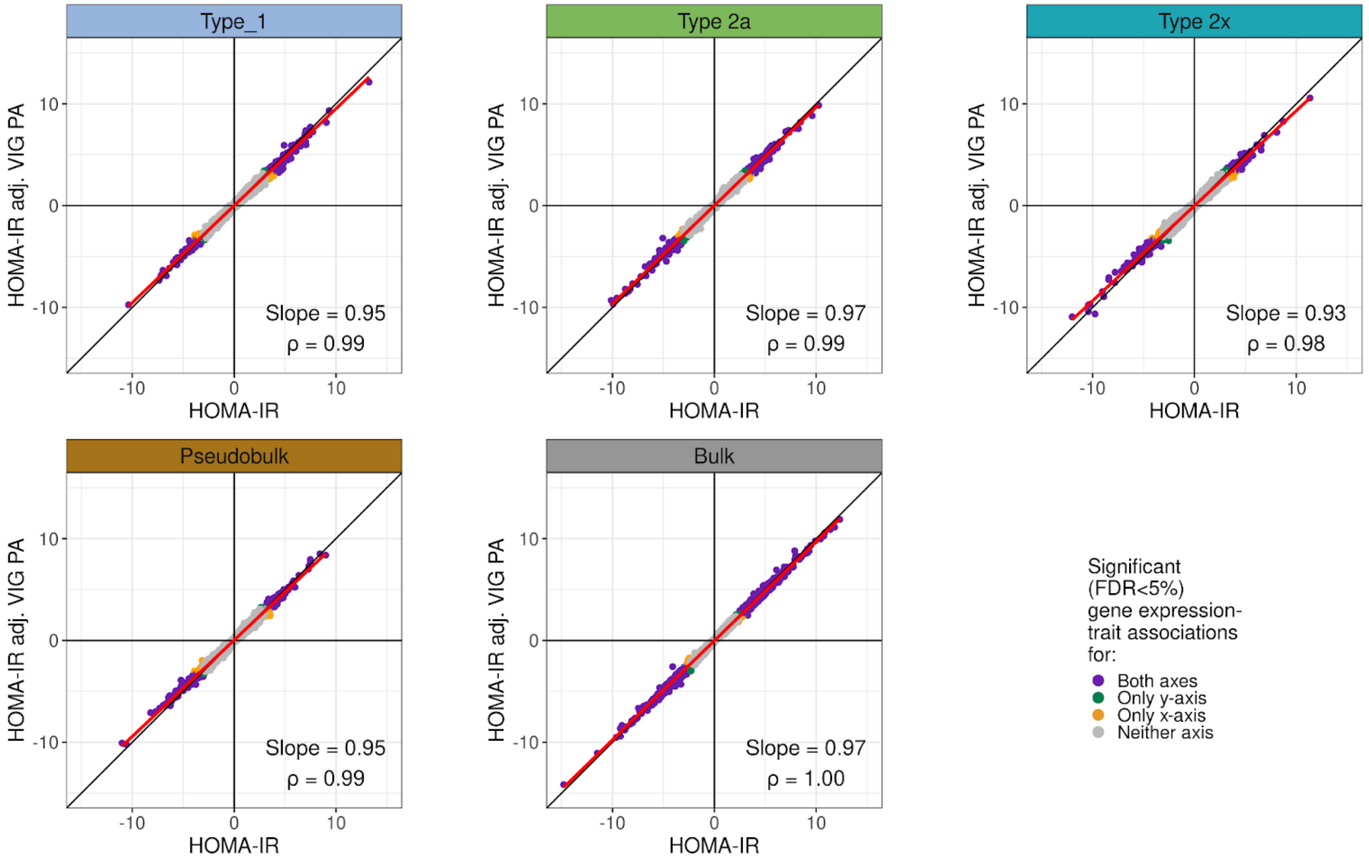


**Figure S12. Scatter plots of signed −log10 p-value of gene expression–HOMA-IR associations with and without adjusting (adj.) for vigorous physical activity (VIG PA) in the muscle fiber types, total pseudobulk (pseudobulk), and bulk tissue.** Each point is a gene and is colored based on which axis the gene is significant (FDR<5%) in. The red line represents a simple linear regression line with the slope of the line indicated in each plot.

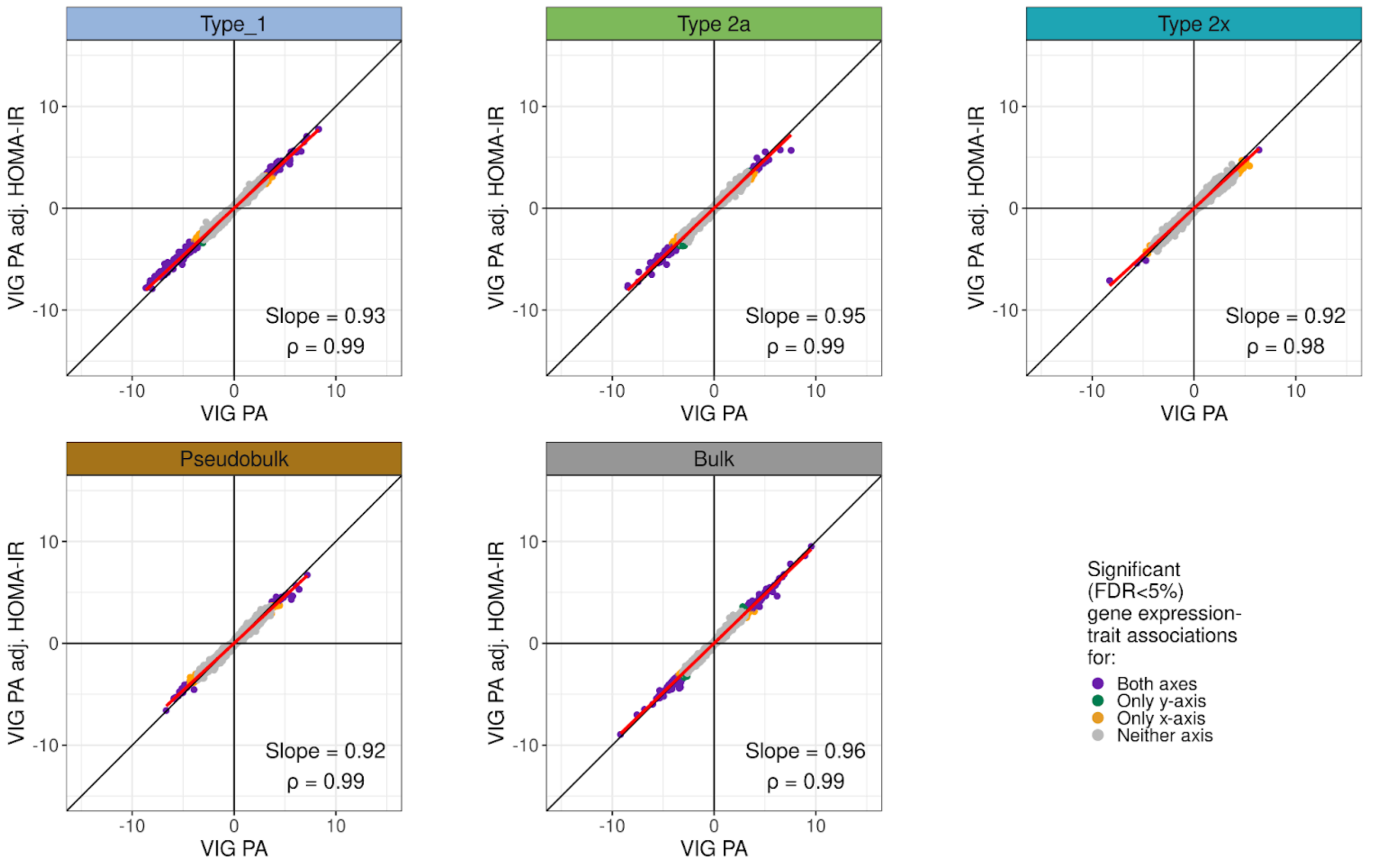


**Figure S13. Scatter plots of signed −log10 p-value of gene expression–vigorous physical activity (VIG PA) associations with and without adjusting (adj.) for HOMA-IR in the muscle fiber types, total pseudobulk (pseudobulk), and bulk tissue.** Each point is a gene and is colored based on which axis the gene is significant (FDR<5%) in. The red line represents a simple linear regression line with the slope of the line indicated in each plot.

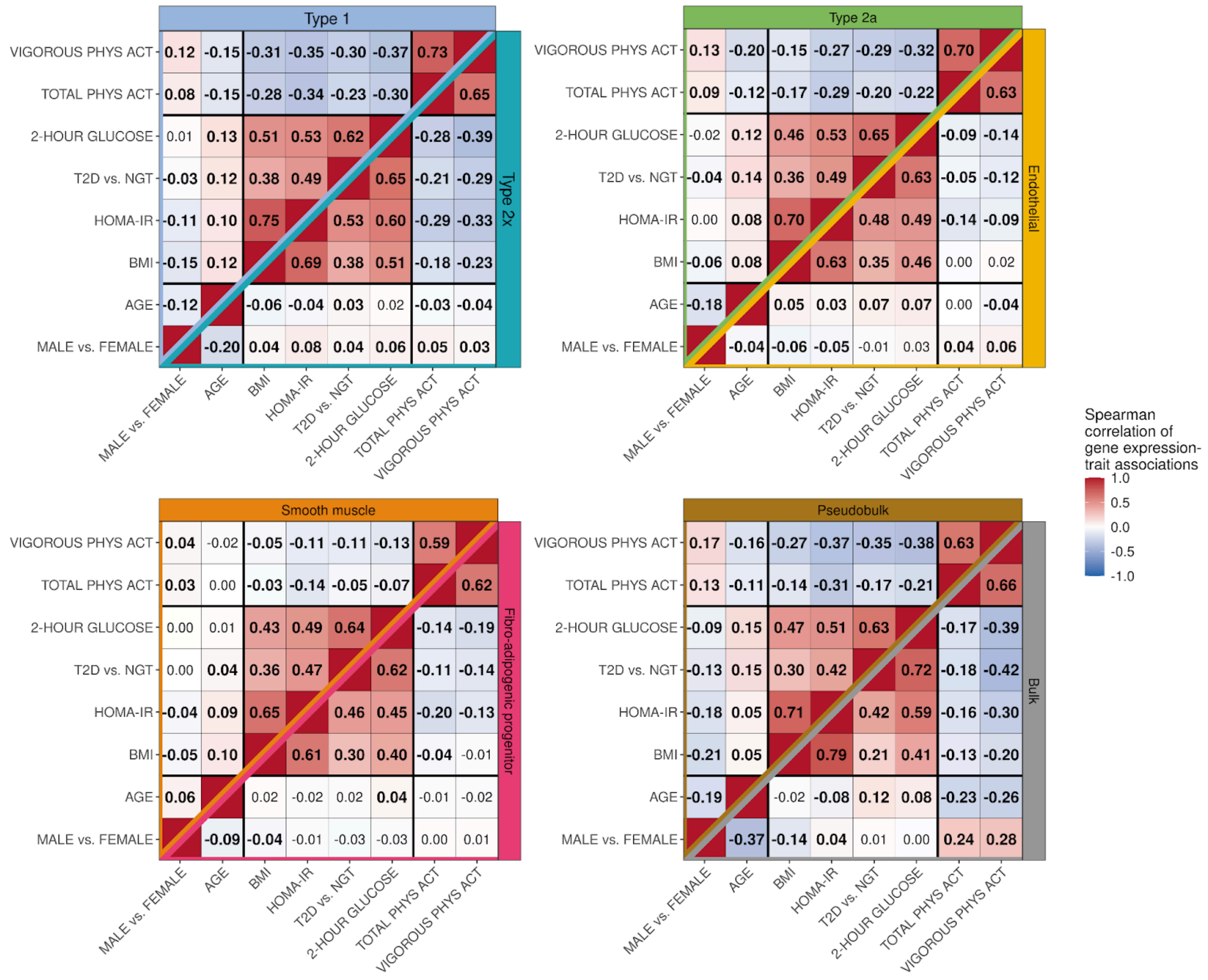


**Figure S14. Heatmap of Spearman correlation of signed (by beta coefficient) −log10 p-values of gene expression–trait associations between traits in cell types, total pseudobulk (pseudobulk), and bulk tissue.** Numbers in bold represent correlations significantly different than 0 tested at a Bonferroni adjusted significance level of p<0.05/224.

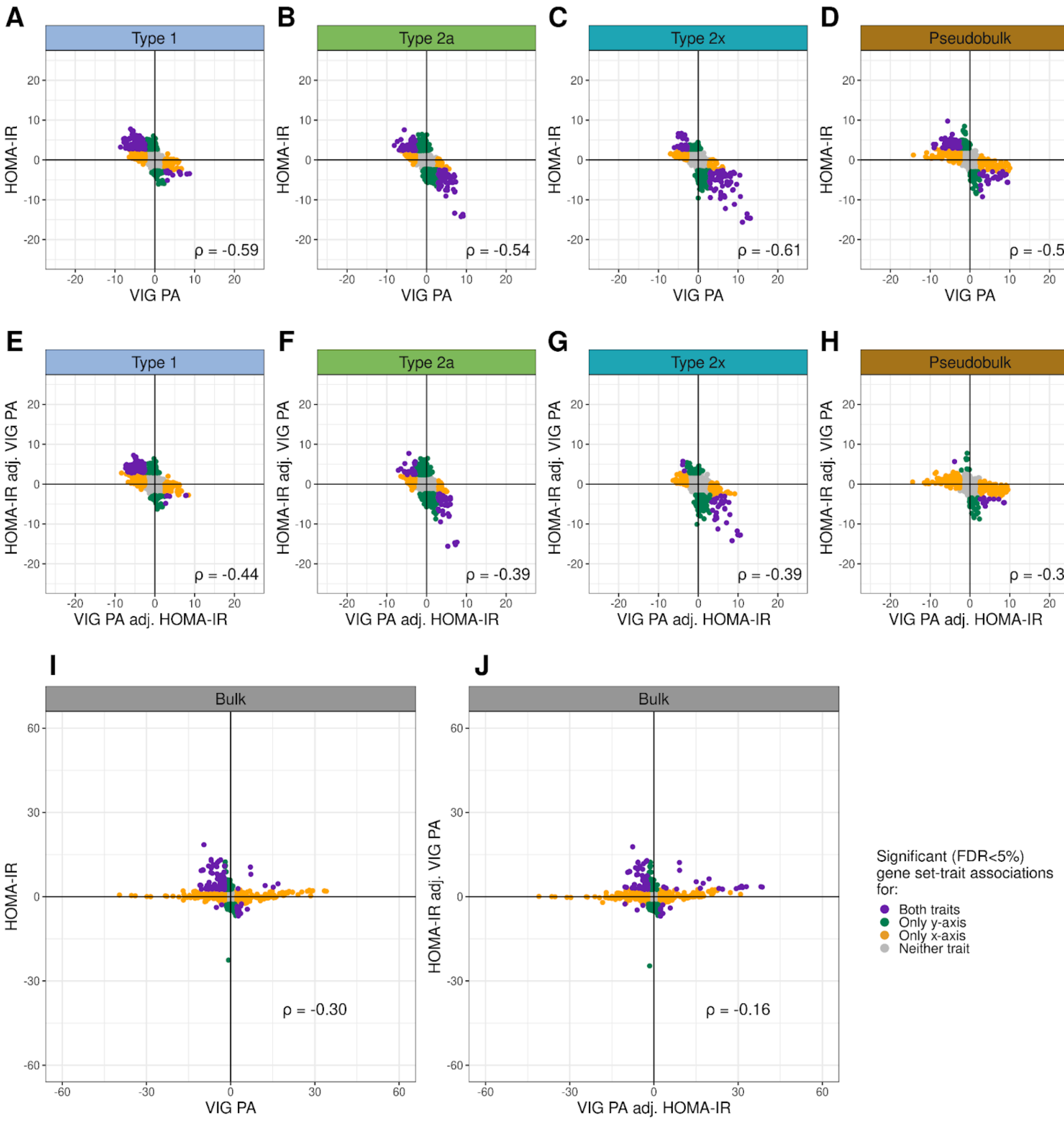


**Figure S15. Scatter plots of Signed −log10 p-value of gene set–trait associations for vigorous physical activity vs. HOMA-IR.** (A-D, I) Signed −log10 p-value of gene set–trait associations for vigorous physical activity (VIG PA) vs. HOMA-IR in models only adjusting for base covariates in muscle fiber types, total pseudobulk (pseudobulk), and bulk tissue. (E-H, J) Signed −log10 p-value of gene set–trait associations for vigorous physical activity vs. HOMA-IR when adjusting (adj.) for both traits in addition to base covariates in the same gene expression model in muscle fiber types, total pseudobulk, and bulk tissue. Each point is a gene set and is colored based on which axis the gene set is significant (FDR<5%) in.

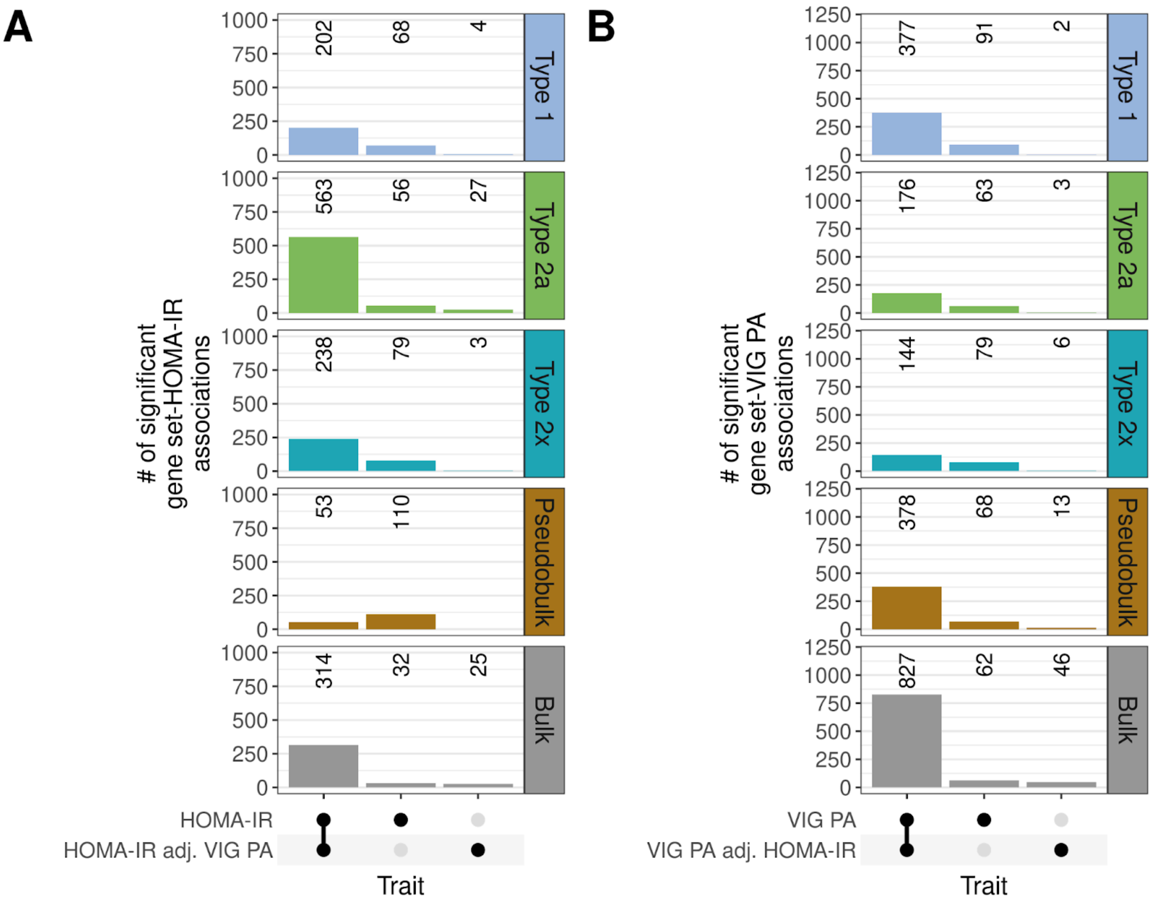


**Figure S16. Upset plot of number of significant gene set–trait associations when adjusting for another trait in the same model.** Upset plot of the number of significant (A) gene set–HOMA-IR and (B) gene set–vigorous physical activity (PA: VIG) associations whether or not adjusting (adj.) for the other trait in addition to base covariates in the same model in each muscle fiber type, total pseudobulk (pseudobulk), and bulk tissue.

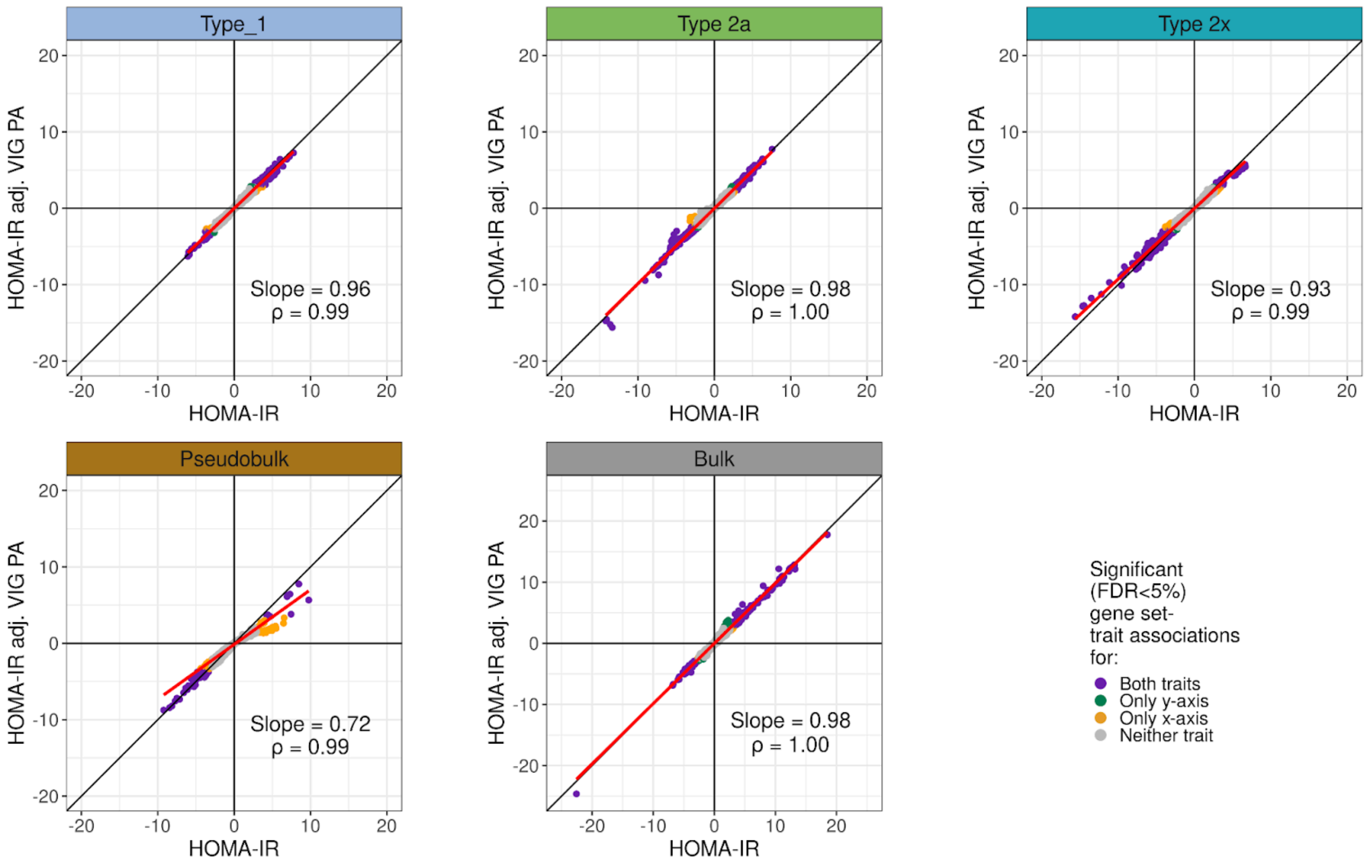


**Figure S17. Scatter plots of signed −log10 p-value of gene set–HOMA-IR associations with and without adjusting (adj.) for vigorous physical activity (VIG PA) in the muscle fiber types, total pseudobulk (pseudobulk), and bulk tissue.** Each point is a gene set and is colored based on whether the gene set is significant (FDR<5%) for HOMA-IR in a model only adjusting for base covariates (x axis) or a model adjusting (adj.) for both HOMA-IR and vigorous physical activity in addition to the base covariates (y axis). The red line represents a simple linear regression line with the slope of the line indicated in each plot.

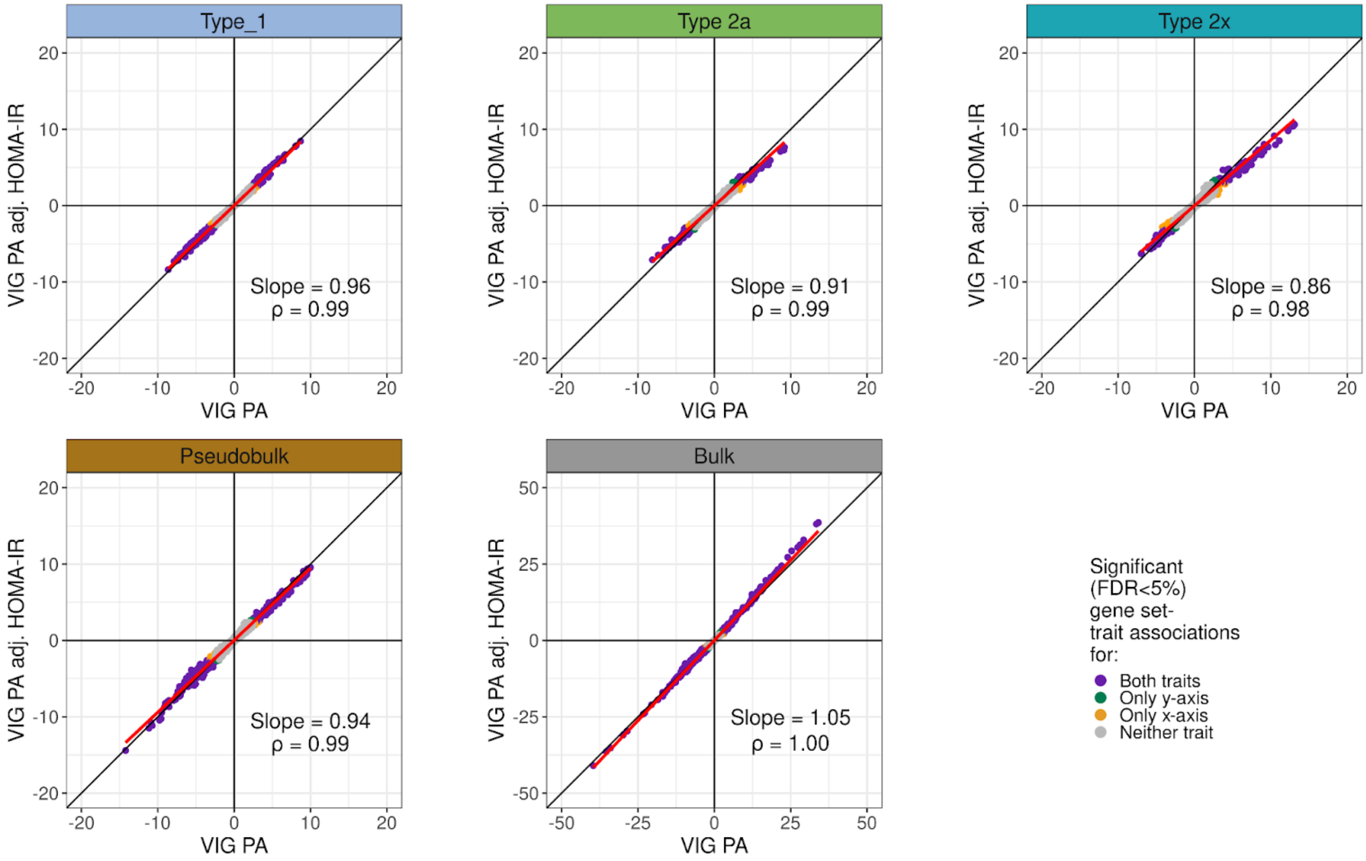


**Figure S18. Scatter plots of signed −log10 p-value of gene set–vigorous physical activity (VIG PA) associations with and without adjusting (adj.) for HOMA-IR in the muscle fiber types, total pseudobulk (pseudobulk), and bulk tissue.** Each point is a gene set and is colored based on whether the gene set is significant (FDR<5%) for vigorous physical activity in a model only adjusting for base covariates (x axis) or a model adjusting (adj.) for both HOMA-IR and vigorous physical activity in addition to the base covariates (y axis). The red line represents a simple linear regression line with the slope of the line indicated in each plot.

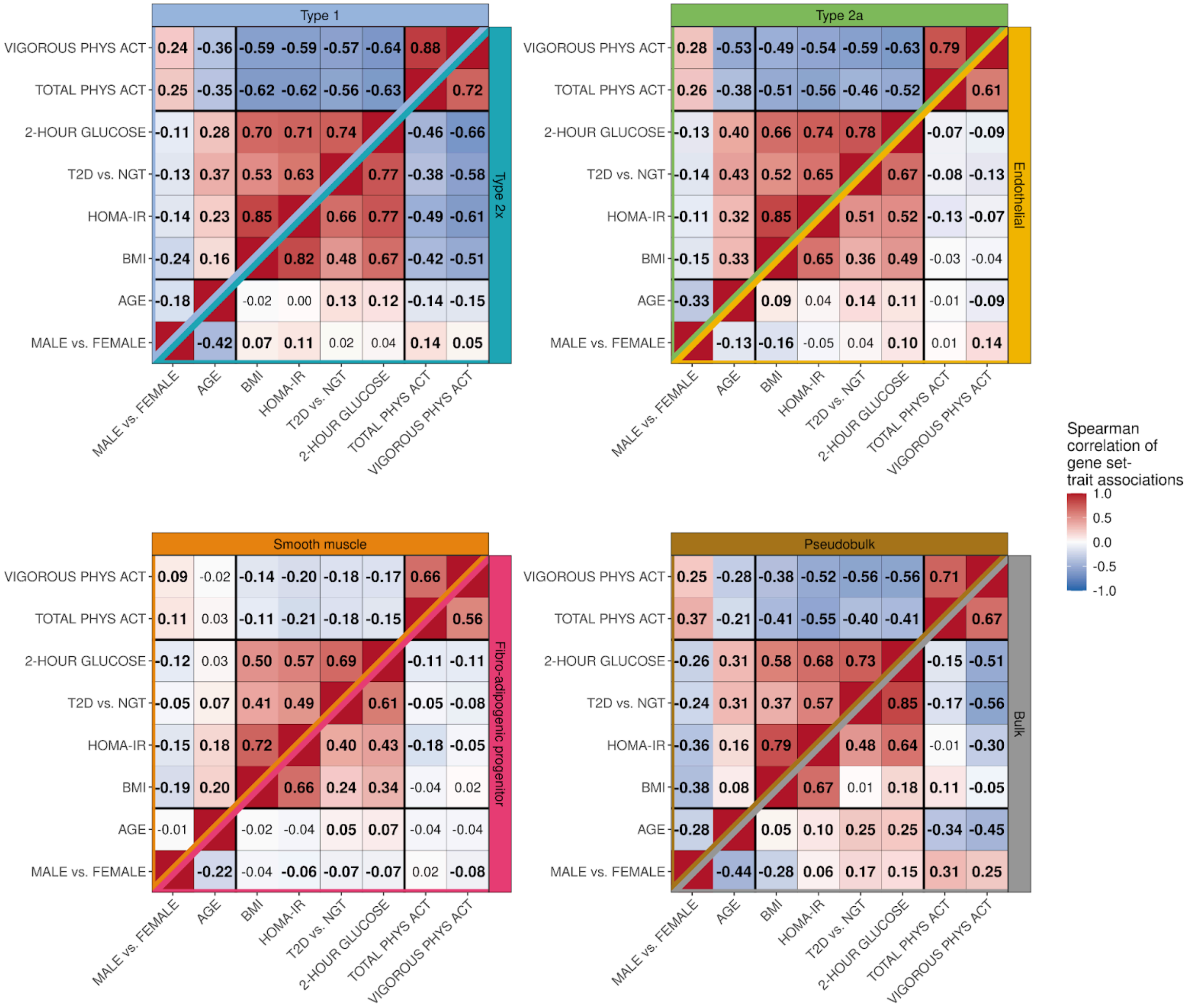


**Figure S19. Heatmap of Spearman correlation of signed (by beta coefficient) −log10 p-values of gene set–trait associations between traits in cell types, total pseudobulk (pseudobulk), and bulk tissue.** Numbers in bold represent correlations significantly different than 0 tested at a Bonferroni adjusted significance level of $p<0.05/224$.

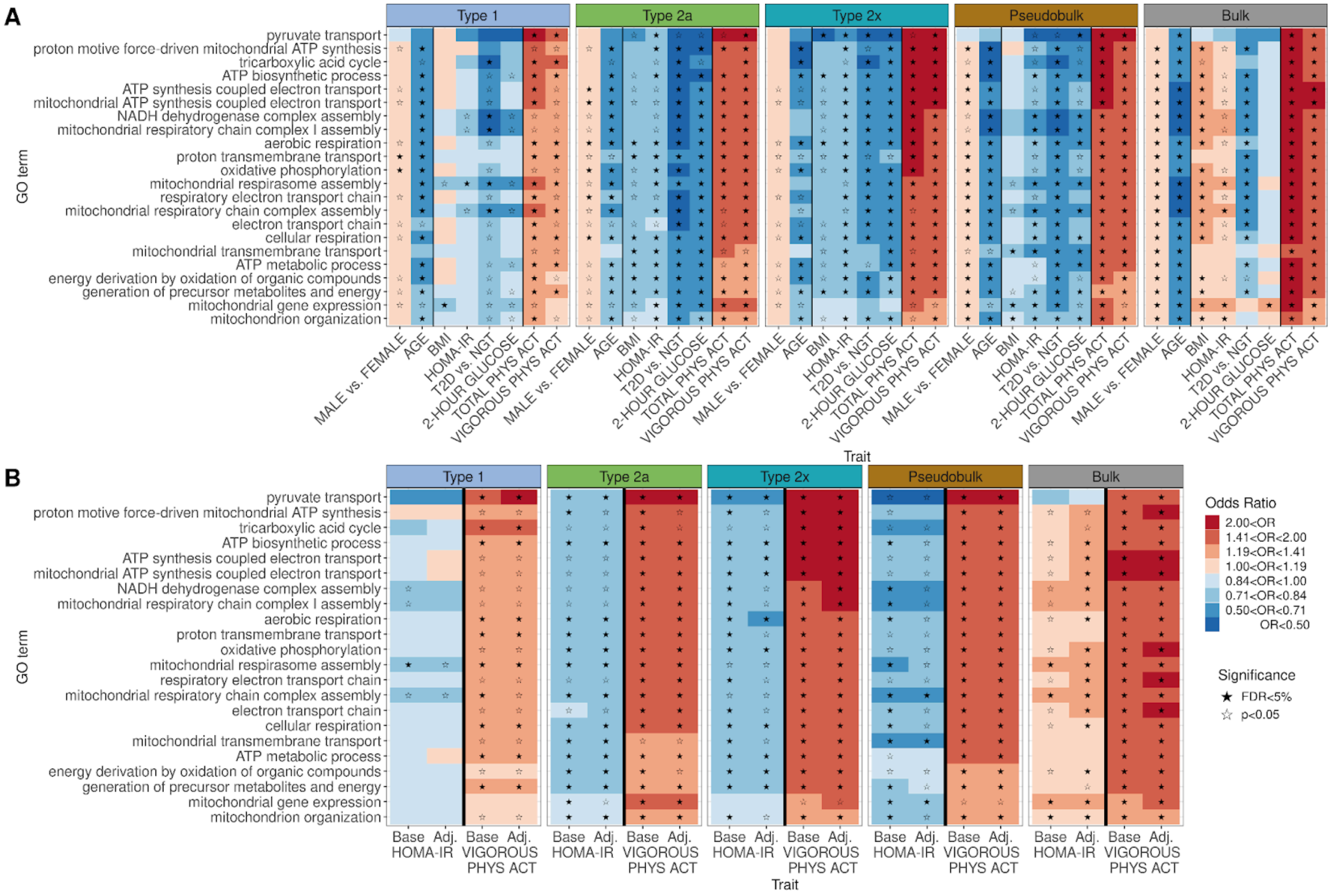


**Figure S20. Examples of significant energy metabolism-related Gene Ontology (GO) terms.** (A) Gene set results of energy metabolism-related GO terms across all traits and muscle fibers, total pseudobulk (pseudobulk), and bulk tissue. A filled-in star indicates that the gene set–trait association is significant (FDR<5%). (B) Gene set–HOMA-IR and gene set–vigorous physical activity associations when adjusting only for base covariates (Base) or when adjusting for the other trait (either vigorous physical activity or HOMA-IR) in addition to base covariates (Adj.).

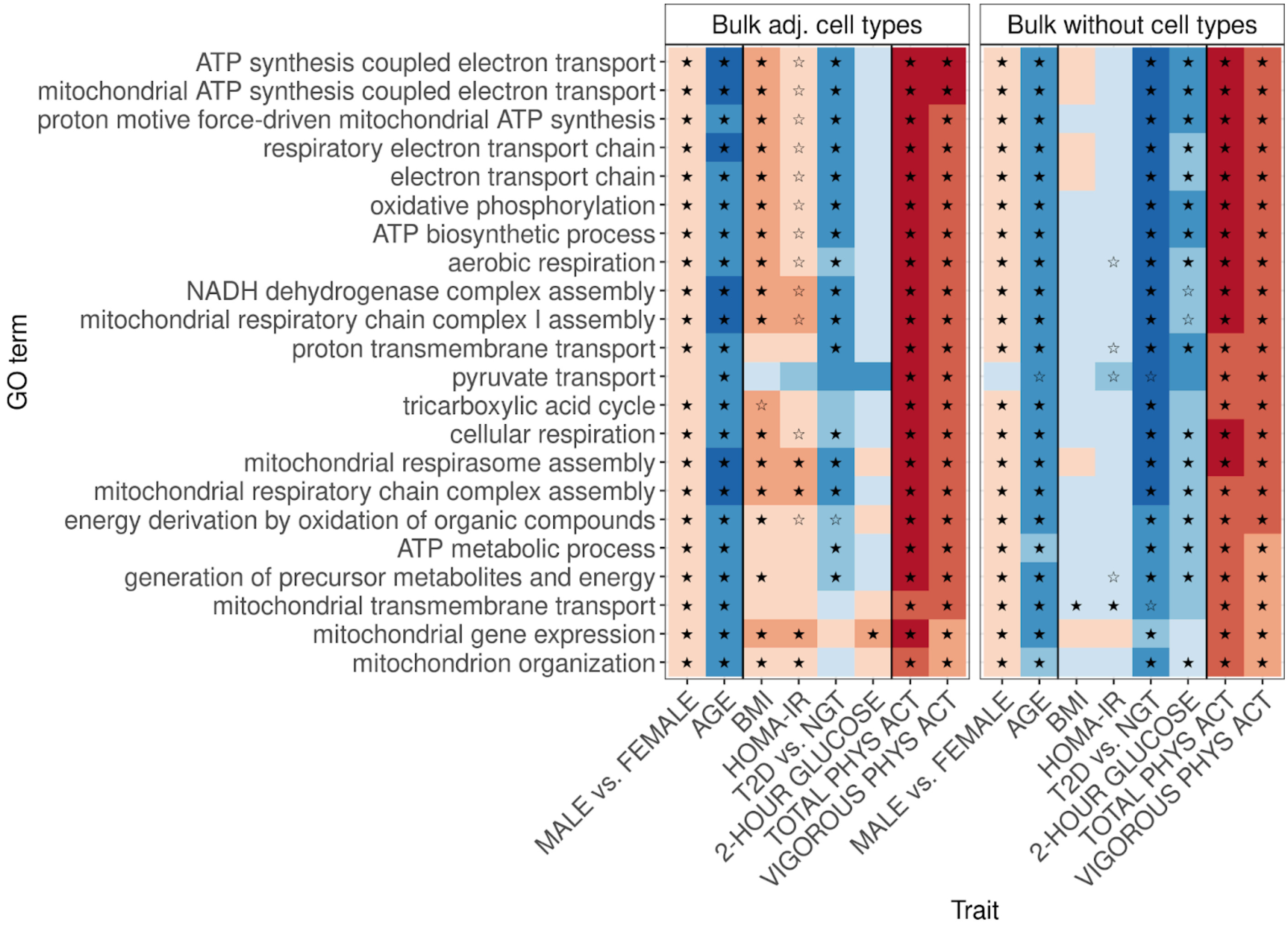


**Figure S21. Examples of significant energy metabolism-related Gene Ontology (GO) terms in bulk tissue (left) when adjusting for observed single nucleus cell type proportions and (right) when not adjusting for cell type proportions.** A filled-in star indicates that the gene set–trait association is significant (FDR<5%). A hollow star indicates that the gene set-trait association had a p-value<0.05.

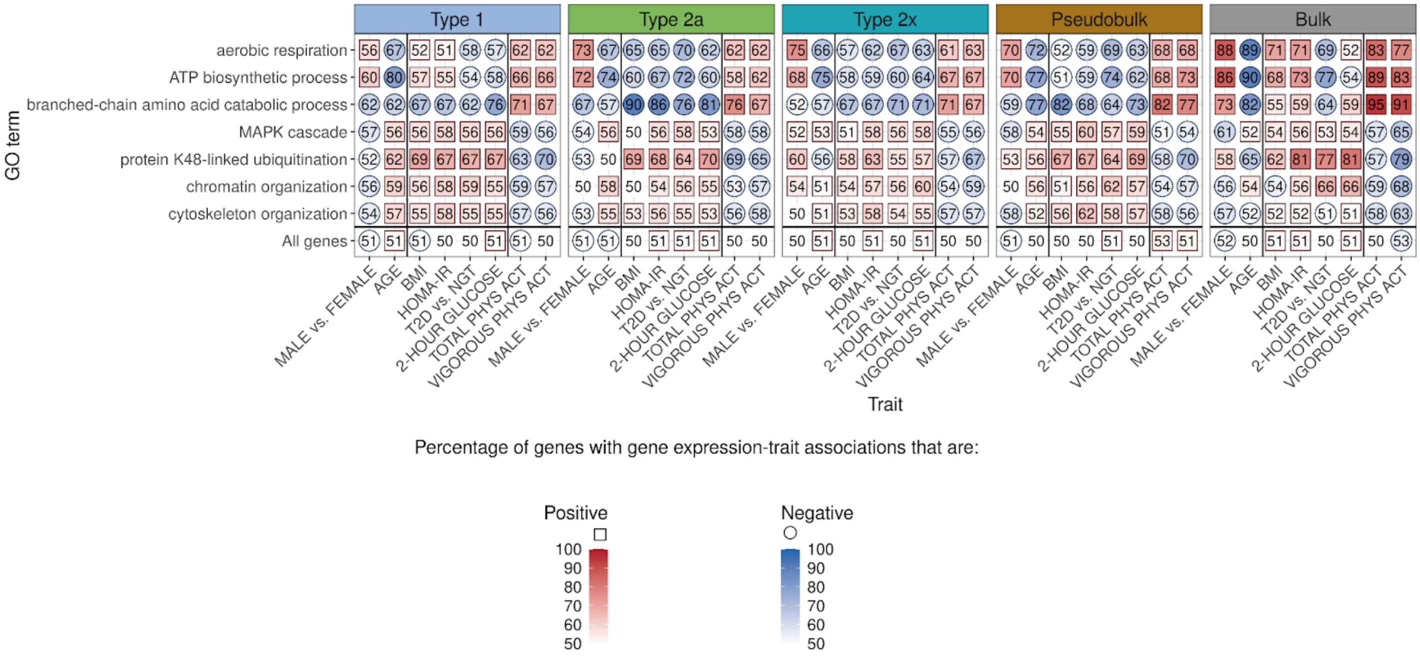


**Figure S22. Percentage of genes with positive (square) or negative (circle) gene expression–trait associations for each gene set across traits and muscle fiber types, total pseudobulk (pseudobulk), and bulk tissue.**

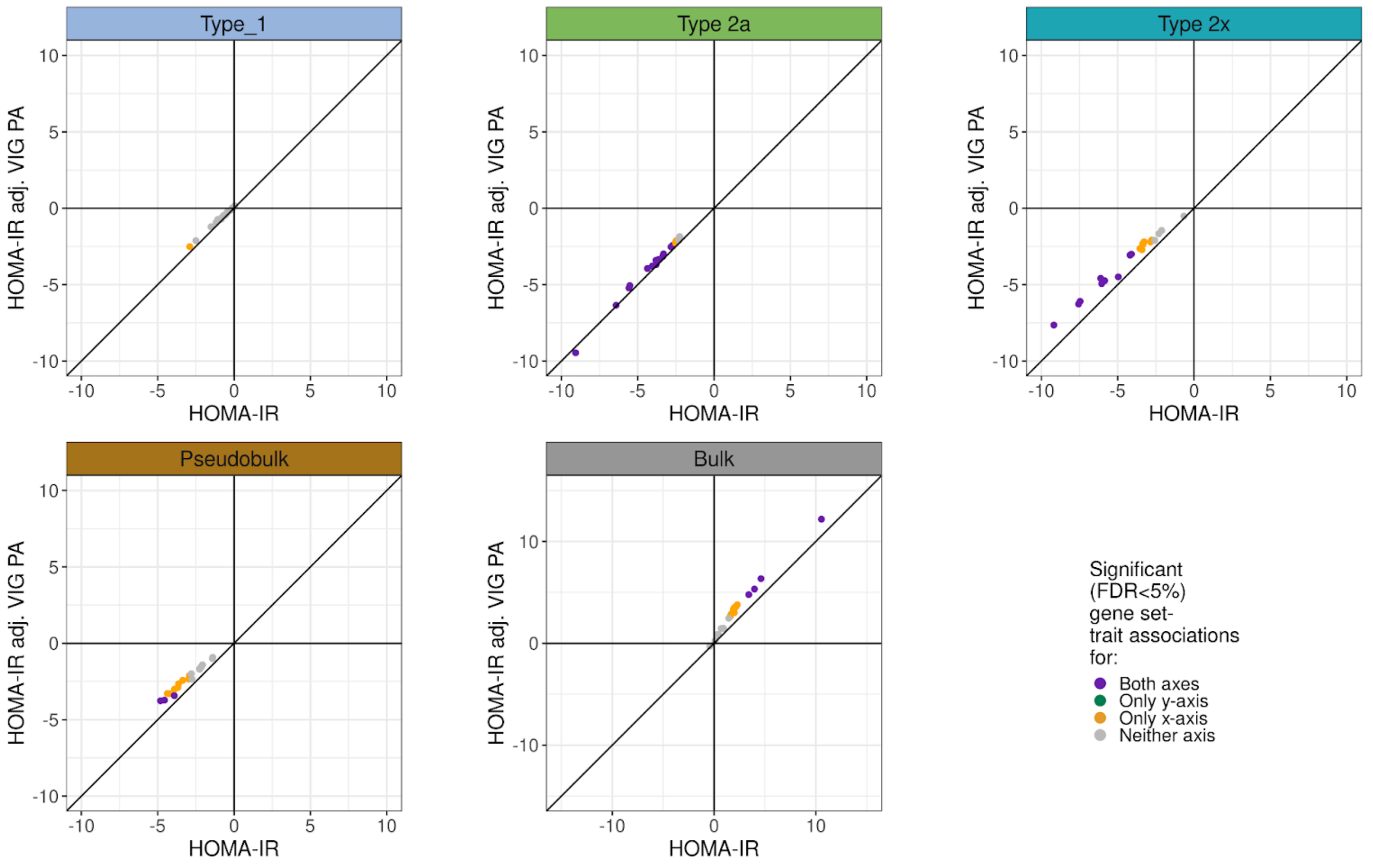


**Figure S23. Scatter plots of signed −log10 p-value of gene set–HOMA-IR associations of energy metabolism-related gene sets with and without adjusting (adj.) for vigorous physical activity (VIG PA) in the muscle fiber types, total pseudobulk (pseudobulk), and bulk tissue.** Each point is an energy metabolism-related gene set and is colored based on whether the gene set is significant (FDR<5%) for HOMA-IR in a model only adjusting for base covariates (x-axis) or a model adjusting (adj.) for both HOMA-IR and vigorous physical activity in addition to the base covariates (y-axis).

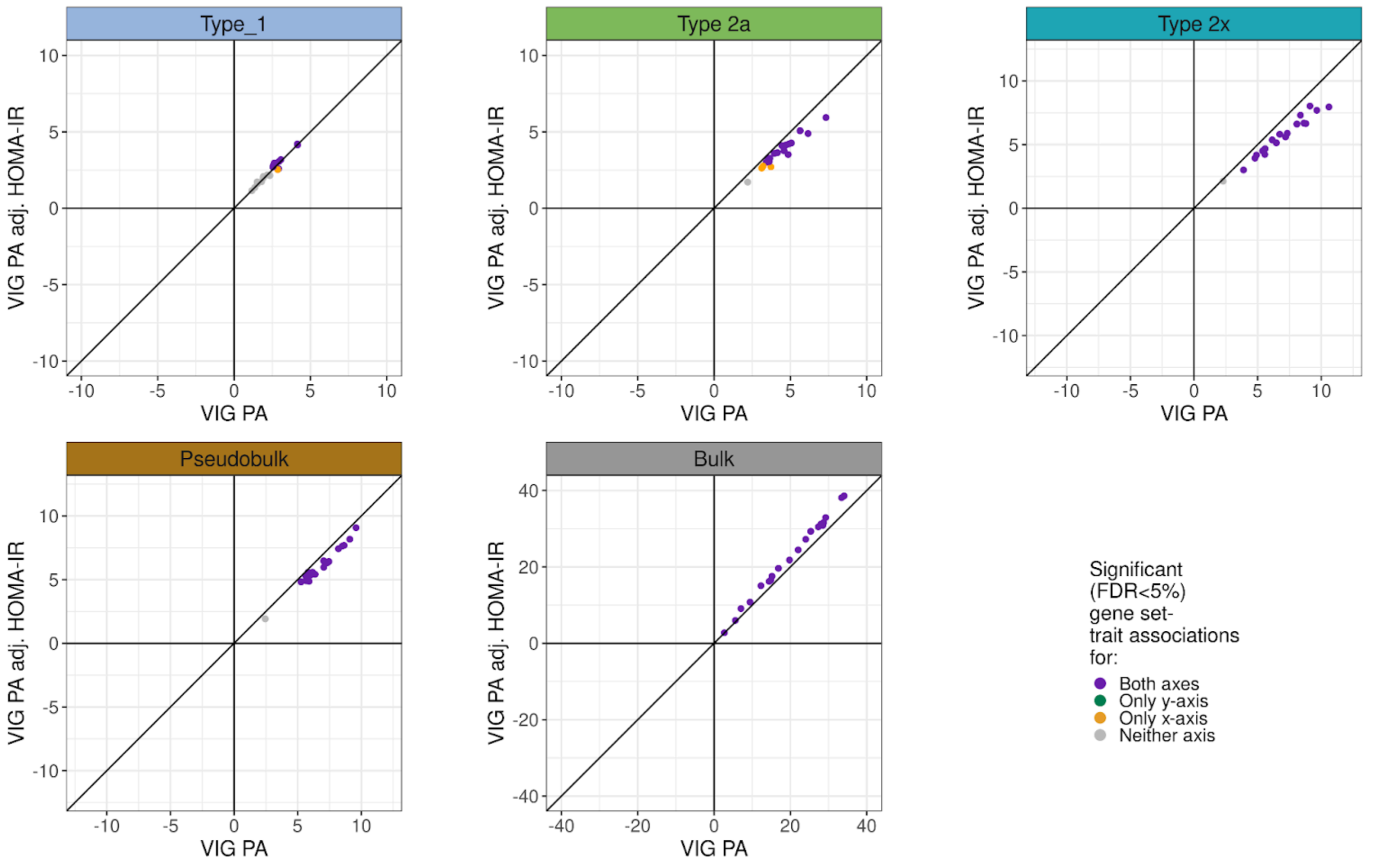


**Figure S24. Scatter plots of signed −log10 p-value of gene set–vigorous physical activity (VIG PA) associations of energy metabolism-related gene sets with and without adjusting (adj.) for HOMA-IR in the muscle fiber types, total pseudobulk (pseudobulk), and bulk tissue.** Each point is an energy metabolism-related gene set and is colored based on whether the gene set is significant (FDR<5%) for vigorous physical activity in a model only adjusting for base covariates (x-axis) or a model adjusting (adj.) for both HOMA-IR and vigorous physical activity in addition to the base covariates (y-axis).

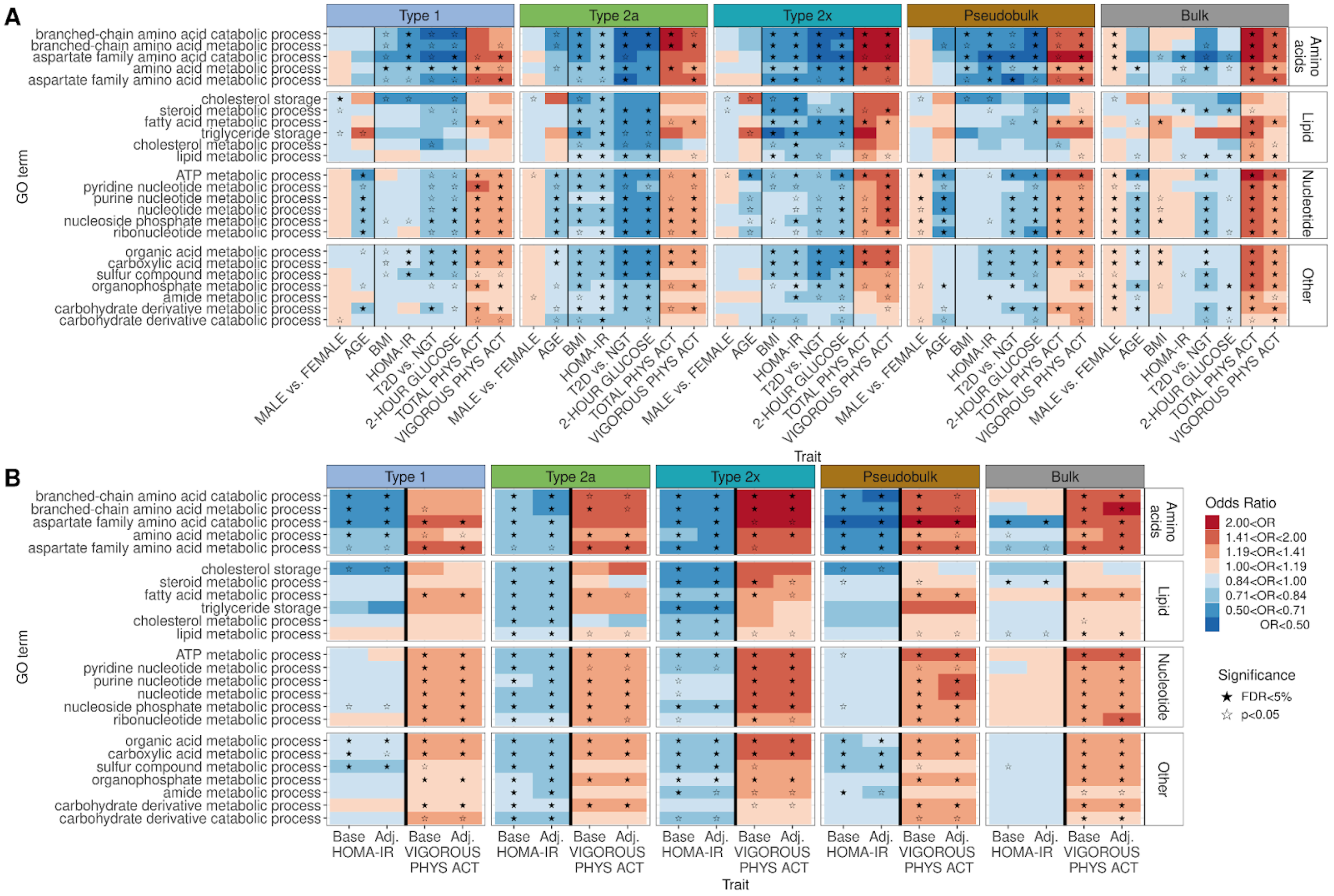


**Figure S25. Examples of significant metabolic and catabolic process-related Gene Ontology (GO) terms.** (A) Gene set results of metabolic and catabolic process-related GO terms across all traits and muscle fibers, total pseudobulk (pseudobulk), and bulk tissue. A filled-in star indicates that the gene set–trait association is significant (FDR<5%). (B) Gene set–HOMA-IR and gene set–vigorous physical activity associations when adjusting only for base covariates (Base) or when adjusting for the other trait (either vigorous physical activity or HOMA-IR) in addition to base covariates (Adj.).

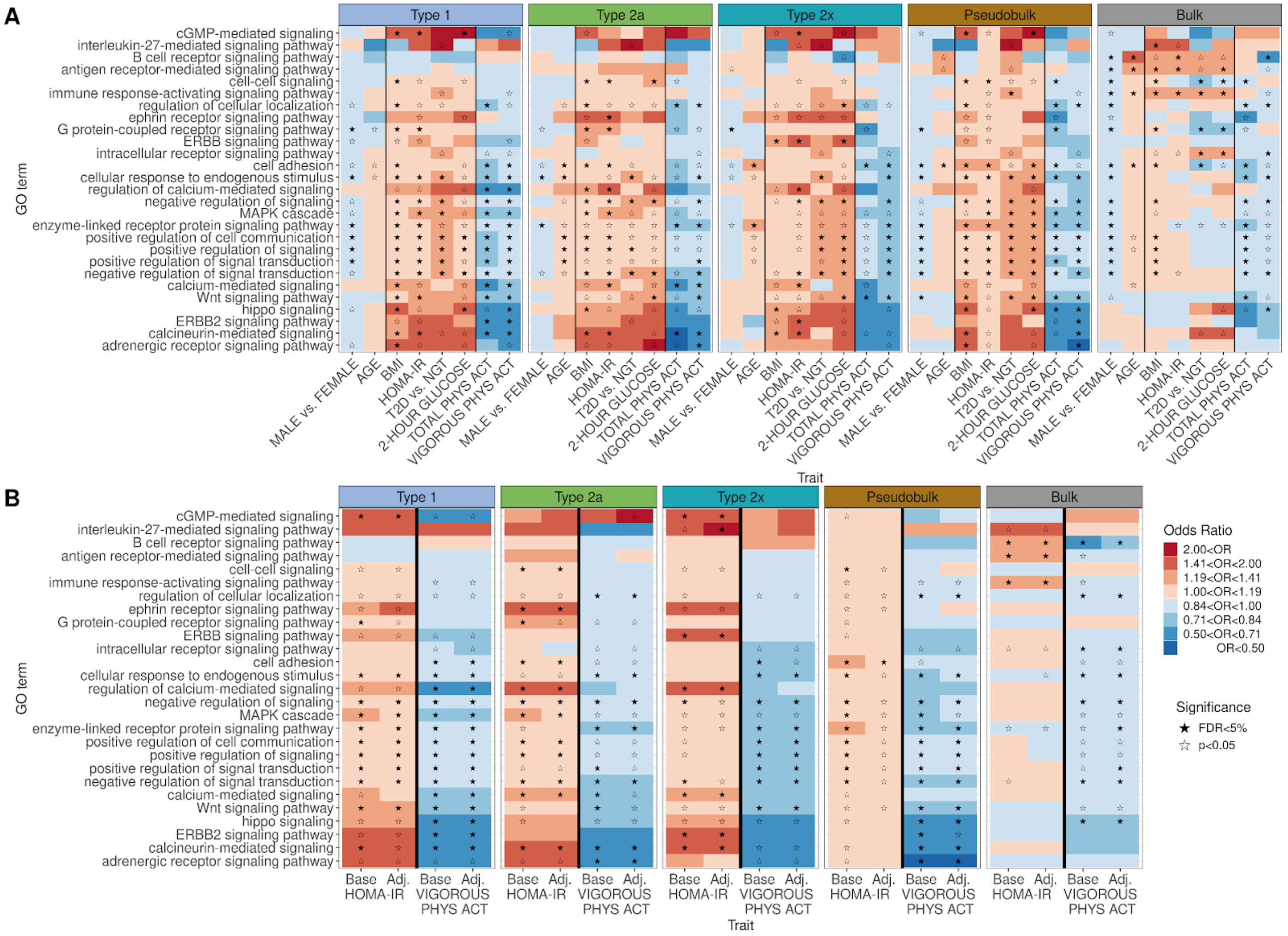


**Figure S26. Examples of significant signaling pathway-related Gene Ontology (GO) terms.** (A) Gene set results of signaling pathway-related GO terms across all traits and muscle fibers, total pseudobulk (pseudobulk), and bulk tissue. A filled-in star indicates that the gene set–trait association is significant (FDR<5%). (B) Gene set–HOMA-IR and gene set–vigorous physical activity associations when adjusting only for base covariates (Base) or when adjusting for the other trait (either vigorous physical activity or HOMA-IR) in addition to base covariates (Adj.).

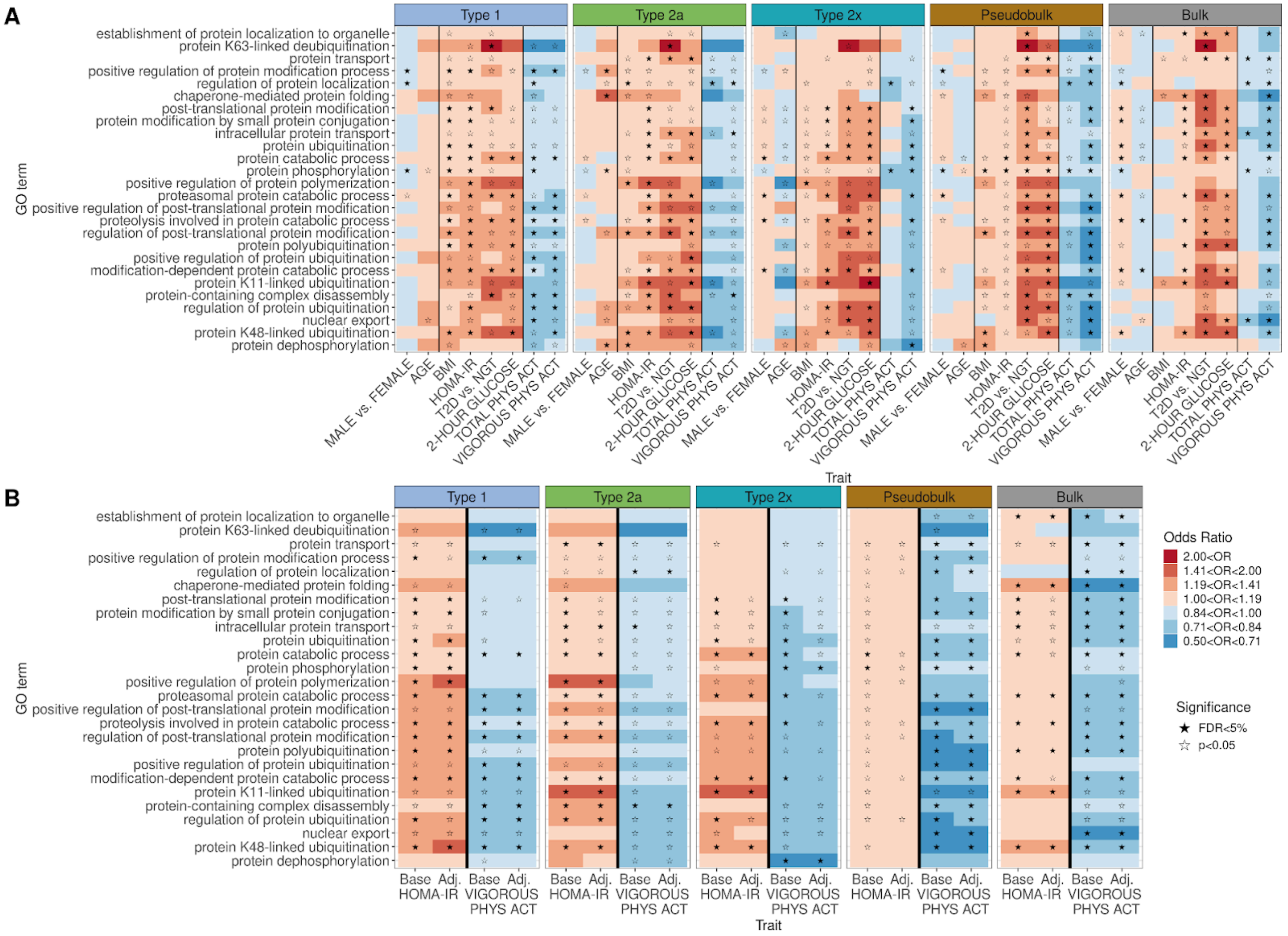


**Figure S27. Examples of significant protein function and modification-related Gene Ontology (GO) terms.** (A) Gene set results of protein function modification-related GO terms across all traits and muscle fibers, total pseudobulk (pseudobulk), and bulk tissue. A filled-in star indicates that the gene set–trait association is significant (FDR<5%). (B) Gene set–HOMA-IR and gene set–vigorous physical activity associations when adjusting only for base covariates (Base) or when adjusting for the other trait (either vigorous physical activity or HOMA-IR) in addition to base covariates (Adj.).

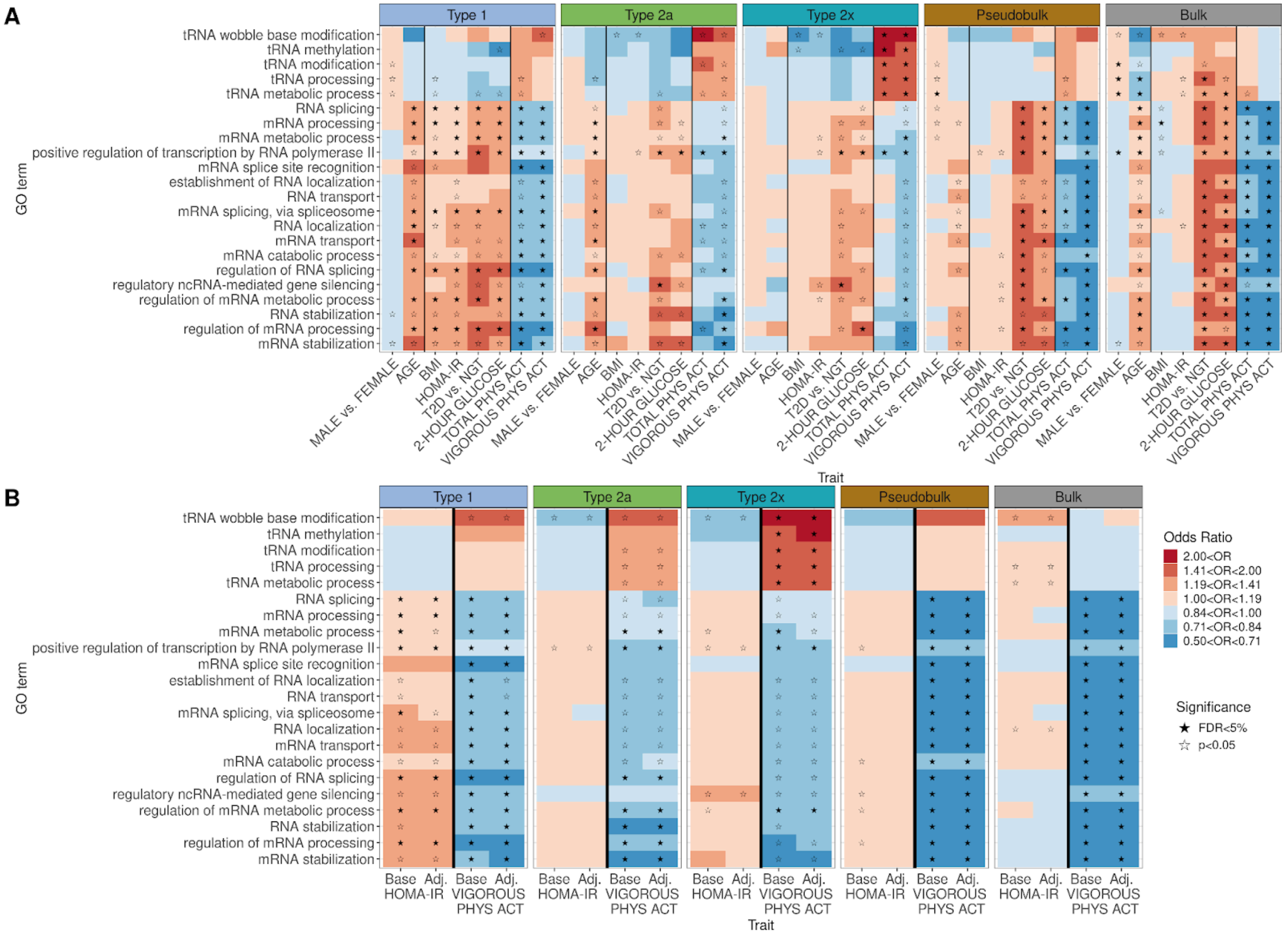


**Figure S28. Examples of significant RNA and mRNA function-related Gene Ontology (GO) terms.** (A) Gene set results of RNA and mRNA function-related GO terms across all traits and muscle fibers, total pseudobulk (pseudobulk), and bulk tissue. A filled-in star indicates that the gene set–trait association is significant (FDR<5%). (B) Gene set–HOMA-IR and gene set–vigorous physical activity associations when adjusting only for base covariates (Base) or when adjusting for the other trait (either vigorous physical activity or HOMA-IR) in addition to base covariates (Adj.).

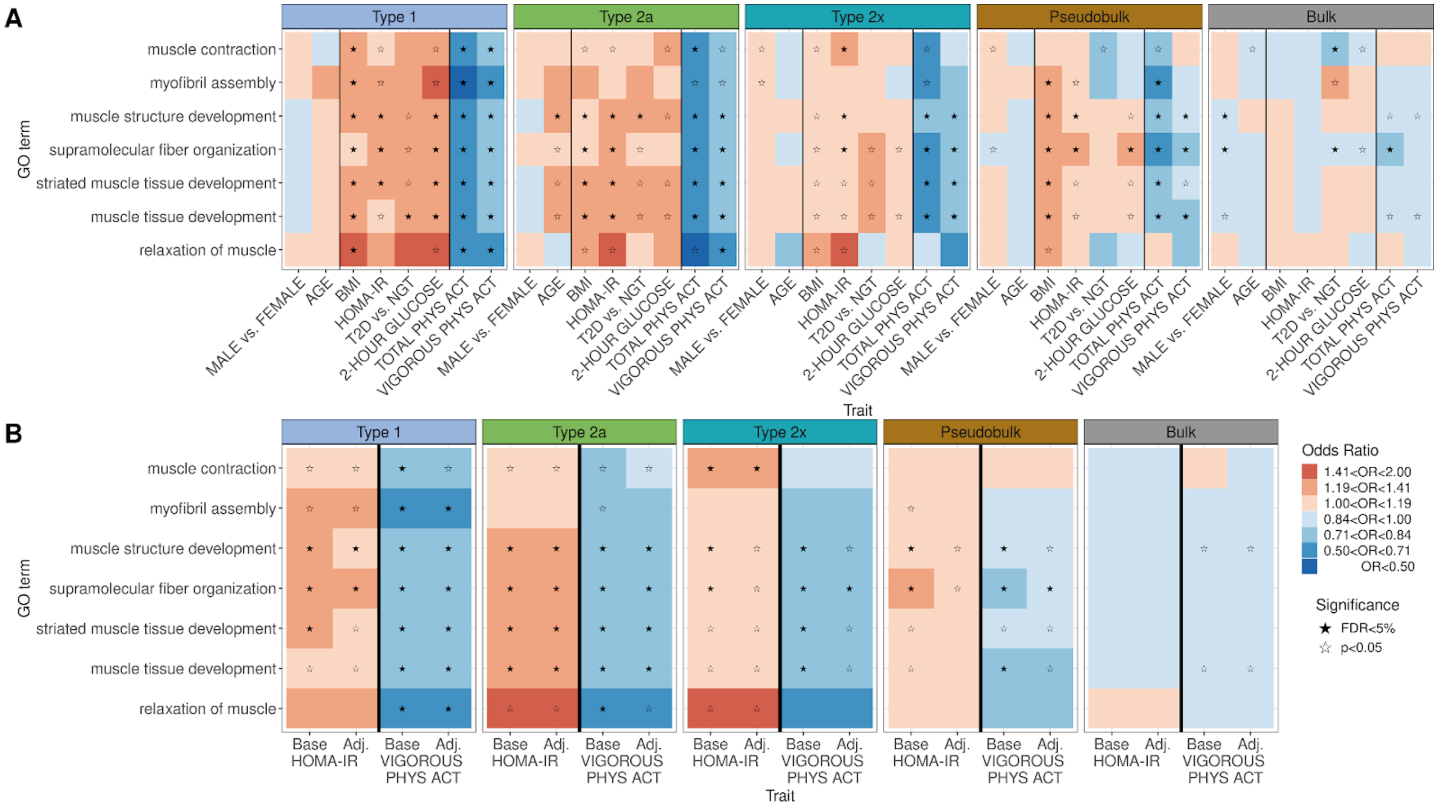


**Figure S29. Examples of significant muscle function and development-related Gene Ontology (GO) terms.** (A) Gene set results of muscle function and development-related GO terms across all traits and muscle fibers, total pseudobulk (pseudobulk), and bulk tissue. A filled-in star indicates that the gene set–trait association is significant (FDR<5%). (B) Gene set–HOMA-IR and gene set–vigorous physical activity associations when adjusting only for base covariates (Base) or when adjusting for the other trait (either vigorous physical activity or HOMA-IR) in addition to base covariates (Adj.).

**Table S1. Sample characteristics of 263 vastus lateralis biopsy donors from the FUSION Tissue Biopsy Study.** Each individual's glycemic status was characterized as normal glucose tolerance (NGT), impaired fasting glucose (IFG), impaired glucose tolerance (IGT), and type 2 diabetes (T2D) using an oral glucose tolerance test (OGTT). Characteristics are shown for females and males separately and for the total sample.

| | | Female (n=113) | Male (n=150) | Total (n=263) |
|---|---|---|---|---|
| Age (years) | mean (SD) | 60.9 (7.1) | 59.8 (7.9) | 60.3 (7.6) |
| | [Range] | [33–76] | [25–79] | [25–79] |
| BMI (kg/m²) | mean (SD) | 27.4 (4.1) | 27.9 (4.3) | 27.7 (4.2) |
| 2-hour glucose (mmol/L) | mean (SD) | 7.9 (2.5) | 8.0 (2.9) | 7.9 (2.8) |
| | n (%) missing | 0 (0%) | 1 (0.7%) | 1 (0.4%) |
| HOMA-IR | mean (SD) | 2.2 (1.4) | 2.7 (2.0) | 2.5 (1.8) |
| | n ≤ 2.9 (%) | 88 (77.9%) | 99 (66.0%) | 187 (71.1%) |
| | n > 2.9 (%) | 25 (22.1%) | 51 (34.0%) | 76 (28.9%) |
| OGTT status | (n (%)) | | | |
| | NGT | 51 (45.1%) | 43 (28.7%) | 94 (35.7%) |
| | IFG | 11 (9.7%) | 23 (15.3%) | 34 (12.9%) |
| | IGT | 29 (25.7%) | 36 (24.0%) | 65 (24.7%) |
| | T2D | 22 (19.5%) | 48 (32.0%) | 70 (26.6%) |
| Total physical activity | mean (SD) | 2366.5 (1545.6) | 2750.3 (1985.2) | 2586.2 (1817.1) |
| (METS*kg*hr/wk) | n (%) missing | 1 (0.9%) | 0 (0%) | 1 (0.4%) |
| Vigorous physical activity | mean (SD) | 737.5 (872.3) | 1271.0 (1442.6) | 1042.9 (1257.6) |
| (METS*kg*hr/wk) | n (%) missing | 1 (0.9%) | 0 (0%) | 1 (0.4%) |

**Table S2. Mean and standard deviation of cell type proportions across individuals.** For each cell type, each individual's cell type proportion was calculated as the number of nuclei of that cell type divided by their total number of nuclei (combining snATAC-seq and snRNA-seq nuclei) (n=263).

| Cell type | Mean proportion per individual (SD) |
|---|---|
| Adipocyte | 0.01 (0.01) |
| Endothelial cell | 0.10 (0.03) |
| Fibro-adipogenic progenitor cell | 0.05 (0.03) |
| Macrophage | 0.01 (0.01) |
| Neuromuscular junction (muscle fiber) | 0.02 (0.02) |
| Neuronal cell | 0.02 (0.01) |
| Satellite cell | 0.01 (0.01) |
| Smooth muscle cell | 0.04 (0.02) |
| T cell | 0.03 (0.01) |
| Type 1 muscle fiber | 0.34 (0.11) |
| Type 2a muscle fiber | 0.20 (0.07) |
| Type 2x muscle fiber | 0.16 (0.08) |

**Table S3 (separate file). Differential cell type abundance by trait results.** Cell type, trait of interest, 2nd trait adjusted for, Log2 Fold Change, standard error, statistic, FDR, sample size, and modality. NAs for "2nd trait adjusted" indicates that no additional trait was adjusted for. Rows with an FDR<5% are considered significant. Modality indicates whether snRNA, snATAC, or both nuclei were used in the analysis. For detailed information on the models and how FDR was calculated please consult the method section.

**Table S4. Number of significant (FDR<5%) genes per trait across cell types.** Number (#) of significant (FDR<5%) genes and percentage (%) of significant genes of all genes tested. Results were from models adjusting for base covariates and only 1 trait of interest. Values specified by NA indicate that less than 25 samples were available with that trait in that cell type and no models were run for these trait-cell type combinations.

| Trait of interest | Endo-thelial cell | | Fibro-adipo-genic progenitor cell | | Macro-phage | | Neuro-muscular junction (muscle fiber) | | Neuronal cell | | Satellite cell | | Smooth muscle cell | | Type 1 muscle fiber | | Type 2a muscle fiber | | Type 2x muscle fiber | | Total pseudobulk | | Bulk | |
|---|---|---|---|---|---|---|---|---|---|---|---|---|---|---|---|---|---|---|---|---|---|---|---|---|
| | # | % | # | % | # | % | # | % | # | % | # | % | # | % | # | % | # | % | # | % | # | % | # | % |
| AGE | 0 | 0.0 | 4 | 0.0 | 0 | 0.0 | 0 | 0.0 | 1 | 0.0 | 0 | 0.0 | 1 | 0.0 | 53 | 0.2 | 192 | 1.0 | 107 | 0.6 | 56 | 0.3 | 1275 | 5.7 |
| BMI | 0 | 0.0 | 6 | 0.0 | 0 | 0.0 | 0 | 0.0 | 0 | 0.0 | 0 | 0.0 | 0 | 0.0 | 963 | 4.0 | 406 | 2.1 | 257 | 1.5 | 684 | 3.7 | 1377 | 6.2 |
| 2-HOUR GLUCOSE | 1 | 0.0 | 1 | 0.0 | 0 | 0.0 | 0 | 0.0 | 1 | 0.0 | 0 | 0.0 | 0 | 0.0 | 0 | 0.0 | 20 | 0.1 | 52 | 0.3 | 4 | 0.0 | 19 | 0.1 |
| HOMA-IR | 0 | 0.0 | 3 | 0.0 | 0 | 0.0 | 0 | 0.0 | 0 | 0.0 | 0 | 0.0 | 0 | 0.0 | 413 | 1.7 | 366 | 1.9 | 260 | 1.5 | 358 | 1.9 | 1955 | 8.8 |
| TOTAL PHYS ACT | 0 | 0.0 | 0 | 0.0 | 0 | 0.0 | 0 | 0.0 | 3 | 0.0 | 0 | 0.0 | 0 | 0.0 | 87 | 0.4 | 0 | 0.0 | 0 | 0.0 | 0 | 0.0 | 14 | 0.1 |
| VIGOROUS PHYS ACT | 0 | 0.0 | 0 | 0.0 | 0 | 0.0 | 0 | 0.0 | 0 | 0.0 | 0 | 0.0 | 0 | 0.0 | 403 | 1.7 | 151 | 0.8 | 22 | 0.1 | 83 | 0.4 | 354 | 1.6 |
| MALE vs. FEMALE | 59 | 0.4 | 379 | 2.6 | 6 | 0.1 | 114 | 0.9 | 0 | 0.0 | 156 | 1.4 | 36 | 0.3 | 3117 | 13.1 | 2499 | 13.0 | 1974 | 11.3 | 2861 | 15.4 | 7192 | 32.2 |
| T2D vs. NGT | 1 | 0.0 | 0 | 0.0 | NA | NA | 0 | 0.0 | NA | NA | 0 | 0.0 | 0 | 0.0 | 0 | 0.0 | 1 | 0.0 | 10 | 0.1 | 2 | 0.0 | 0 | 0.0 |

**Table S5 (separate file). Gene expression-trait associations across cell types, total pseudobulk, and bulk tissue.** Gene, mean gene count, Log2 Fold Change, Log2 Fold Change Standard error, Statistic, P-value, FDR, Cell type, Trait of interest, 2nd trait adjusted for, sample size. NAs for the 2nd trait adjusted for indicates that no additional trait was adjusted for. Rows with an FDR<5% are significant. For detailed information on the models and how FDR was calculated please consult the method section.

**Table S6. Number of significant (FDR<5%) genes per trait for Type 1 sample size and UMI count downsampled to the matched cell type.** Values specified by NA indicate that less than 25 samples were available with that trait in that cell type and no models were run for these trait-cell type combinations.

| Trait of interest | Endothelial cell | Fibro-adipogenic progenitor cell | Macro-phage | Neuro-muscular junction (muscle fiber) | Neuronal cell | Satellite cell | Smooth muscle cell | Type 2a muscle fiber | Type 2x muscle fiber |
|---|---|---|---|---|---|---|---|---|---|
| AGE | 7 | 8 | 0 | 0 | 0 | 0 | 3 | 19 | 9 |
| BMI | 233 | 211 | 1 | 49 | 0 | 0 | 140 | 889 | 523 |
| 2-HOUR GLUCOSE | 8 | 0 | 1 | 4 | 0 | 0 | 4 | 14 | 0 |
| HOMA-IR | 88 | 94 | 0 | 11 | 0 | 0 | 70 | 465 | 283 |
| TOTAL PHYS ACT | 7 | 14 | 0 | 0 | 0 | 3 | 1 | 31 | 0 |
| VIGOROUS PHYS ACT | 67 | 61 | 0 | 0 | 0 | 1 | 38 | 301 | 221 |
| MALE vs. FEMALE | 561 | 596 | 20 | 327 | 5 | 103 | 394 | 2043 | 1244 |
| T2D vs. NGT | 1 | 0 | *NA* | 0 | *NA* | 0 | 1 | 0 | 0 |

**Table S7. Number of significant (FDR<5%) genes in both cell types by trait.** Total number (T) of significant genes in both cell type 1 and cell type 2 for the given trait and percentage of these significant genes that have a concordant (C) or discordant (D) direction of association for the given trait.

| Cell type 1 | Cell type 2 | HOMA-IR | | | VIGOROUS PHYS ACT | | |
|---|---|---|---|---|---|---|---|
| | | C (%) | D (%) | T (#) | C (%) | D (%) | T (#) |
| Type 2a | Type 1 | 100 | 0 | 112 | 100 | 0 | 79 |
| Type 2x | Type 1 | 100 | 0 | 69 | 100 | 0 | 9 |
| Type 2x | Type 2a | 100 | 0 | 133 | 100 | 0 | 12 |
| Total pseudobulk | Type 1 | 100 | 0 | 180 | 98 | 2 | 65 |
| Total pseudobulk | Type 2a | 100 | 0 | 136 | 100 | 0 | 36 |
| Total pseudobulk | Type 2x | 100 | 0 | 91 | 100 | 0 | 9 |
| Total pseudobulk | Bulk | 99 | 1 | 115 | 100 | 0 | 17 |
| Bulk | Type 1 | 99 | 1 | 116 | 100 | 0 | 36 |
| Bulk | Type 2a | 98 | 2 | 120 | 100 | 0 | 17 |
| Bulk | Type 2x | 100 | 0 | 83 | 100 | 0 | 4 |

**Table S8. Number of significant (FDR<5%) genes for both traits across muscle fiber types, total pseudobulk, and bulk tissue.** Total number (T) of significant genes for both trait 1 and trait 2 from results with models adjusting for only base covariates and percentage of these significant genes that have a concordant (C) or discordant (D) direction of association for these two traits.

| Trait 1 | Trait 2 | Type 1 | | | Type 2a | | | Type 2x | | | Total pseudobulk | | | Bulk | | |
|---|---|---|---|---|---|---|---|---|---|---|---|---|---|---|---|---|
| | | C (%) | D (%) | T (#) | C (%) | D (%) | T (#) | C (%) | D (%) | T (#) | C (%) | D (%) | T (#) | C (%) | D (%) | T (#) |
| AGE | MALE vs. FEMALE | 30 | 70 | 27 | 11 | 89 | 89 | 8 | 92 | 61 | 17 | 83 | 24 | 7 | 93 | 765 |
| BMI | MALE vs. FEMALE | 21 | 79 | 280 | 45 | 55 | 114 | 68 | 32 | 98 | 18 | 82 | 200 | 32 | 68 | 490 |
| BMI | AGE | 57 | 43 | 7 | 59 | 41 | 17 | 0 | 100 | 6 | 83 | 17 | 6 | 38 | 62 | 82 |
| HOMA-IR | MALE vs. FEMALE | 36 | 64 | 131 | 67 | 33 | 123 | 74 | 26 | 85 | 21 | 79 | 102 | 55 | 45 | 671 |
| HOMA-IR | AGE | 0 | 100 | 1 | 55 | 45 | 11 | 50 | 50 | 4 | NA | NA | 0 | 27 | 73 | 135 |
| HOMA-IR | BMI | 100 | 0 | 275 | 100 | 0 | 174 | 100 | 0 | 110 | 100 | 0 | 180 | 100 | 0 | 823 |
| T2D vs. NGT | MALE vs. FEMALE | NA | NA | 0 | NA | NA | 0 | 0 | 100 | 1 | 0 | 100 | 1 | NA | NA | 0 |
| T2D vs. NGT | AGE | NA | NA | 0 | NA | NA | 0 | NA | NA | 0 | NA | NA | 0 | NA | NA | 0 |
| T2D vs. NGT | BMI | NA | NA | 0 | NA | NA | 0 | 100 | 0 | 2 | NA | NA | 0 | NA | NA | 0 |
| T2D vs. NGT | HOMA-IR | NA | NA | 0 | 100 | 0 | 1 | 100 | 0 | 4 | NA | NA | 0 | NA | NA | 0 |
| 2-HOUR GLUCOSE | MALE vs. FEMALE | NA | NA | 0 | 36 | 64 | 11 | 50 | 50 | 22 | 50 | 50 | 2 | 20 | 80 | 5 |
| 2-HOUR GLUCOSE | AGE | NA | NA | 0 | 100 | 0 | 2 | 100 | 0 | 1 | NA | NA | 0 | 100 | 0 | 1 |
| 2-HOUR GLUCOSE | BMI | NA | NA | 0 | 100 | 0 | 5 | 100 | 0 | 25 | 100 | 0 | 2 | 100 | 0 | 8 |
| 2-HOUR GLUCOSE | HOMA-IR | NA | NA | 0 | 100 | 0 | 8 | 100 | 0 | 29 | 100 | 0 | 2 | 100 | 0 | 12 |
| 2-HOUR GLUCOSE | T2D vs. NGT | NA | NA | 0 | 100 | 0 | 1 | 100 | 0 | 3 | NA | NA | 0 | NA | NA | 0 |
| TOTAL PHYS ACT | MALE vs. FEMALE | 89 | 11 | 27 | NA | NA | 0 | NA | NA | 0 | NA | NA | 0 | 75 | 25 | 4 |
| TOTAL PHYS ACT | AGE | NA | NA | 0 | NA | NA | 0 | NA | NA | 0 | NA | NA | 0 | 0 | 100 | 3 |
| TOTAL PHYS ACT | BMI | 0 | 100 | 25 | NA | NA | 0 | NA | NA | 0 | NA | NA | 0 | 0 | 100 | 2 |
| TOTAL PHYS ACT | HOMA-IR | 0 | 100 | 14 | NA | NA | 0 | NA | NA | 0 | NA | NA | 0 | 0 | 100 | 4 |
| TOTAL PHYS ACT | T2D vs. NGT | NA | NA | 0 | NA | NA | 0 | NA | NA | 0 | NA | NA | 0 | NA | NA | 0 |
| TOTAL PHYS ACT | 2-HOUR GLUCOSE | NA | NA | 0 | NA | NA | 0 | NA | NA | 0 | NA | NA | 0 | NA | NA | 0 |
| VIGOROUS PHYS ACT | MALE vs. FEMALE | 82 | 18 | 121 | 74 | 26 | 61 | 86 | 14 | 7 | 93 | 7 | 28 | 95 | 5 | 175 |
| VIGOROUS PHYS ACT | AGE | 20 | 80 | 5 | 0 | 100 | 18 | 0 | 100 | 2 | NA | NA | 0 | 4 | 96 | 55 |
| VIGOROUS PHYS ACT | BMI | 1 | 99 | 127 | 6 | 94 | 18 | 0 | 100 | 5 | 0 | 100 | 18 | 8 | 92 | 40 |
| VIGOROUS PHYS ACT | HOMA-IR | 0 | 100 | 56 | 0 | 100 | 22 | 0 | 100 | 5 | 0 | 100 | 14 | 6 | 94 | 69 |
| VIGOROUS PHYS ACT | T2D vs. NGT | NA | NA | 0 | 0 | 100 | 1 | 0 | 100 | 1 | NA | NA | 0 | NA | NA | 0 |
| VIGOROUS PHYS ACT | 2-HOUR GLUCOSE | NA | NA | 0 | 0 | 100 | 3 | 0 | 100 | 1 | NA | NA | 0 | 33 | 67 | 3 |
| VIGOROUS PHYS ACT | TOTAL PHYS ACT | 100 | 0 | 62 | NA | NA | 0 | NA | NA | 0 | NA | NA | 0 | 100 | 0 | 8 |

**Table S9. Number of significant (FDR<5%) gene sets per trait across cell types.** Number of significant (FDR<5%) gene sets for a trait from models adjusting only for base covariates. Values specified by NA indicate that less than 25 samples were available with that trait in that cell type and no models were run for these trait-cell type combinations.

| # of significant (FDR<5%) gene sets for trait of interest | | | | | | | | | | | | |
|---|---|---|---|---|---|---|---|---|---|---|---|---|
| Trait of interest | Endothelial cell | Fibro-adipogenic progenitor cell | Macro-phage | Neuro-muscular junction (muscle fiber) | Neuronal cell | Satellite dell | Smooth muscle cell | Type 1 muscle fiber | Type 2a muscle fiber | Type 2x muscle fiber | Total pseudobulk | Bulk |
| AGE | 0 | 22 | 1 | 0 | 0 | 10 | 0 | 153 | 207 | 78 | 180 | 787 |
| BMI | 1 | 15 | 0 | 2 | 0 | 0 | 3 | 341 | 399 | 162 | 394 | 624 |
| 2-HOUR GLUCOSE | 0 | 0 | 0 | 3 | 0 | 0 | 0 | 113 | 260 | 269 | 357 | 495 |
| HOMA-IR | 0 | 0 | 0 | 17 | 0 | 0 | 0 | 270 | 619 | 317 | 163 | 346 |
| TOTAL PHYS ACT | 0 | 4 | 1 | 10 | 0 | 0 | 2 | 683 | 172 | 196 | 479 | 756 |
| VIGOROUS PHYS ACT | 0 | 0 | 0 | 31 | 0 | 0 | 0 | 468 | 239 | 223 | 446 | 889 |
| MALE vs. FEMALE | 0 | 0 | 0 | 0 | 0 | 0 | 0 | 178 | 108 | 73 | 410 | 1172 |
| T2D vs. NGT | 2 | 0 | NA | 6 | NA | 0 | 0 | 212 | 309 | 184 | 472 | 665 |

**Table S10 (separate file). Gene set-trait associations across cell types, total pseudobulk, and bulk tissue.** Concept ID, Concept name, Trait of interest, 2nd trait adjusted for, Cell type, Odds ratio, Coefficient, Status, P-value, FDR, Number of genes. NAs for the 2nd trait adjusted for indicates that no additional trait was adjusted for. Rows with an FDR<5% are significant. For detailed information on the models and how FDR was calculated please consult the method section.

**Table S11. Number of significant (FDR<5%) gene sets for both traits across muscle fiber types, total pseudobulk, and bulk tissue.** Total number (T) of significant gene sets for both trait 1 and trait 2 from models adjusting only for base covariates and percentage (%) of total gene sets that have a concordant (C) or discordant (D) direction of association of gene set-trait association for these two traits.

| # of significant (FDR<5%) gene sets for both trait 1 and trait 2 | | | | | | | | | | | | | | | | |
|---|---|---|---|---|---|---|---|---|---|---|---|---|---|---|---|---|
| Trait 1 | Trait 2 | Type 1 | | | Type 2a | | | Type 2x | | | Total pseudobulk | | | Bulk | | |
| | | C (%) | D (%) | T (#) | C (%) | D (%) | T (#) | C (%) | D (%) | T (#) | C (%) | D (%) | T (#) | C (%) | D (%) | T (#) |
| AGE | MALE vs. FEMALE | 0 | 100 | 4 | 5 | 95 | 21 | 0 | 100 | 6 | 2 | 98 | 54 | 1 | 99 | 391 |
| BMI | MALE vs. FEMALE | 2 | 98 | 40 | 0 | 100 | 9 | 50 | 50 | 2 | 0 | 100 | 110 | 13 | 87 | 310 |
| BMI | AGE | 93 | 7 | 14 | 97 | 3 | 30 | 75 | 25 | 4 | 65 | 35 | 17 | 68 | 32 | 292 |
| HOMA-IR | MALE vs. FEMALE | 0 | 100 | 27 | 31 | 69 | 16 | 100 | 0 | 7 | 0 | 100 | 59 | 23 | 77 | 173 |
| HOMA-IR | AGE | 100 | 0 | 15 | 93 | 7 | 56 | 100 | 0 | 18 | 81 | 19 | 16 | 90 | 10 | 150 |
| HOMA-IR | BMI | 100 | 0 | 173 | 100 | 0 | 294 | 100 | 0 | 119 | 100 | 0 | 92 | 100 | 0 | 205 |
| T2D vs. NGT | MALE vs. FEMALE | 0 | 100 | 17 | 0 | 100 | 14 | 100 | 0 | 7 | 2 | 98 | 115 | 58 | 42 | 235 |
| T2D vs. NGT | AGE | 100 | 0 | 29 | 100 | 0 | 73 | 100 | 0 | 18 | 100 | 0 | 78 | 90 | 10 | 178 |
| T2D vs. NGT | BMI | 100 | 0 | 70 | 100 | 0 | 95 | 100 | 0 | 38 | 100 | 0 | 79 | 28 | 72 | 121 |
| T2D vs. NGT | HOMA-IR | 100 | 0 | 79 | 100 | 0 | 176 | 100 | 0 | 110 | 100 | 0 | 90 | 97 | 3 | 97 |
| 2-HOUR GLUCOSE | MALE vs. FEMALE | 0 | 100 | 2 | 0 | 100 | 13 | 100 | 0 | 7 | 2 | 98 | 98 | 64 | 36 | 166 |
| 2-HOUR GLUCOSE | AGE | 100 | 0 | 17 | 100 | 0 | 55 | 100 | 0 | 21 | 100 | 0 | 51 | 90 | 10 | 124 |
| 2-HOUR GLUCOSE | BMI | 100 | 0 | 72 | 100 | 0 | 112 | 100 | 0 | 64 | 100 | 0 | 108 | 63 | 37 | 93 |
| 2-HOUR GLUCOSE | HOMA-IR | 100 | 0 | 72 | 100 | 0 | 166 | 100 | 0 | 153 | 100 | 0 | 99 | 100 | 0 | 120 |
| 2-HOUR GLUCOSE | T2D vs. NGT | 100 | 0 | 59 | 100 | 0 | 151 | 100 | 0 | 132 | 100 | 0 | 208 | 100 | 0 | 366 |
| TOTAL PHYS ACT | MALE vs. FEMALE | 100 | 0 | 59 | 100 | 0 | 12 | 100 | 0 | 2 | 100 | 0 | 139 | 100 | 0 | 303 |
| TOTAL PHYS ACT | AGE | 0 | 100 | 75 | 0 | 100 | 43 | 0 | 100 | 27 | 0 | 100 | 71 | 1 | 99 | 256 |
| TOTAL PHYS ACT | BMI | 0 | 100 | 164 | 0 | 100 | 74 | 0 | 100 | 26 | 0 | 100 | 135 | 72 | 28 | 123 |
| TOTAL PHYS ACT | HOMA-IR | 0 | 100 | 126 | 0 | 100 | 104 | 0 | 100 | 62 | 0 | 100 | 99 | 49 | 51 | 37 |
| TOTAL PHYS ACT | T2D vs. NGT | 0 | 100 | 102 | 0 | 100 | 69 | 0 | 100 | 47 | 0 | 100 | 168 | 26 | 74 | 227 |
| TOTAL PHYS ACT | 2-HOUR GLUCOSE | 0 | 100 | 66 | 0 | 100 | 59 | 0 | 100 | 57 | 1 | 99 | 136 | 33 | 67 | 161 |
| VIGOROUS PHYS ACT | MALE vs. FEMALE | 100 | 0 | 33 | 100 | 0 | 16 | 0 | 100 | 9 | 99 | 1 | 102 | 94 | 6 | 291 |
| VIGOROUS PHYS ACT | AGE | 0 | 100 | 66 | 0 | 100 | 74 | 0 | 100 | 24 | 0 | 100 | 77 | 3 | 97 | 331 |
| VIGOROUS PHYS ACT | BMI | 0 | 100 | 131 | 0 | 100 | 79 | 0 | 100 | 37 | 2 | 98 | 90 | 48 | 52 | 200 |
| VIGOROUS PHYS ACT | HOMA-IR | 0 | 100 | 112 | 0 | 100 | 110 | 0 | 100 | 98 | 0 | 100 | 78 | 6 | 94 | 131 |
| VIGOROUS PHYS ACT | T2D vs. NGT | 0 | 100 | 90 | 0 | 100 | 122 | 0 | 100 | 87 | 0 | 100 | 214 | 1 | 99 | 317 |
| VIGOROUS PHYS ACT | 2-HOUR GLUCOSE | 0 | 100 | 70 | 0 | 100 | 99 | 0 | 100 | 106 | 0 | 100 | 143 | 3 | 97 | 229 |
| VIGOROUS PHYS ACT | TOTAL PHYS ACT | 100 | 0 | 338 | 100 | 0 | 101 | 100 | 0 | 110 | 100 | 0 | 229 | 100 | 0 | 417 |

**Table S12. Number of significant (FDR<5%) peaks per trait across cell types.** Number of significant (FDR<5%) genes for a trait from models adjusting only for base covariates.

| Trait of interest | Adi-pocyte | | Endothe-lial cell | | Fibro-adipo-genic progenitor cell | | Macro-phage | | Neuro-muscular junction (muscle fiber) | | Neuronal cell | | Satellite cell | | Smooth muscle cell | | T cell | | Type 1 muscle fiber | | Type 2a muscle fiber | | Type 2x muscle fiber | |
|---|---|---|---|---|---|---|---|---|---|---|---|---|---|---|---|---|---|---|---|---|---|---|---|---|
| | # | % | # | % | # | % | # | % | # | % | # | % | # | % | # | % | # | % | # | % | # | % | # | % |
| AGE | 1 | 0.0 | 0 | 0.0 | 0 | 0.0 | 0 | 0.0 | 0 | 0.0 | 2 | 0.0 | 0 | 0.0 | 0 | 0.0 | 1 | 0.0 | 306 | 0.0 | 2307 | 0.5 | 41 | 0.0 |
| BMI | 0 | 0.0 | 0 | 0.0 | 0 | 0.0 | 0 | 0.0 | 0 | 0.0 | 0 | 0.0 | 0 | 0.0 | 0 | 0.0 | 0 | 0.0 | 3194 | 0.3 | 2514 | 0.5 | 1200 | 0.2 |
| 2-HOUR GLUCOSE | 0 | 0.0 | 0 | 0.0 | 0 | 0.0 | 0 | 0.0 | 0 | 0.0 | 0 | 0.0 | 0 | 0.0 | 0 | 0.0 | 0 | 0.0 | 1 | 0.0 | 3 | 0.0 | 3 | 0.0 |
| HOMA-IR | 0 | 0.0 | 0 | 0.0 | 0 | 0.0 | 1 | 0.0 | 0 | 0.0 | 1 | 0.0 | 0 | 0.0 | 0 | 0.0 | 0 | 0.0 | 1128 | 0.1 | 1882 | 0.4 | 471 | 0.1 |
| TOTAL PHYS ACT | 0 | 0.0 | 0 | 0.0 | 0 | 0.0 | 0 | 0.0 | 0 | 0.0 | 0 | 0.0 | 0 | 0.0 | 0 | 0.0 | 0 | 0.0 | 160 | 0.0 | 34 | 0.0 | 1 | 0.0 |
| VIGOROUS PHYS ACT | 1 | 0.0 | 0 | 0.0 | 0 | 0.0 | 0 | 0.0 | 0 | 0.0 | 0 | 0.0 | 0 | 0.0 | 0 | 0.0 | 0 | 0.0 | 1010 | 0.1 | 3005 | 0.6 | 318 | 0.0 |
| MALE vs. FEMALE | 26 | 0.1 | 376 | 0.1 | 9510 | 4.8 | 27 | 0.0 | 659 | 0.8 | 569 | 1.3 | 2497 | 3.0 | 139 | 0.1 | 421 | 0.4 | 51039 | 5.5 | 62206 | 12.7 | 36919 | 5.1 |
| T2D vs. NGT | 0 | 0.0 | 2 | 0.0 | 0 | 0.0 | 0 | 0.0 | 0 | 0.0 | 0 | 0.0 | 0 | 0.0 | 0 | 0.0 | 0 | 0.0 | 0 | 0.0 | 0 | 0.0 | 0 | 0.0 |

**Table S13 (separate file). Chromatin accessibility-trait associations across cell types.** Peak, mean peak count, Log2 Fold Change, Log2 Fold Change Standard error, Statistic, P-value, FDR, Cell type, Trait of interest, 2nd trait adjusted for, sample size. NAs for the 2nd trait adjusted for indicates that no additional trait was adjusted for. Rows with an FDR<5% are significant. For detailed information on the models and how FDR was calculated please consult the method section.

**Table S14. Number of samples passing QC metrics.** For each modality, only samples with at least 10 nuclei were included in the analysis for each cell type. The ATAC modality was used for chromatin accessibility analyses while the RNA modality was used for gene expression analyses. The mean and range of nuclei of each individual retained in the sample for each cell type is listed by modality.

| Cell Type | Modality | # of samples with ≥10 nuclei | Mean # of nuclei per retained individual | Range of nuclei per retained individual |
|---|---|---|---|---|
| Adipocyte | ATAC | 176 | 19.94 | [10,65] |
| | RNA | 4 | 15.25 | [10,18] |
| | RNA + ATAC | 181 | 21.08 | [10,83] |
| Endothelial cell | ATAC | 263 | 110.80 | [11,451] |
| | RNA | 253 | 35.23 | [10,184] |
| | RNA + ATAC | 263 | 144.95 | [15,635] |
| Macrophage | ATAC | 73 | 16.81 | [10,60] |
| | RNA | 29 | 13.76 | [10,22] |
| | RNA + ATAC | 128 | 19.92 | [10,76] |
| Fibro-adipogenic progenitor cell | ATAC | 255 | 48.89 | [10,422] |
| | RNA | 250 | 31.16 | [10,158] |
| | RNA + ATAC | 263 | 77.61 | [12,580] |
| Neuromuscular junction (muscle fiber) | ATAC | 137 | 24.15 | [10,199] |
| | RNA | 147 | 30.27 | [10,412] |
| | RNA + ATAC | 221 | 39.80 | [10,611] |
| Neuronal cell | ATAC | 236 | 37.33 | [10,156] |
| | RNA | 36 | 17.67 | [10,48] |
| | RNA + ATAC | 246 | 41.35 | [10,161] |
| Satellite cell | ATAC | 163 | 17.60 | [10,48] |
| | RNA | 122 | 14.78 | [10,35] |
| | RNA + ATAC | 231 | 25.19 | [10,79] |
| Smooth muscle cell | ATAC | 251 | 39.16 | [10,467] |
| | RNA | 218 | 25.30 | [10,306] |
| | RNA + ATAC | 262 | 60.00 | [10,773] |
| T cell | ATAC | 260 | 53.29 | [12,138] |
| | RNA | 2 | 12.50 | [12,13] |
| | RNA + ATAC | 260 | 54.21 | [12,140] |
| Type 1 muscle fiber | ATAC | 263 | 273.86 | [26,992] |
| | RNA | 263 | 257.97 | [24,907] |
| | RNA + ATAC | 263 | 531.83 | [59,1899] |
| Type 2a muscle fiber | ATAC | 261 | 147.13 | [19,428] |
| | RNA | 262 | 160.34 | [16,495] |
| | RNA + ATAC | 263 | 305.79 | [12,851] |
| Type 2x muscle fiber | ATAC | 262 | 149.23 | [10,556] |
| | RNA | 258 | 92.15 | [13,366] |
| | RNA + ATAC | 262 | 240.05 | [13,922] |
| Total pseudobulk | ATAC | 263 | 902.82 | [119,2317] |
| | RNA | 263 | 635.27 | [122,1665] |
| | RNA + ATAC | 263 | 1538.10 | [289,3822] |

**Table S15. Number of tested genes and peaks in gene expression and chromatin accessibility analyses respectively.** The same number of genes and peaks was tested for each trait.

| Cell Type | # of genes tested | # of peaks tested |
|---|---|---|
| Bulk | 22309 | Not tested |
| Total pseudobulk | 18618 | Not tested |
| Adipocyte | Not tested | 20706 |
| Endothelial cell | 14294 | 317823 |
| Macrophage | 11524 | 58936 |
| Fibro-adipogenic progenitor cell | 14826 | 196257 |
| Neuromuscular junction (muscle fiber) | 13400 | 78618 |
| Neuronal cell | 13933 | 45438 |
| Satellite cell | 11490 | 83481 |
| Smooth muscle cell | 13756 | 97305 |
| T cell | Not tested | 109228 |
| Type 1 muscle fiber | 23849 | 927588 |
| Type 2a muscle fiber | 19283 | 488001 |
| Type 2x muscle fiber | 17518 | 718348 |

**Table S16. Number of samples included for gene expression analyses for each trait by cell type.** Values specified by NA indicate that less than 25 samples were available with that trait in that cell type and no models were run for these trait-cell type combinations.

| Cell Type | MALE vs. FEMALE, AGE, BMI, HOMA-IR | 2-HOUR GLUCOSE, TOTAL PHYS ACT, VIGOROUS PHYS ACT | T2D vs. NGT |
|---|---|---|---|
| Bulk | 252 | 251 | 155 |
| Total pseudobulk | 263 | 262 | 164 |
| Endothelial cell | 253 | 252 | 158 |
| Macrophage | 29 | 29 | NA |
| Fibro-adipogenic progenitor cell | 250 | 249 | 154 |
| Neuromuscular junction (muscle fiber) | 147 | 147 | 84 |
| Neuronal cell | 36 | 36 | NA |
| Satellite cell | 122 | 121 | 73 |
| Smooth muscle cell | 218 | 217 | 138 |
| Type 1 muscle fiber | 263 | 262 | 164 |
| Type 2a muscle fiber | 262 | 261 | 163 |
| Type 2x muscle fiber | 258 | 257 | 162 |

**Table S17. Number of samples included for chromatin accessibility analyses for each trait by cell type.**

| Cell Type | MALE vs. FEMALE, AGE, BMI, HOMA-IR | 2-HOUR GLUCOSE, TOTAL PHYS ACT, VIGOROUS PHYS ACT | T2D vs. NGT |
|---|---|---|---|
| Adipocyte | 176 | 175 | 110 |
| Endothelial cell | 263 | 262 | 164 |
| Macrophage | 73 | 73 | 46 |
| Fibro-adipogenic progenitor cell | 255 | 254 | 160 |
| Neuromuscular junction (muscle fiber) | 137 | 136 | 86 |
| Neuronal cell | 236 | 235 | 147 |
| Satellite cell | 163 | 162 | 97 |
| Smooth muscle cell | 251 | 250 | 160 |
| T cell | 260 | 259 | 163 |
| Type 1 muscle fiber | 263 | 262 | 164 |
| Type 2a muscle fiber | 261 | 260 | 163 |
| Type 2x muscle fiber | 262 | 261 | 164 |
